# Single-dye, transfection-free FLIM multiplexing via bioorthogonal chemistry

**DOI:** 10.1101/2025.08.13.670148

**Authors:** Neville Dadina, Justin H. Kwon, Lauren Lesiak, Shuai Zheng, Madeline Zoltek, Daniel Brauer, Alanna Schepartz

**Author notes:** These authors contributed equally (Neville Dadina, Justin H. Kwon).

## Abstract

Fluorescence lifetime imaging microscopy (FLIM) can visualize multiple targets in a single spectral window, making it a powerful tool to overcome multiplexing limitations during live cell fluorescence microscopy. Here we show that small molecule probes–which are well-suited for imaging applications due to high specificity, low toxicity, and the elimination of transfection requirements–can be fine-tuned via bioorthogonal chemistry to exhibit predictably different fluorescent lifetimes suitable for FLIM multiplexing.

## Main

Fluorescence lifetime imaging microscopy (FLIM) is a powerful technique for multiplexed live cell imaging because it overcomes the limited working range of visible light. Multiplexed FLIM has thus far relied on multiple dyes or engineered transfected self-labeling proteins to tune fluorescence lifetime^1,2^. Here we show that three organelles or cell structures can be visualized selectively in live cells without transfection using a single dye and one bioorthogonal reaction–the tetrazine ligation–by exploiting the fact that different tetrazine reaction products possess the same emission wavelength but predictably different fluorescent lifetimes. This tool–which we refer to as Fluorescence Lifetime Imaging Microscopy INduced by bioorthoGOnal chemistry (FLIMINGO)–establishes standalone small molecule probes capable of imaging non-transfected cells on the basis of only fluorescence lifetime.

HD555 is a rhodamine fluorophore that is self-quenched by a pendant tetrazine^3^ (**Fig. 1a**). Inverse electron-demand Diels-Alder (IEDDA) reaction of the HD555 tetrazine^4,5^ eliminates self-quenching and results in a fluorescence increase (“turn-on”) whose magnitude varies with tetrazine reaction partner. Reaction of HD555 with bicyclo[6.1.0]non-4-yne (BCN)-containing partners provides greater turn-on than reaction with *trans*-cyclooctene (TCO)-containing partners^3,6^. As fluorescence turn-on and lifetime both correlate with quantum yield, we hypothesized that reaction partner-dependent turn-on could support multiplexed fluorescence lifetime imaging (FLIM) using a single dye. Indeed, reaction of HD555 *in vitro* with TCO, BCN, or spiro[2.3]hex-1-ene (SpH)^7^ partners generated products (**Fig. 1a, Supplementary Fig. 1**) with distinct amplitude-weighted fluorescence lifetimes (**Fig. 1b**). The longer lifetime of the BCN product is anticipated by the higher turn-on observed when HD555 reacts with derivatives of BCN^3,6^.

**Figure 1.**
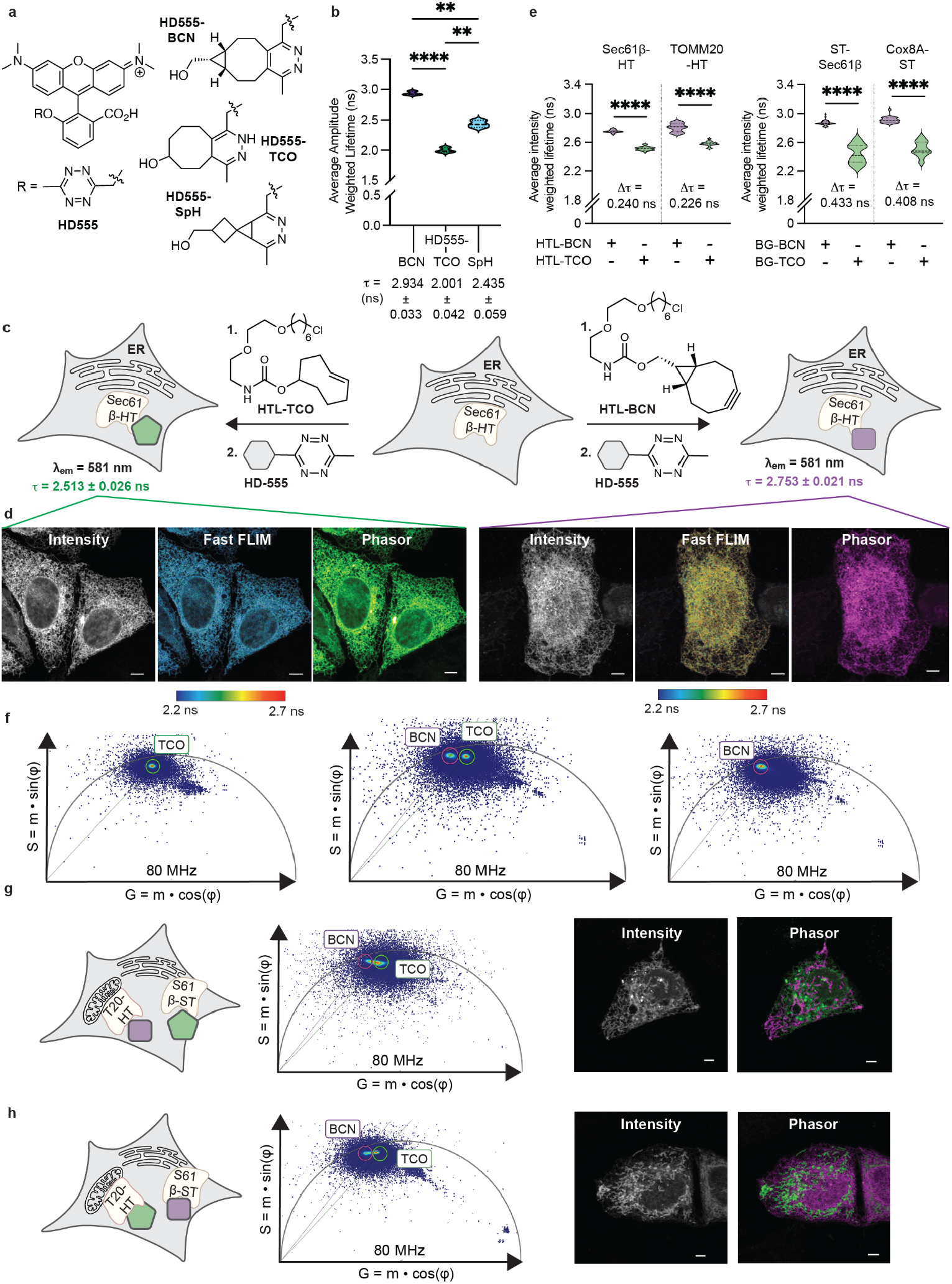
Fluorescence turn-on correlates with fluorescence lifetime. **a**, HD555 and the products of its reaction with BCN, TCO, and SpH-containing partners along with **b**, their average amplitude-weighted lifetimes determined *in vitro*. Adjusted P values for HD555-BCN vs. HD555-TCO (^****^P < 0.0001), HD555-BCN vs. HD555-SpH (^**^P = 0.0024), and HD555-TCO vs. HD555-SpH (^**^P = 0.0012) calculated using Dunnett’s T3 multiple comparisons test. Statistical significance was determined via Welch’s and Brown-Forsythe ANOVA with post hoc Dunnett’s T3 test accounting for heteroscedasticity. **c**, HeLa cells expressing Sec61β-HaloTag were treated with an HTL-TCO or HTL-BCN adapter followed by HD555 and imaged by detecting **d**, fluorescence intensity (Intensity), average photon arrival time (Fast FLIM, color-coded) or phasor analysis (Phasor). **e**, Average intensity-weighted lifetimes of cells expressing the indicated organelle marker and treated with an HTL-BCN or HTL-TCO adapter followed by HD555. ^****^P < 0.0001 for all pairwise comparisons between BCN and TCO with Sec61β-HT, TOMM20-HT, ST-Sec61β, or COX8A-ST: Sec61β-HT (Δ = 0.240 ns), TOMM20-HT (Δ = 0.226 ns), ST-Sec61β (Δ = 0.433 ns), and COX8A-ST (Δ = 0.408 ns). Adjusted P values calculated using Dunnett’s T3 multiple comparisons test. Statistical significance was determined via Welch’s and Brown-Forsythe ANOVA with post hoc Dunnett’s T3 test accounting for heteroscedasticity. **f**, Individual and overlayed phasor plots of FLIM data. **g**, Two-species FLIM imaging of HeLa cells expressing the indicated organelle markers (TOMM20-HaloTag and Sec61β-SNAPTag) and treated with HTL-BCN and BG-TCO or **h**, HTL-TCO and BG-BCN then HD555. Scale bars, 5 µm.

To test if reaction partner-dependent changes in fluorescence lifetime could be detected in cells, HeLa cells expressing Sec61β-HaloTag were first treated with an adapter, a HaloTag substrate carrying a terminal *trans*-cyclooctene (HTL-TCO) or a terminal bicyclo[6.1.0]nonyne (HTL-BCN) (**Fig. 1c**), to localize the two tetrazine partners to the cytoplasmic face of the endoplasmic reticulum (ER). Subsequent treatment with HD555 (30 min) yielded HeLa cell populations that were indistinguishable when visualized using fluorescence emission, but distinct when visualized by fluorescence lifetime. Lifetimes were determined on the basis of both average photon arrival time (Fast FLIM) and average intensity-weighted fluorescence lifetime (AIWFL). Regardless of method, the fluorescence lifetimes of cells treated with HTL-BCN were higher (2.483 ± 0.063 ns and 2.753 ± 0.021 ns, respectively) than cells treated with HTL-TCO (2.483 ± 0.063 ns and 2.513 ± 0.026 ns, respectively) (**Fig. 1d,e**). Similar differences were observed when analogous experiments were performed in three additional cellular locales (**Fig. 1e, Supplementary Fig. 2-4**). In all cases, the product resulting from reaction of HD555 with a BCN partner displayed a higher lifetime than the product with the otherwise analogous TCO partner (**Fig. 1e**).

Partner-dependent lifetime differences could also be distinguished using phasor analysis^8^ (**Fig. 1d, Supplementary Fig. 2-4)**, suggesting that they could support two-species imaging. Indeed, phasor analysis of cells expressing both Sec61β-SNAPTag and TOMM20-HaloTag, and treated with both BG-TCO and HTL-BCN followed by HD555, reveals two lifetime populations that effectively differentiate the ER and mitochondria (**Fig. 1g**). The ER and mitochondria were also differentiated when reactive handles were flipped (BG-BCN and HTL-TCO) (**Fig. 1h**).

These experiments provide evidence that two cell components labeled upon transfection can be visualized selectively using one bioorthogonal reaction and one spectral channel, by exploiting the fact that different tetrazine ligation reaction products possess predictably different fluorescent lifetimes. Next, we asked whether the same outcome could be achieved without transfection, using cell-permeant small molecules carrying different tetrazine reaction partners. TCO-tagged small molecules can localize selectively to discrete cellular locales^9–14^. We reasoned that if different organelle or structure-selective small molecules carried different tetrazine reaction partners, then we could exploit their reaction with HD555 for multiplexed FLIM.

To evaluate whether reaction partner-guided changes in fluorescence lifetime would differentiate organelles tagged by cell-permeant small molecules, we synthesized TCO and BCN conjugates of HAO, a lipid-like small molecule that localizes to the inner mitochondrial membrane (**Fig. 2a**)^14^. HeLa cells were treated–at first, separately–with HAO-BCN or HAO-TCO and then with HD555 (**Fig. 2b)**. Confocal microscopy confirmed the colocalization of fluorescence signals due to HD555 with those of HAO and MitoTracker Deep Red but not with the nuclear marker Hoechst 33342, as expected (**Supplementary Fig. 5**).

**Figure 2.**
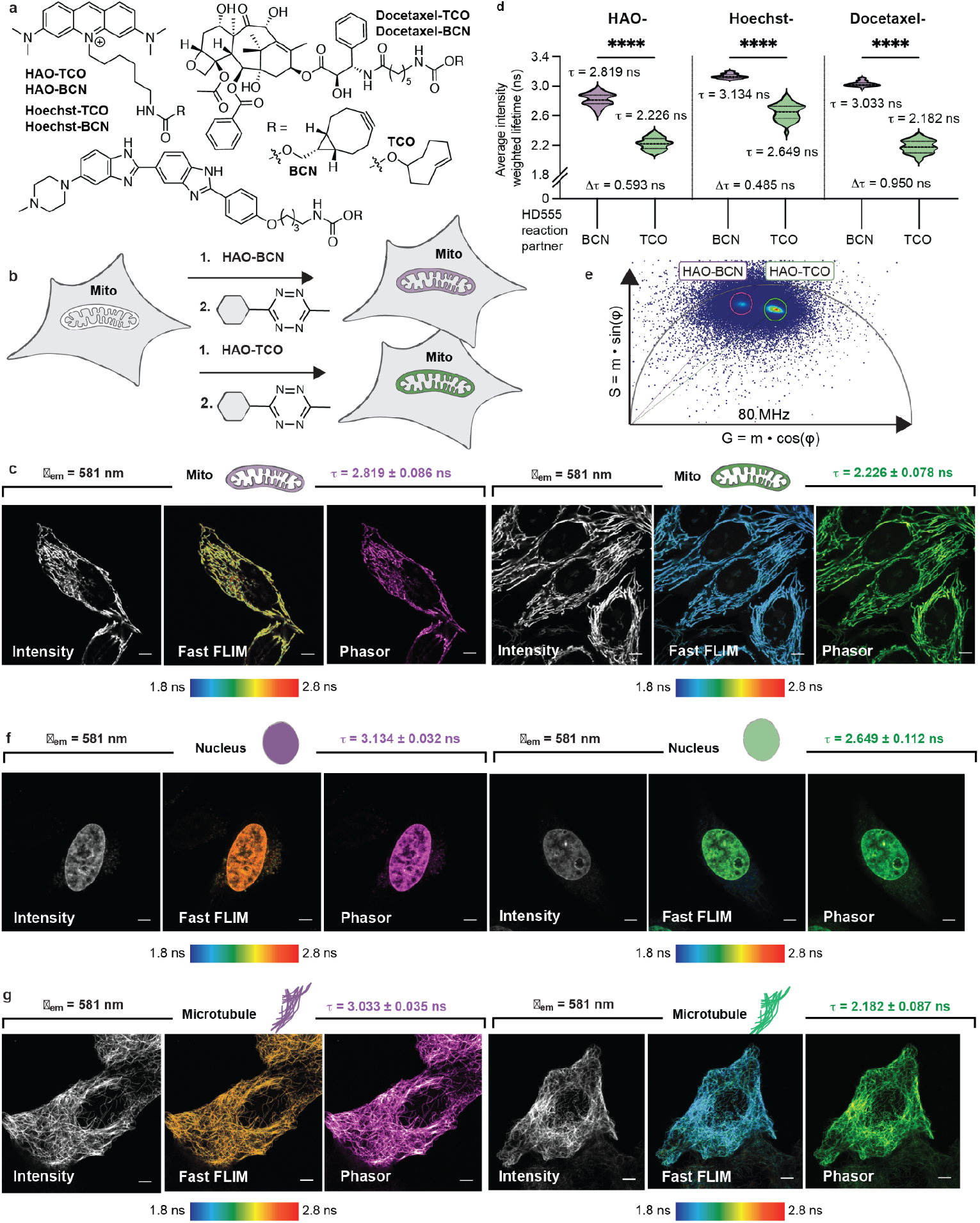
Single-species FLIM using multiple organelle- or structure-selective small molecules and one fluorophore. **a**, TCO and BCN derivatives of HAO, Hoechst, and Docetaxel, which localize to the inner mitochondrial membrane (IMM), nucleus, or microtubule network, respectively. **b**, HeLa cells were treated with HAO-BCN or HAO-TCO and **c**, imaged by detecting fluorescence intensity (Intensity), average photon arrival time (Fast FLIM), or phasor analysis (Phasor). **d**, Plot of average intensity-weighted fluorescence lifetimes associated with cells treated with the indicated small molecules. ^****^P < 0.0001 for all pairwise comparisons between BCN and TCO small molecules-derivatives: HAO (Δ = 0.5930 ns), Hoechst (Δ = 0.4853 ns), and Docetaxel (Δ = 0.8504 ns). Adjusted P values calculated using Dunnett’s T3 multiple comparisons test. Statistical significance was determined via Welch’s and Brown-Forsythe ANOVA with post hoc Dunnett’s T3 test accounting for heteroscedasticity. **e**, Composite phasor plot of FLIM data acquired from cells treated with HAO-BCN and HAO-TCO. **f**,**g**, HeLa cells were treated with **f**, Hoechst-BCN or Hoechst-TCO or **g**, Docetaxel-BCN or Docetaxel-TCO and imaged by detecting fluorescence intensity (intensity), average photon arrival time (Fast FLIM), or phasor analysis (Phasor). Scale bars, 5 µm.

Again, cells treated with HAO-BCN or HAO-TCO followed by HD555 appeared identical when visualized on the basis of fluorescent intensity but not fluorescence lifetime (**Fig. 2c**). HAO-BCN-treated cells displayed a longer fluorescent lifetime, determined by either Fast FLIM or AIWFL (2.435 ± 0.054 ns and 2.819 ± 0.086 ns, respectively) than HAO-TCO-treated cells (1.993 ± 0.070 ns and 2.226 ± 0.078 ns, respectively) (**Fig 2c,d**) and the signals due to the two HD555 products were easily distinguished using phasor analysis^8^ (**Fig. 2c,e)**. Analogous reaction product-dependent differences in fluorescence lifetime were also seen outside the mitochondria, in cells treated with the BCN and TCO derivatives of Docetaxel (to image microtubules) or Hoechst (to image the nucleus) (**Fig. 2d,f,g** and **Supplementary Fig. 6**).

When cells were treated with both Hoechst-BCN and HAO-TCO, the signals could be separated using either Fast FLIM or phasor analysis to clearly differentiate the nucleus and mitochondria (**Fig. 3a**). The same was true when the cells were treated with HAO-BCN and Hoechst-TCO to differentiate the mitochondria and nucleus or with Hoechst-BCN and Docetaxel-TCO to differentiate the nucleus and microtubules (**Fig. 3c**). Thus, in three separate cases, two different cell components could be visualized selectively without transfection using only one bioorthogonal reaction, the tetrazine ligation, and one dye, by exploiting the fact that different tetrazine ligation products possess predictably different fluorescent lifetimes.

**Fig. 3.**
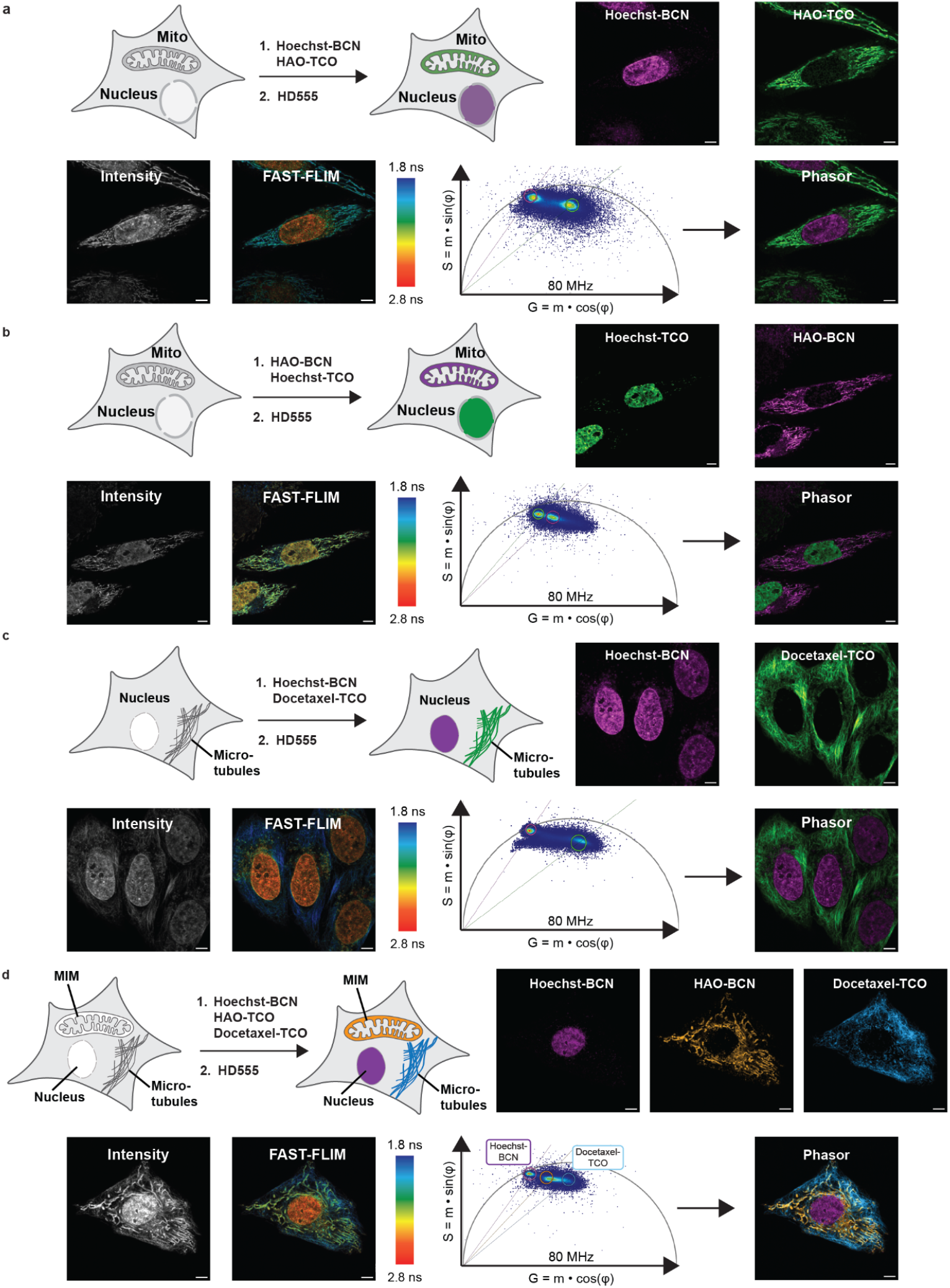
Two- and three-species FLIM multiplexing in a live cell using a single dye and only cell-permeant localization probes. HeLa cells were incubated with **a**, Hoechst-BCN and HAO-TCO; **b**, HAO-BCN and Hoechst-TCO; and **c**, Hoechst-BCN and Docetaxel-TCO followed by HD555. Cells were imaged by detecting total fluorescence intensity (Intensity) or average photon arrival time (Fast FLIM, color-coded) or using Phasor analysis to generate the single-species and dual-species images shown. **d**, HeLa cells were incubated with Hoechst-BCN, HAO-BCN, and Docetaxel-TCO. Scale bars, 5 µm.

Finally, we asked whether fluorescence lifetime differences could establish a three-species image in non-transfected cells using a single fluorophore. Although *in vitro* reaction of HD555 with a spiro[2.3]hex-1-ene (SpH) partner generated a product with a distinct lifetime (**Fig. 1b**), in our hands the SpH moiety was unstable. We speculated that we could leverage the especially long fluorescence lifetimes of HD555 reaction products in the nucleus to differentiate the nucleus and mitochondria with molecules equipped with the identical tetrazine reaction partner (**Fig. 2d**). Doing so would demand that phasor analysis could differentiate signals generated from cells treated with HAO-BCN and Hoechst-BCN followed by HD555. Indeed, the fluorescence lifetimes of the two HD555 reaction products–HAO-BCN-HD555 and Hoechst-BCN-HD555–provided excellent contrast between the nucleus and mitochondria, regardless of whether the lifetimes were determined by Fast FLIM or Phasor analysis (**Supplementary Fig. 7**). We next incubated cells with Hoechst-BCN, HAO-BCN, and Docetaxel-TCO, followed by HD555 (**Fig. 3d**). When visualized by Fast FLIM, we observed a distinct set of lifetime populations for each of the three cellular targets. Separation of the signals using phasor analysis resulted in three individual images, one for each of the three cellular targets (**Fig. 3d**). Thus three cell components can be visualized selectively without transfection using one bioorthogonal reaction–the tetrazine ligation–and one spectral channel, by exploiting the fact that different tetrazine ligation reaction products are associated with predictably different fluorescent lifetimes.

FLIMINGO was conceived on the basis of a predicted correlation between fluorescence turn-on and fluorescence lifetime. For HD555, fluorescent turn-on is large (**Supplementary Fig. 1**)^3^. To test whether a dye with a lower turn-on could also generate TCO and BCN products with distinct fluorescent lifetimes, we turned to TAMRA-tetrazine, a widely used, modestly quenched fluorophore. Reaction of TAMRA-tetrazine *in vitro* with BCN-, TCO-, and SpH-containing partners resulted in turn-on values an order of magnitude lower than those observed with HD555 (**Supplementary Fig. 8**). Regardless, the fluorescence lifetimes of the products were clearly separable and their relative values followed the expected trend TAMRA-SpH ∼ TAMRA-BCN > TAMRA-TCO (**Supplementary Fig. 9**). Thus, we expect that a range of modestly quenched, click-reactive dyes may be FLIMINGO compatible.

Fluorescence microscopy is a foundational tool in the cell biology arsenal. Multi-target fluorescence imaging remains limited by a persistent hurdle— spectral overlap. Fluorescence lifetime imaging microscopy (FLIM) offers a solution to this limitation. The method described herein allows FLIM to be applied in non-transfected cells using only small molecule reagents. By modulating the tetrazine reaction partner, a single cell-compatible bioorthogonal reaction can establish a set of intracellular fluorophores possessing distinct but predictable fluorescent lifetimes. This feature enables multiplexing within a single spectral window.

## Acknowledgements

This work was supported by the NIH (Grant No. 1R35GM134963 to A.S.). Instruments in the CoC-NMR are supported in part by NIH S10OD024998.

## Author contributions

N.D. conceived the project; N.D., J.K., and A.S. developed the project and designed experiments. N.D., L.L., S.Z., J.K., synthesized molecules for experiments. M.Z. generated genetic materials for experiments. N.D. and J.K. analyzed the data with inputs from A.S. N.D., J.K., D.B., and A.S. wrote the manuscript.

## Methods

### Materials

Materials for chemical synthesis were purchased from commercial vendors. Culturing medium DMEM (12430054, 21063029 and A1443001), Opti-MEM with no phenol red (11058021) and FBS were all purchased from Thermo Fisher. The plasmid encoding Halo-TOMM20 was a gift from K. McGowan (HHMI Janelia, Halo-TOMM20-N-10, Addgene plasmid 123284); that encoding COX8A-SNAP was a gift from A. Egana (New England Biolabs, pSNAPf-COX8A, Addgene plasmid 101129); that encoding Halo-Sec61-C-18 was a gift from Kevin McGowan (Addgene plasmid # 123285; RRID:Addgene_123285); that encoding SNAP-Sec61β was a gift from Gia Voeltz (Addgene plasmid # 141152; RRID:Addgene_141152).

#### General cell culture notes

HeLa cells (University of California, Berkeley, Cell Culture Facility) were cultured in DMEM (Thermo Fisher) supplemented with 10% FBS, penicillin (100 U mL^−1^) and streptomycin (100 μg mL^−1^). Cells were cultured at 37 °C in a humidified CO_2_/air (5%/95%) incubator. HeLa cells obtained from the University of California, Berkeley, Cell Culture Facility were periodically tested for *Mycoplasma* by using DNA methods. Cells for imaging were seeded in four-well glass-bottom μslides (Ibidi®, 80427, number #1.5H, 2.5 cm^2^) at the indicated densities.

#### Microscope specifications

All imaging experiments were performed on a Leica STELLARIS 8 microscope (Leica Microsystems) equipped with a Leica DMi8 CS scanhead, an HC Plan-Apo ×63/1.4-NA water immersion objective, and a pulsed white-light laser (440–790 nm; 440 nm: >1.1 mW; 488 nm: >1.6 mW; 560 nm: >2.0 mW; 630 nm: >2.6 mW; 790 nm: >3.5 mW, 78 MHz). Fluorescence Lifetime Imaging was performed using HyD X detectors in photon counting mode. Live-cell imaging conditions were maintained using a blacked-out cage enclosure from Okolab (748-4206). The temperature was maintained by heating the enclosure and was monitored with the Oko-Touch temperature display. The pH was maintained by supplying humidified 5% CO_2_ to the sample chamber.

#### Synthesis and characterization

Experimental procedures for the synthesis and characterization of HTL, HTL, BG, HAO, Hoechst, and Docetaxel derivatives (and intermediates), and their respective characterization data are included as Supplementary Information.

#### *In vitro* determination of fluorescence lifetimes

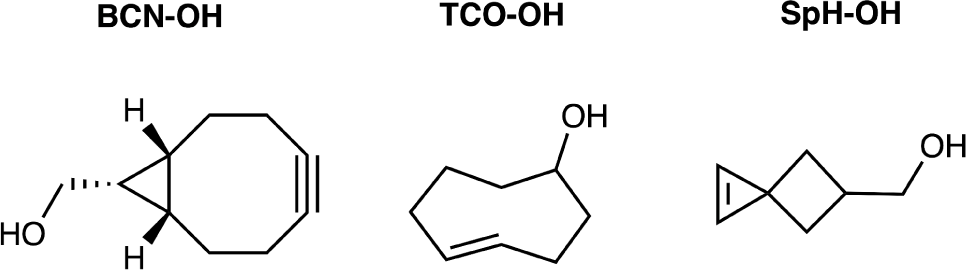

100 µM solutions of the tetrazine reaction partners BCN-OH (Sigma-Aldrich, ≥80%), TCO-OH (BroadPharm, 90%), and SpH-OH (Enamine, 95%) in Milli-Q water were prepared by diluting a 10 mM DMSO stock solution of each compound. A 10 µM solution of HD555 in Milli-Q water was prepared by diluting a 2 mM DMSO stock solution. 250 µL of the solution containing each tetrazine reaction partner was combined with 250 µL of the HD555 solution to generate a set of three final reaction mixtures containing 5 µM HD555 and 50 µM of a tetrazine reaction partner. Each mixture was incubated at 37°C for 2 h, then diluted 10-fold in Milli-Q water before acquiring lifetime measurements. All lifetime measurements were acquired via time-correlated single photon counting (TCSPC) using a FluoroMax Plus (HORIBA) fluorimeter and 1 cm quartz cuvettes (Hellma®). Samples were excited by a portable DeltaDiode laser (510 nm, HORIBA), and fluorescence was measured at an emission of 575 nm. Amplitude-weighted lifetimes were determined by fitting the curves to mono- or bi-exponential decay models. The number of exponents was determined by the best fit () with the least degrees of freedom.

#### Transfection and labelling of HeLa cells for single-species imaging

Forty-eight hours before imaging, HeLa cells were plated (1.5 × 10^4^ cells per well) in a four-well chamber and incubated overnight. After 16 h, the cells were transfected with plasmids encoding fusions of organelle-specific proteins with either HaloTag or SNAPTag (Halo-TOMM20-N-10, pSNAPf-COX8A, Halo-Sec61-C-18, or SNAP-Sec61β, 500 ng/well) using FuGENE HD (1.5 µL/well) transfection reagent (Promega), according to the manufacturer’s protocol, incubated for 6 hours at 37 °C, and washed three times with DMEM ph(−) supplemented with 10% FBS. The following day, the cells were labeled with 700 μL of either 10 µM of HTL-BCN or 10 µM HTL-TCO in Opti-MEM ph(–) for 30 min, washed three times with 700 μL of DMEM ph(–), incubated with 700 μL of 1 µM HD555 in Opti-MEM ph(–) for 0.5 h, washed three times with DMEM ph(–) supplemented with 10% FBS, and imaged.

#### Transfection and labelling of HeLa cells for two-species imaging using Halo-TOMM20+HTL-BCN and SNAP-Sec61β+BG-TCO

Forty-eight hours before imaging, HeLa cells were plated (1.5 × 10^4^ cells per well) in a four-well chamber and incubated overnight. After 16 h, the cells were transfected with two separate plasmids encoding Halo-TOMM20-N-10 (375 ng/well) and SNAP-Sec61β (125 ng/well) using FuGENE HD (1.5 µL/well) (Promega), according to manufacturer’s protocol, incubated for 6 h, and washed three times with DMEM ph(−) supplemented with 10% FBS. The following day, the cells were labeled with 700 μL of 4 µM HTL-BCN and 1 µM BG-TCO in Opti-MEM ph(–) for 30 min, washed three times with 700 μL of DMEM ph(–), incubated with 700 μL of 4 µM HD555 in Opti-MEM ph(–) for 30 min, washed three times with DMEM ph(–) supplemented with 10% FBS, and imaged.

#### Transfection and Labelling of HeLa cells for two-species imaging using Halo-TOMM20+HTL-TCO and SNAP-Sec61β+BG-BCN

Forty-eight hours before imaging, HeLa cells were plated (1.5 × 10^4^ cells per well) in a four-well chamber and incubated overnight. After 16 h, the cells were transfected with two separate plasmids encoding Halo-TOMM20-N-10 (250 ng/well) and SNAP-Sec61β (500 ng/well) using FuGENE HD (1.5 µL/well) transfection reagent (Promega), according to manufacturer’s protocol. The plasmids were added simultaneously to the well, incubated for 6 hrs, and washed three times with DMEM ph(−) supplemented with 10% FBS. The following day, the cells were labeled with 700 μL of 1 µM HTL-TCO and 4 µM BG-BCN in Opti-MEM ph(–) for 30 min, washed three times with 700 μL of DMEM ph(–), incubated with 700 μL of 4 µM HD555 in Opti-MEM ph(–) for 30 min, washed three times with DMEM ph(–) supplemented with 10% FBS, and imaged.

#### Labelling of HeLa cells for single-species imaging using HAO-BCN and HAO-TCO

Twenty-four hours before imaging, HeLa cells were plated (2.5 × 10^4^ cells per well) in a four-well chamber and incubated overnight. After 16 h, the cells were labeled with 700 μL of 100 nM HAO-BCN or 100 nM HAO-TCO in Opti-MEM ph(–) for 1 h. Cells treated with HAO-BCN were washed three times with 700 μL of DMEM ph(–), incubated with 700 μL of 200 nM HD555 in Opti-MEM ph(–) for 30 min, washed three times with DMEM ph(–) supplemented with 10% FBS, then rested for 1 h before imaging. Cells treated with HAO-TCO were washed three times with 700 μL of DMEM ph(–) supplemented with 10% FBS and allowed to rest for 1 h. Cells were then incubated with 700 μL of 200 nM HD555 in Opti-MEM ph(–) for 0.5 h, washed three times with DMEM ph(–) supplemented with 10% FBS, and imaged.

#### Labelling of HeLa cells for single-species imaging using Hoechst-BCN and Hoechst-TCO

Twenty-four hours before imaging, HeLa cells were plated (2.5 × 10^4^ cells per well) in a four-well chamber and incubated overnight. After 16 h, the cells were labeled with 700 μL of 200 nM Hoechst-BCN or 200 nM Hoechst-TCO in Opti-MEM ph(–) for 0.5 h, washed three times with 700 μL of DMEM ph(–), incubated with 700 μL of 200 nM HD555 in Opti-MEM ph(–) for 0.5 h, washed three times with DMEM ph(–) supplemented with 10% FBS, and imaged.

#### Labelling of HeLa cells for single-species imaging using Docetaxel-BCN and Docetaxel-TCO

Twenty-four hours before imaging, HeLa cells were plated (2.5 × 10^4^ cells per well) in a four-well chamber and incubated overnight. After 16 h, the cells were labeled with 700 μL of 100 nM Docetaxel-BCN or 100 nM Docetaxel-TCO in Opti-MEM ph(–) for 1 h, washed three times with 700 μL of DMEM ph(–), incubated with 700 μL of 500 nM HD555 in Opti-MEM ph(–) for 0.5 h, washed three times with DMEM ph(–) supplemented with 10% FBS, and imaged.

#### Labelling of HeLa cells for two-species imaging using HAO-TCO and Hoechst-BCN

Twenty-four hours before imaging, HeLa cells were plated (2.5 × 10^4^ cells per well) in a four-well chamber and incubated overnight. After 16 h, the cells were labeled with 700 μL of 100 nM HAO-TCO in Opti-MEM ph(–) for 1 h, washed three times with 700 μL of DMEM ph(–) supplemented with 10% FBS, then allowed to rest for 0.5 h. Cells were subsequently incubated with 700 μL of 200 nM Hoechst-BCN in Opti-MEM ph(–) for 0.5 h, washed three times with 700 μL of DMEM ph(–), then incubated with 700 μL of 400 nM HD555 in Opti-MEM ph(–) for 0.5 h, washed three times with DMEM ph(–) supplemented with 10% FBS, and imaged.

#### Labelling of HeLa cells for two-species imaging using HAO-BCN and Hoechst-BCN

Twenty-four hours before imaging, HeLa cells were plated (2.5 × 10^4^ cells per well) in a four-well chamber and incubated overnight. After 16 h, the cells were labeled with 700 μL of 100 nM HAO-BCN in Opti-MEM ph(–) for 1 h, washed three times with 700 μL of DMEM ph(–), incubated with 700 μL of 400 nM HD555 in Opti-MEM ph(–) for 0.5 h, then washed three times with DMEM ph(–). Cells were subsequently incubated with 700 μL of 200 nM Hoechst-BCN in Opti-MEM ph(–) for 0.5 h, washed three times with 700 μL of DMEM ph(–), incubated with 700 μL of 400 nM HD555 in Opti-MEM ph(–) for 0.5 h, then washed three times with DMEM ph(–) supplemented with 10% FBS, and imaged.

#### Labelling of HeLa cells for three-species imaging using Hoechst-BCN, HAO-BCN and Docetaxel-TCO

Twenty-four hours before imaging, HeLa cells were plated (4.0 × 10^4^ cells per well) in a two-well chamber and incubated overnight. After 16 h, the cells were labeled with 700 μL of 100 nM HAO-BCN in Opti-MEM ph(–) for 1 h, washed three times with 700 μL of DMEM ph(–), incubated with 700 μL of 1 µM HD555 in Opti-MEM ph(–) for 0.5 h, then washed three times with DMEM ph(–). Cells were then labeled with 700 μL of 200 nM Docetaxel-TCO in Opti-MEM ph(–) for 0.5 h and washed three times with 700 μL of DMEM ph(–). These cells were subsequently incubated with a 700 µL mixture containing both 200 nM Docetaxel-TCO and 100 nM Hoechst-BCN for 0.5 h, washed three times with 700 μL of DMEM ph(–), incubated with 700 μL of 500 nM HD555 in Opti-MEM ph(–) for 0.5 h, then washed three times with DMEM ph(–) supplemented with 10% FBS, and imaged.

#### Labelling of HeLa cells for colocalization between HAO-BCN/TCO, HD555, MTDR, and Hoechst 33342

Twenty-four hours before imaging, HeLa cells were plated (2.5 × 10^4^ cells per well) in a four-well chamber and incubated overnight. After 16 h, the cells were labeled with 700 μL of 100 nM HAO-BCN or HAO-TCO in Opti-MEM ph(–) for 1 h, washed three times with 700 μL of DMEM ph(–), incubated with 700 μL of 400 nM HD555 in Opti-MEM ph(–) for 0.5 h, then washed three times with DMEM ph(–). Cells were subsequently incubated with 700 μL of 50 nM Mito Tracker Deep Red in Opti-MEM ph(–) for 0.5 h, washed three times with 700 μL of DMEM ph(–), incubated with 700 μL of 300 nM Hoechst 33342 in Opti-MEM ph(–) for 5 min, then washed three times with DMEM ph(–) supplemented with 10% FBS, and imaged.

#### General fluorescence lifetime image acquisition settings

All experiments were performed on a Leica Stellaris 8 (as detailed above) equipped with FALCON for fluorescence lifetime imaging microscopy (FLIM). All images were acquired using the same beam path and scanhead settings. Given the fact that only one fluorophore was imaged across all experiments, settings remained consistent. The HC Plan-Apo ×63/1.4-NA water immersion objective was used for all experiments, and the correction collar was adjusted to correct for differences in coverslip thickness. **Beam path setting:** The excitation wavelength was set to 561 nm and the laser power (1-5%) was adjusted to collect between 0.3 and 0.5 photons per laser pulse. The HyD X detector window was set between 575 nm – 750 nm and the detector was set to photon counting mode. A 561 nm filter was engaged to filter out stray excitation photons. **Scanhead settings:** The pixel size was kept constant over all images at 60.06 × 60.06 nm. The field of view was also kept constant at 61.5 × 61.5 µm. The pixel dwell time was set to 3.8375 µs (scanning frequency of 200 lines per second). Line accumulation (between 8-12) was adjusted to ensure that >150 photons per pixel (>200 for multispecies) were collected per image. **IRF:** The instrument response function (IRF) for each data set (i.e. image) was automatically calculated by the LAS X FLIM-FCS Module.

#### Fluorescence lifetime determination of single-species images

All data were processed and worked up using the LAS X and LAS X FLIM-FCS Module (Leica Microsystems). Initially, images were screened to ensure that photons per pixel was >150. Then a brightness (photon count) threshold was set to exclude background photons from media. Next, a region of interest (ROI) was drawn around the cell/organelle to ensure that lifetime data was only coming from pixels belonging to the respective organelles. The decay histogram pertaining to an ROI was fit using the n-exponential reconvolution fitting model (Leica Microsystems—detailed equations included in the **Supplemental Information**). Intensity-weighted and amplitude-weighted lifetimes were determined by fitting the curves to bi- or tri-exponential decay models. The number of exponents was determined by the best fit () with the least degrees of freedom. To determine the average intensity-weighted lifetime of a population of cells, the intensity-weighted lifetime of >20 ROIs was averaged. To determine the average amplitude-weighted lifetime of a population of cells, the amplitude-weighted lifetime of >20 ROIs was averaged.

To determine phasor populations, single-species images were processed in the LAS X FLIM-FCS modules. First, images were thresholded to reduce background photons. Median filtering (filter strength = 9) was applied, and the cursor (radius = 15) was placed over the cluster that encapsulated the pixels from the ROI in the image.

#### Curve fitting for fast FLIM histograms for HAO-TCO and Hoechst-BCN images

Histograms of average photon arrival times (Fast FLIM histograms) were exported from LAS X software and plotted using GraphPad Prism. Data points representing lifetime distributions were then fit to Gaussian or Lorentzian functions (R^2^ > 0.95).

#### Phasor separation of multi-species images

FLIM data from multispecies images was processed and plotted on a phasor plot using the LAS X FLIM-FCS software package. First, images were thresholded to remove and background photons, and a median filter was applied (strength = 9). Cursor position were determined using the lifetimes and positions from the homogenous, single species images. Once the cursors were positioned for each component, the lifetime populations were separated into individual images.

#### Software and image processing

Images were acquired using Leica LAS X and curves were fitted using the LAS X FLIM-FCS Module. Fast FLIM Images were analyzed and exported using the LAS X software package. Statistical analysis and plotting were performed using GraphPad Prism 10.4.1. All images were processed with^15,16^.

## References

1. Frei, M. S. et al. Engineered HaloTag variants for fluorescence lifetime multiplexing. Nat. Methods 19, 65–70 (2022).

2. Frei, M. S., Koch, B., Hiblot, J. & Johnsson, K. Live-Cell Fluorescence Lifetime Multiplexing Using Synthetic Fluorescent Probes. ACS Chem. Biol. 17, 1321–1327 (2022).

3. Werther, P. et al. Bio-orthogonal Red and Far-Red Fluorogenic Probes for Wash-Free Live-Cell and Super-resolution Microscopy. ACS Cent. Sci. 7, 1561–1571 (2021).

4. Blackman, M. L., Royzen, M. & Fox, J. M. Tetrazine Ligation: Fast Bioconjugation Based on Inverse-Electron-Demand Diels−Alder Reactivity. J. Am. Chem. Soc. 130, 13518–13519 (2008).

5. Devaraj, N., Weissleder, R. & Hilderbrand, S. Tetrazine-Based Cycloadditions: Application to Pretargeted Live Cell Imaging. Bioconjug. Chem. 19, 2297–2299 (2008).

6. Hild, F., Werther, P., Yserentant, K., Wombacher, R. & Herten, D.-P. A dark intermediate in the fluorogenic reaction between tetrazine fluorophores and trans-cyclooctene. Biophys. Rep. 2, 100084 (2022).

7. Yu, Z. & Lin, Q. Design of Spiro[2.3]hex-1-ene, a Genetically Encodable Double-Strained Alkene for Superfast Photoclick Chemistry. J. Am. Chem. Soc. 136, 4153–4156 (2014).

8. Digman, M. A., Caiolfa, V. R., Zamai, M. & Gratton, E. The Phasor Approach to Fluorescence Lifetime Imaging Analysis. Biophys. J. 94, L14–L16 (2008).

9. Erdmann, R. S. et al. Super-Resolution Imaging of the Golgi in Live Cells with a Bioorthogonal Ceramide Probe. Angew. Chem.-Int. Ed. 53, 10242–10246 (2014).

10. Takakura, H. et al. Long time-lapse nanoscopy with spontaneously blinking membrane probes. Nat. Biotechnol. 35, 773–780 (2017).

11. Gupta, A., Rivera-Molina, F., Xi, Z., Toomre, D. & Schepartz, A. Endosome motility defects revealed at super-resolution in live cells using HIDE probes. Nat. Chem. Biol. 16, 408–+ (2020).

12. Thompson, A. D., Bewersdorf, J., Toomre, D. & Schepartz, A. HIDE Probes: A New Toolkit for Visualizing Organelle Dynamics, Longer and at Super-Resolution. Biochemistry 56, 5194–5201 (2017).

13. Thompson, A. D. et al. Long-Term Live-Cell STED Nanoscopy of Primary and Cultured Cells with the Plasma Membrane HIDE Probe DiI-SiR. Angew. Chem. Int. Ed. 56, 10408–10412 (2017).

14. Zheng, S. et al. Long-term super-resolution inner mitochondrial membrane imaging with a lipid probe. Nat. Chem. Biol. 20, 83–92 (2024).

15. Rueden, C. T. et al. ImageJ2: ImageJ for the next generation of scientific image data. BMC Bioinformatics 18, (2017).

16. Schindelin, J. et al. Fiji: an open-source platform for biological-image analysis. Nat. Methods 9, 676–682 (2012).

