## Supplementary Information for "Single-dye, transfection-free FLIM multiplexing via bioorthogonal chemistry"

### Supplemental Information

### Supplemental Notes

#### Supplemental Note 1: Decay-curve fitting and determination of intensity and amplitude weighted lifetimes

Fluorescence lifetime decay histograms were fit according to the following equations.

$$y(t) = \{IRF(t + Shift_{IRF}) + Bkgr_{IRF}\} \otimes \left\{ \sum_{i=0}^{n-1} \quad A[i]e^{-\left(\frac{t}{\tau[i]}\right)} + Bkgr \right\}$$

$$I_{Sum} = \sum_{k=0}^{n-1} \quad I[k]$$

$$A_{Sum} = \sum_{k=0}^{n-1} \quad A[k]$$

$$\tau_{Av\ Int} = \frac{\sum_{k=0}^{n-1} \quad I[k]\,\tau[k]}{I_{Sum}}$$

$$\tau_{Av\ Amp} = \frac{\sum_{k=0}^{n-1} \quad A[k]\,\tau[k]}{A_{Sum}}$$

Where:

| | |
|---|---|
| $n$ | Number of exponential components |
| $A$ | Amplitudes – Exponential pre-factors |
| $\tau$ | Exponential Decay Times (e.g. lifetimes) |
| $Bkgr$ | Tail Offset - Correction for background (afterpulsing, dark counts, environmental light) |
| $Shift_{IRF}$ | IRF Shift - Correction for IRF displacement (e.g. wavelength-dependent zero time of the detector) |
| $Bkgr_{IRF}$ | IRF Background - Correction for IRF background |
| $I$ | Intensities associated with each exponential component, normalized to the time resolution of the measured decay curve to be displayed in photon counts |
| $I_{Sum}$ | Sum of fluorescence Intensity for all components |
| $A_{Sum}$ | Sum of fluorescence Intensity for all components at time zero |
| $\tau_{Av\ Int}$ | Mean Photon Arrival Time - Intensity-weighted average lifetime |
| $\tau_{Av\ Amp}$ | Mean Decay Time - Amplitude-weighted average lifetime |

### Supplementary Figures

#### Supplementary Fig. 1

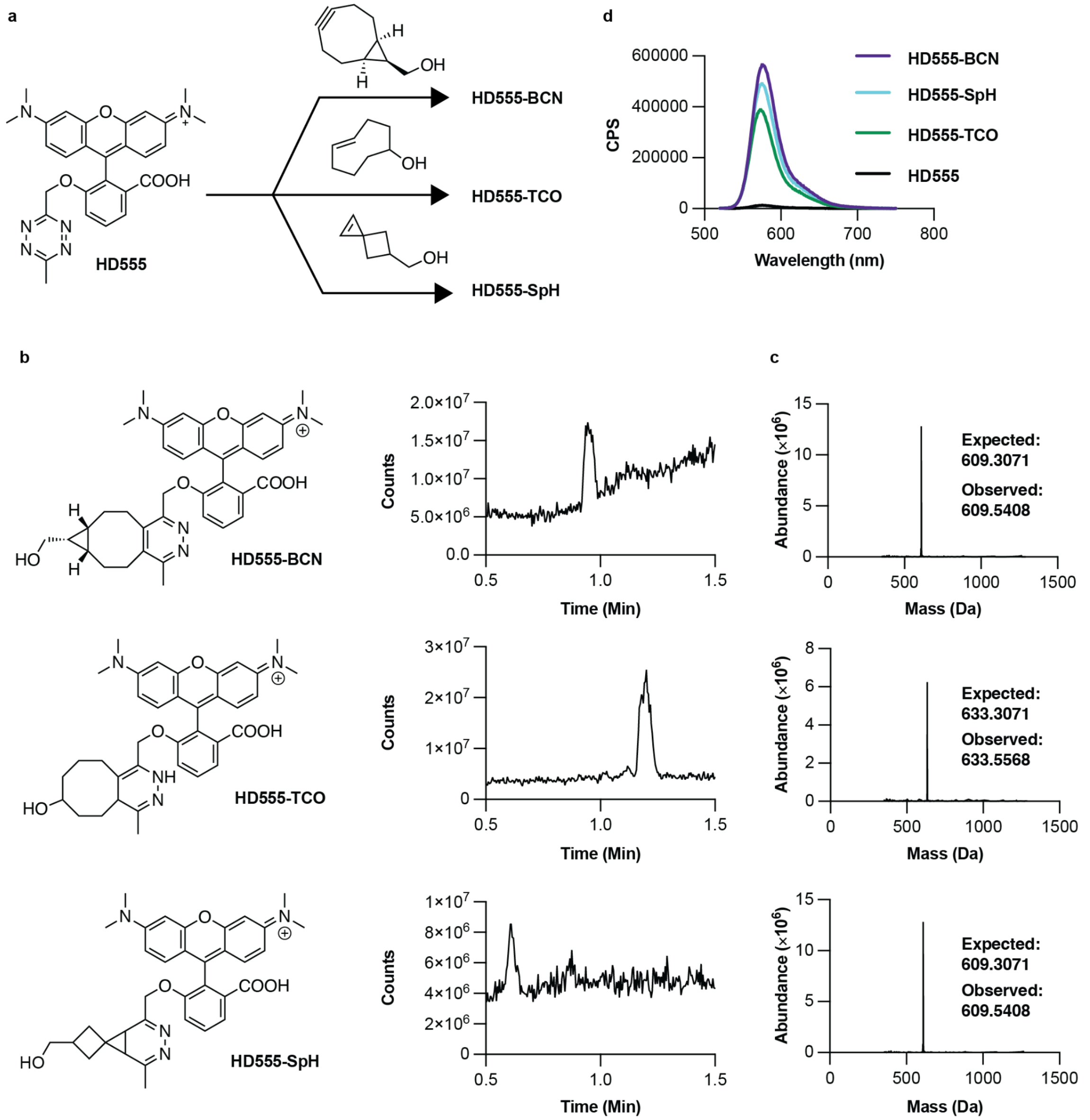

**Supplementary Fig. 1.** Characterization of HD555 reaction products in vitro. **a,** Reactions of HD555 with BCN, TCO, and SpH partners were performed using 5 µM HD555 and 50 µM of the indicated reaction partner in MilliQ water at 37 °C for 2 h. **b,** Product formation was monitored using a Waters Acquity SQD2 UPLC-MS system equipped with a 2.1 mm x 30 mm, CSH C18 column and eluents A ($H_2O$ with 0.1% $HCO_2H$) and B ($CH_3CN$ with 0.1% $HCO_2H$). Two µL of each HD555 reaction was injected directly on the LC-MS and resolved using a linear gradient of 10 to 95% B:A over 3.5 min. **c,** Deconvoluted mass spectra of HD555 reaction products. **d,** Overlaid emission spectra of HD555 reaction products (480 nm excitation, 520 to 750 nm emission).

### Supplementary Fig. 2

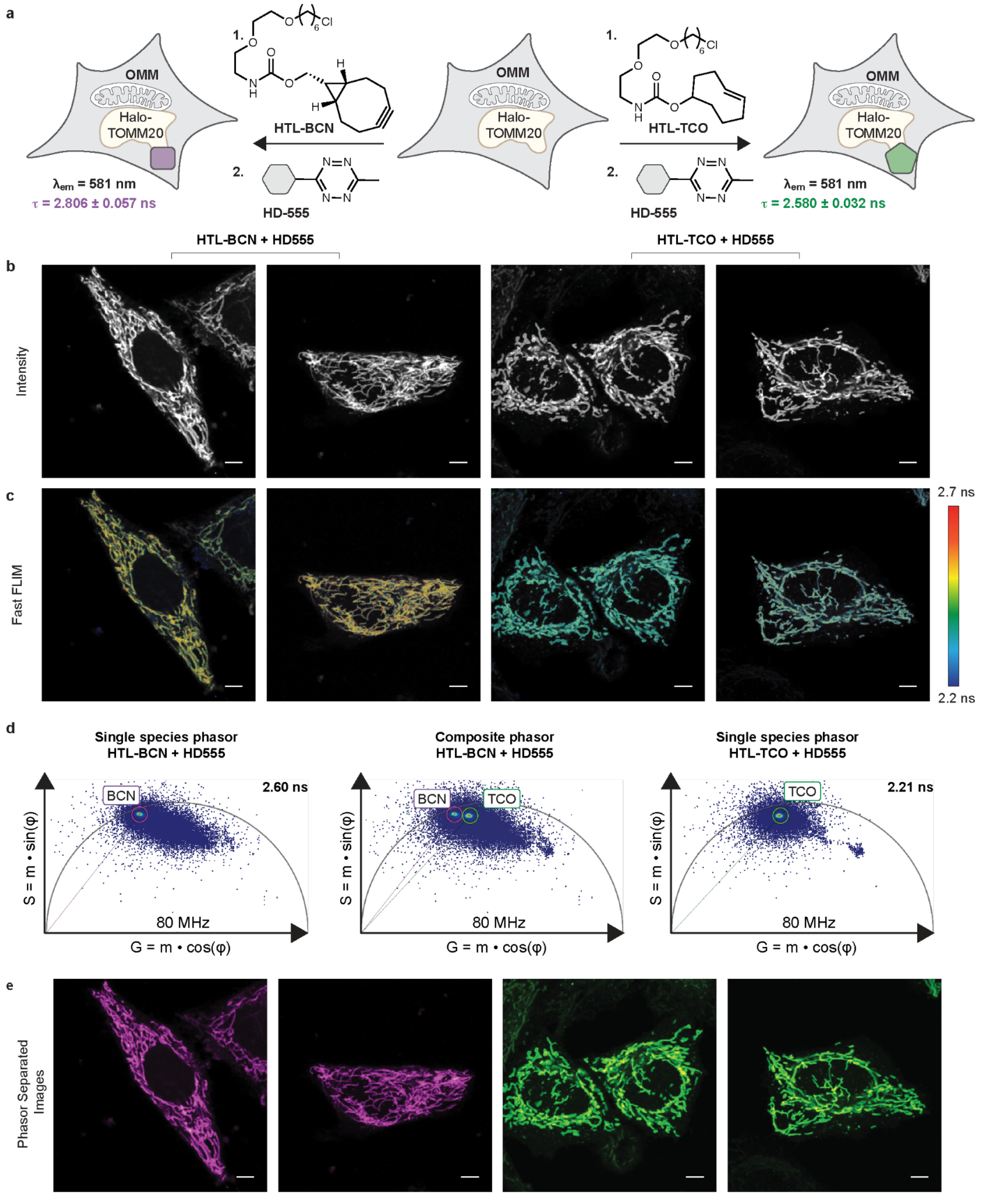

**Supplementary Fig. 2.** Turn-on correlates with fluorescence lifetime when HD555 is appended to HaloTag on the outer mitochondrial membrane (OMM). **a,** HeLa cells expressing HaloTag-TOMM20 were treated with HTL-TCO or HTL-BCN then HD555 and analyzed by detecting **b,** fluorescence intensity (Intensity) or **c,** average photon arrival time (Fast FLIM, color-coded). **d,** Single-species or composite phasor plots of datasets acquired when cells were treated with either HTL-TCO or HTL-BCN and then HD555. **e,** Phasor analysis was used to select densely populated regions on the phasor plot. Selected pixels in the image from the population with the longer fluorescence lifetimes are highlighted in magenta. Selected pixels in the image from the population with the shorter fluorescence lifetimes are highlighted in green. Scale bars, 5 µm.

### Supplementary Fig. 3

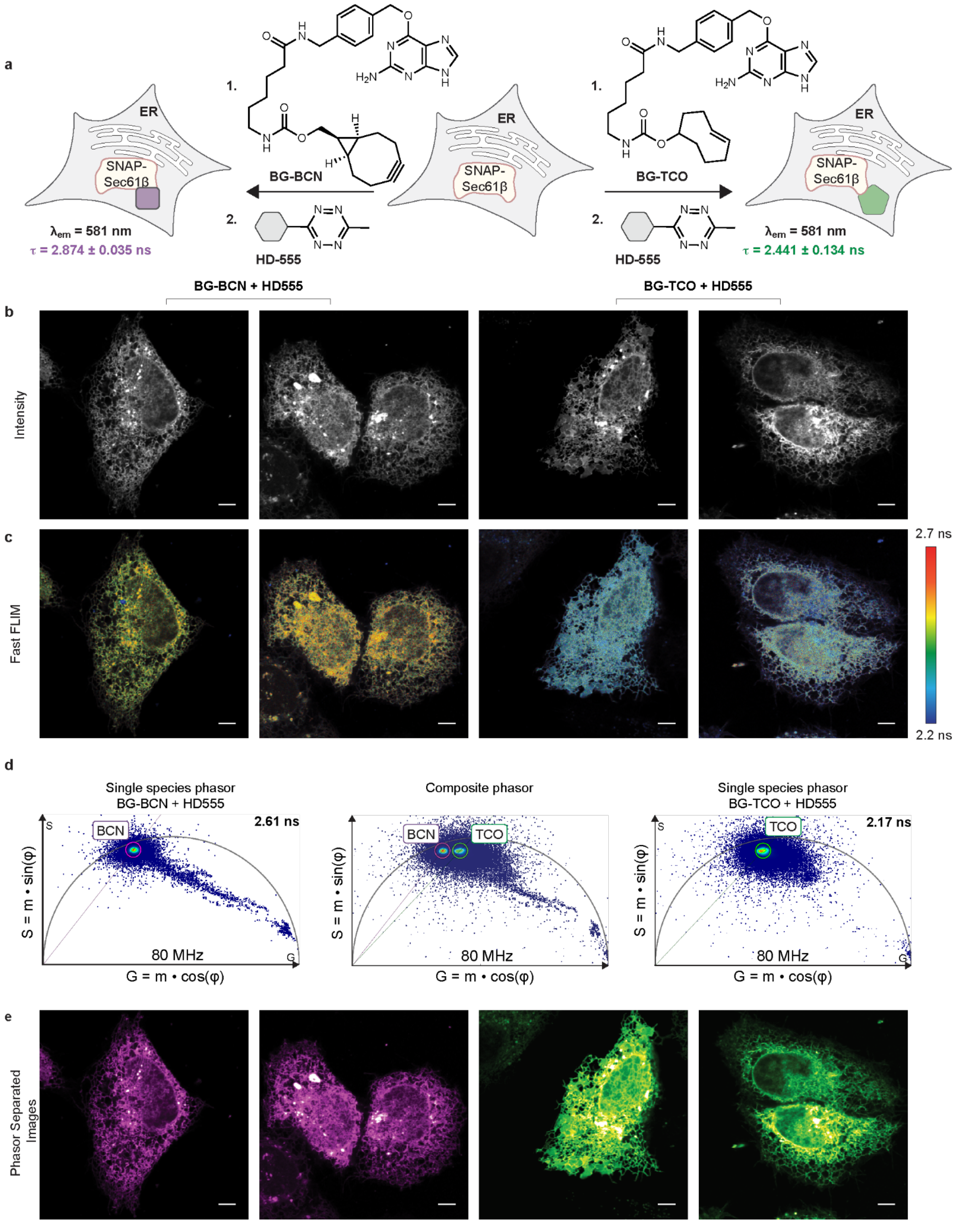

**Supplementary Fig. 3.** Turn-on correlates with fluorescence lifetime when HD555 is presented by SNAP-Tag on the ER membrane. **a,** HeLa cells expressing SNAP-Tag-Sec61β were treated with BG-TCO or BG-BCN then HD555 and analyzed by **b,** fluorescence intensity (Intensity) or **c,** average photon arrival time (Fast FLIM, color-coded). **d**, single-species or composite combined phasor plots of datasets acquired when cells were treated with HD555 and either BG-TCO or BG-BCN. **e,** Phasor analysis was used to select densely populated regions on the phasor plot. Selected pixels in the image from the population with the longer fluorescence lifetimes are highlighted in magenta. Selected pixels in the image from the population with the shorter fluorescence lifetimes are highlighted in green. Scale bars, 5 μm.

**Supplementary Fig. 4**

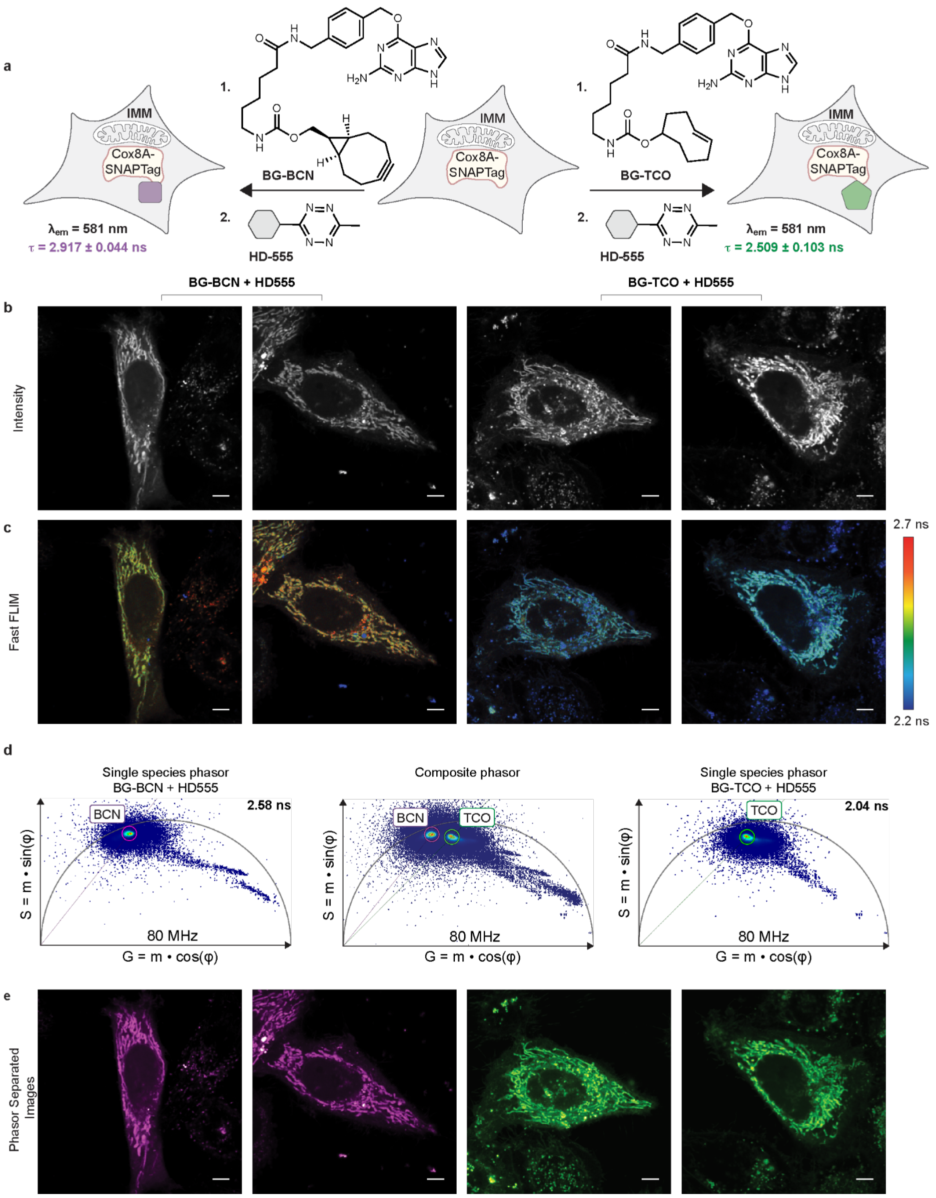

**Supplementary Fig. 4.** Turn-on correlates with fluorescence lifetime when HD555 is presented by SNAP-Tag on the mitochondrial inner membrane (IMM). **a,** HeLa cells expressing Cox8A-SNAP-Tag were treated with BG-TCO or BG-BCN then HD555 and analyzed by **b,** fluorescence intensity (Intensity or **c,** average photon arrival time (Fast FLIM, color-coded). **d**, single-species or composite combined phasor plots of datasets acquired when cells were treated with HD555 and either BG-TCO or BG-BCN. **e,** Phasor analysis was used to select densely populated regions on the phasor plot. Selected pixels in the image from the population with the longer fluorescence lifetimes are highlighted in magenta. Selected pixels in the image from the population with the shorter fluorescence lifetimes are highlighted in green. Scale bars, 5 μm.

### Supplementary Fig. 5

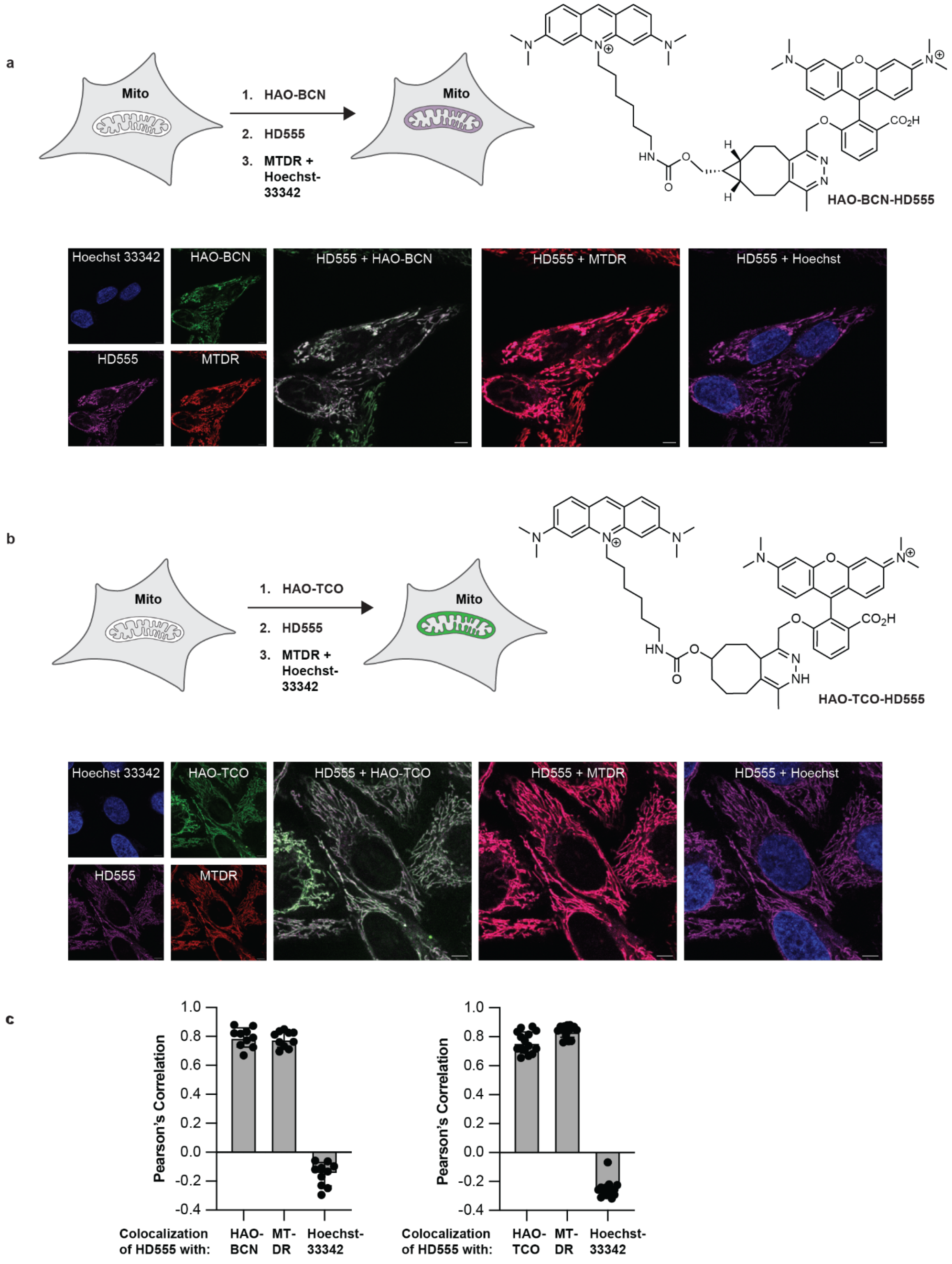

**Supplementary Fig. 5.** Upon reaction with **a**, HAO-BCN or **b**, HAO-TCO, the signals due to the HAO chromophore (502 nm excitation, 507 to 547 nm emission) colocalize with the signals due to the HD555 chromophore (561 nm excitation, 567 to 632 nm emission) and colocalize with the signals due to MitoTracker Deep Red (MTDR) (641 nm excitation, 646 to 829 nm emission). None of the aforementioned fluorophores colocalize with the signals due to Hoechst 33342 (405 nm excitation, 430 to 507 nm emission). **c,** Calculated Pearson's correlation coefficients of HAO-BCN (0.79 ± 0.07), MTDR (0.78 ± 0.06), and Hoechst 33342 (-0.1522 ± 0.08) with respect to HD555 for cells treated with HAO-BCN, MTDR, Hoechst 33342, and HD555. Calculated Pearson's correlation coefficients of HAO-TCO (0.76 ± 0.07), MTDR (0.84 ± 0.04), and Hoechst 33342 (-0.2531 ± 0.06) with respect to HD555 for cells treated with HAO-TCO, MTDR, Hoechst 33342, and HD555. Scale bars, 5 µm.

### Supplementary Fig. 6

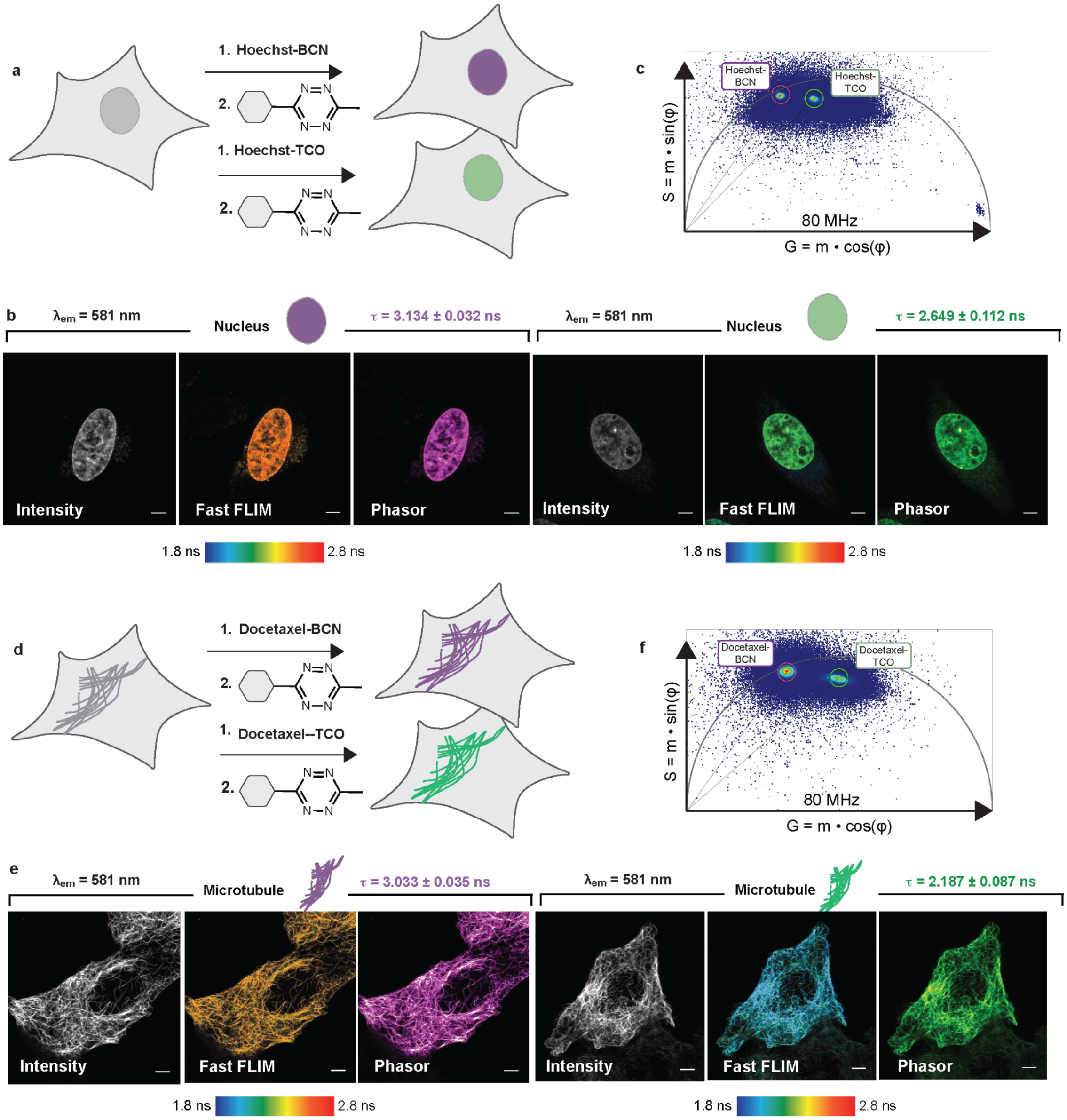

**Supplementary Fig. 6**. Turn-on correlates with fluorescence lifetime when HD555 is targeted to the nucleus with Hoechst or to microtubules with Docetaxel. **a,** HeLa cells were treated with Hoechst-TCO or Hoechst-BCN, then HD555, and analyzed by **b,** fluorescence intensity (Intensity) or average photon arrival time (Fast FLIM, color-coded). **c**, Overlaid phasor plot of images from Hoechst-TCO and Hoechst-BCN treated cells. **d,** HeLa cells were treated with Docetaxel-TCO or Docetaxel-BCN, then HD555, and analyzed by **e,** fluorescence intensity (Intensity) or average photon arrival time (Fast FLIM, color-coded). **f**, Overlaid phasor plot of images from Docetaxel-TCO and Docetaxel-BCN treated cells. Scale bars, 5 µm.

**Supplementary Fig. 7**

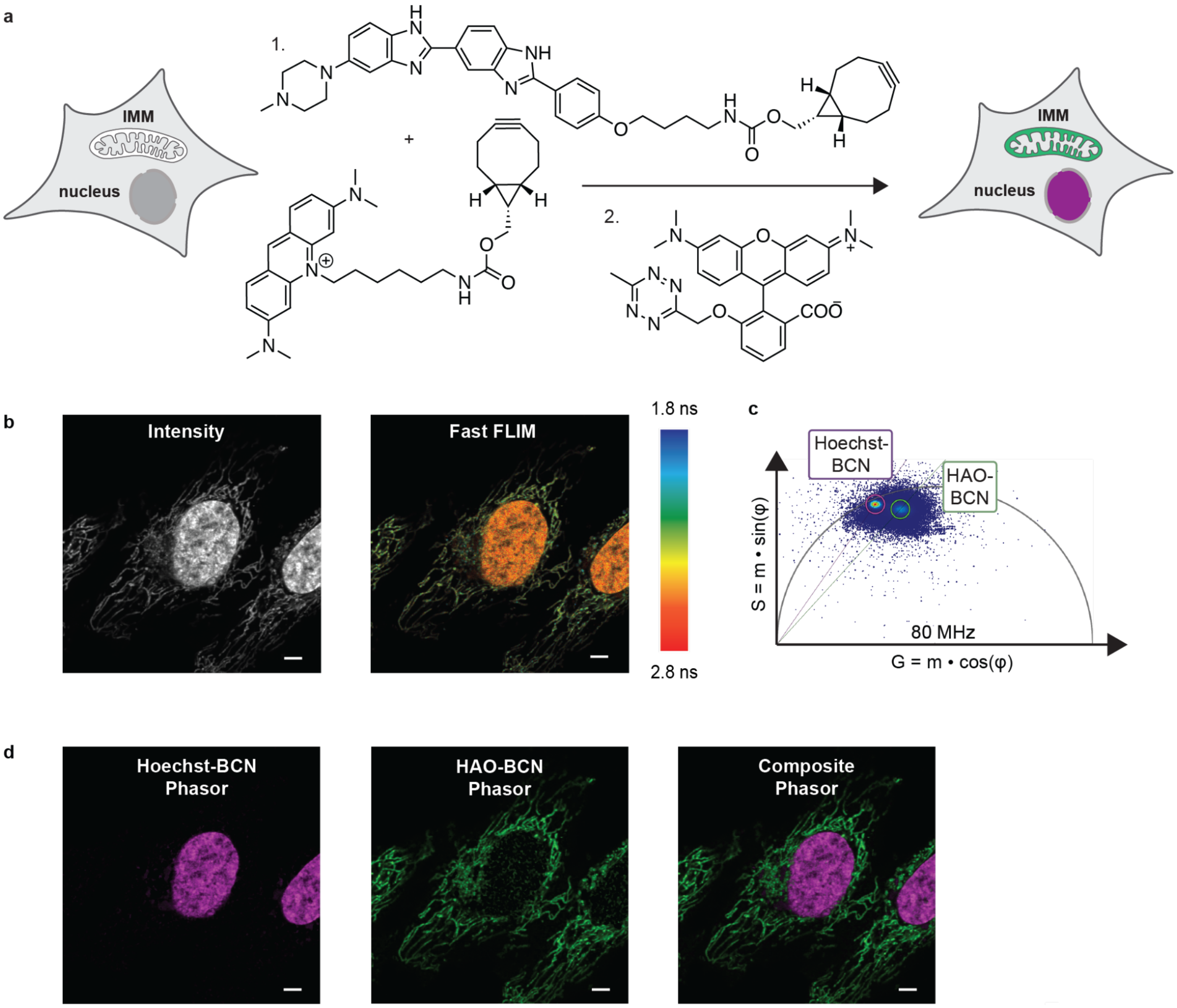

**Supplementary Fig. 7**. Microenvironment affects the fluorescence lifetimes of small molecule probes. **a,** HeLa cells were treated with Hoechst-BCN and HAO-BCN, then HD555, and analyzed by **b,** fluorescence intensity (Intensity) or average photon arrival time (Fast FLIM, color-coded). **c**, Phasor plot of images from Hoechst-BCN and HAO-BCN treated cells. **d,** Phasor-separated and composite images of Hoechst-BCN and HAO-BCN treated cells. Scale bars, 5 μm.

### Supplementary Fig. 8

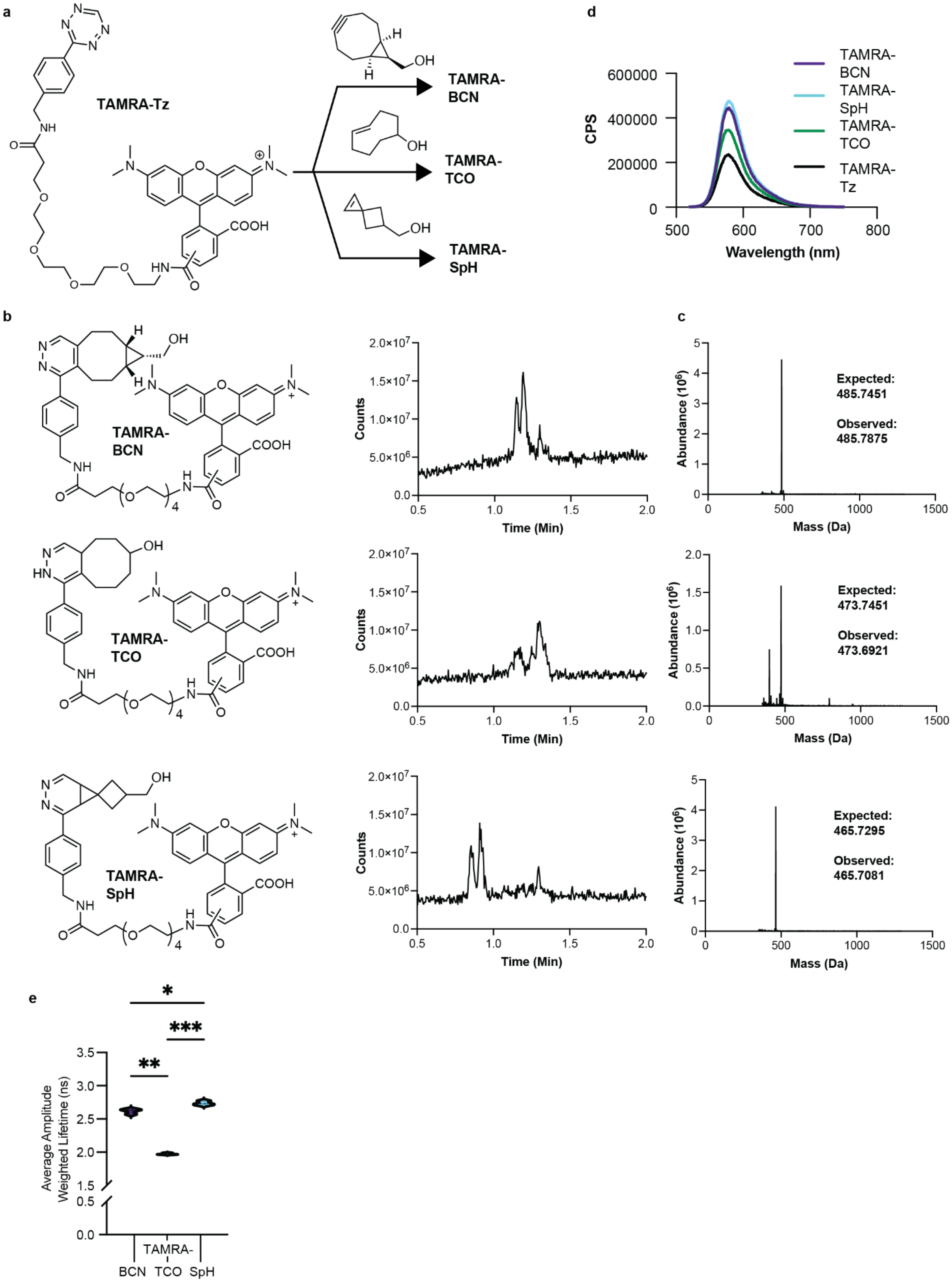

**Supplementary Fig. 8.** Characterization of TAMRA-Tz reaction products *in vitro*. **a,** Reactions of TAMRA-Tz with BCN, TCO, and SpH partners were performed using 5 µM TAMRA-Tz and 50 µM of the indicated reaction partner in MilliQ water at 37 °C for 2 h. **b,** Formation of the indicated TAMRA-Tz reaction products was monitored using a Waters Acquity SQD2 UPLC-MS system equipped with a 2.1 mm x 30 mm, CSH C18 column and eluents A ($H_2O$ with 0.1% $HCO_2H$) and B ($CH_3CN$ with 0.1% $HCO_2H$). Two µL of each TAMRA-Tz reaction was injected directly on the LC-MS and resolved using a linear gradient of 10 to 95% B:A over 3.5 min. **c,** Deconvoluted mass spectra of the TAMRA-Tz reaction products. **d,** Overlaid emission spectra of the TAMRA-Tz reaction products (480 nm excitation, 520 to 750 nm emission). **e,** Average amplitude-weighted lifetimes of TAMRA-Tz reaction products determined *in vitro*. Adjusted P values for TAMRA-BCN vs. TAMRA-TCO (**P = 0.0028), TAMRA-BCN vs. TAMRA-SpH (*P = 0.0367), and TAMRA-TCO vs. TAMRA-SpH (***P = 0.0001) calculated using Dunnett's T3 multiple comparisons test. Statistical significance was determined via Welch's and Brown-Forsythe ANOVA with post hoc Dunnett's T3 test accounting for heteroscedasticity.

### Supplementary Fig. 9

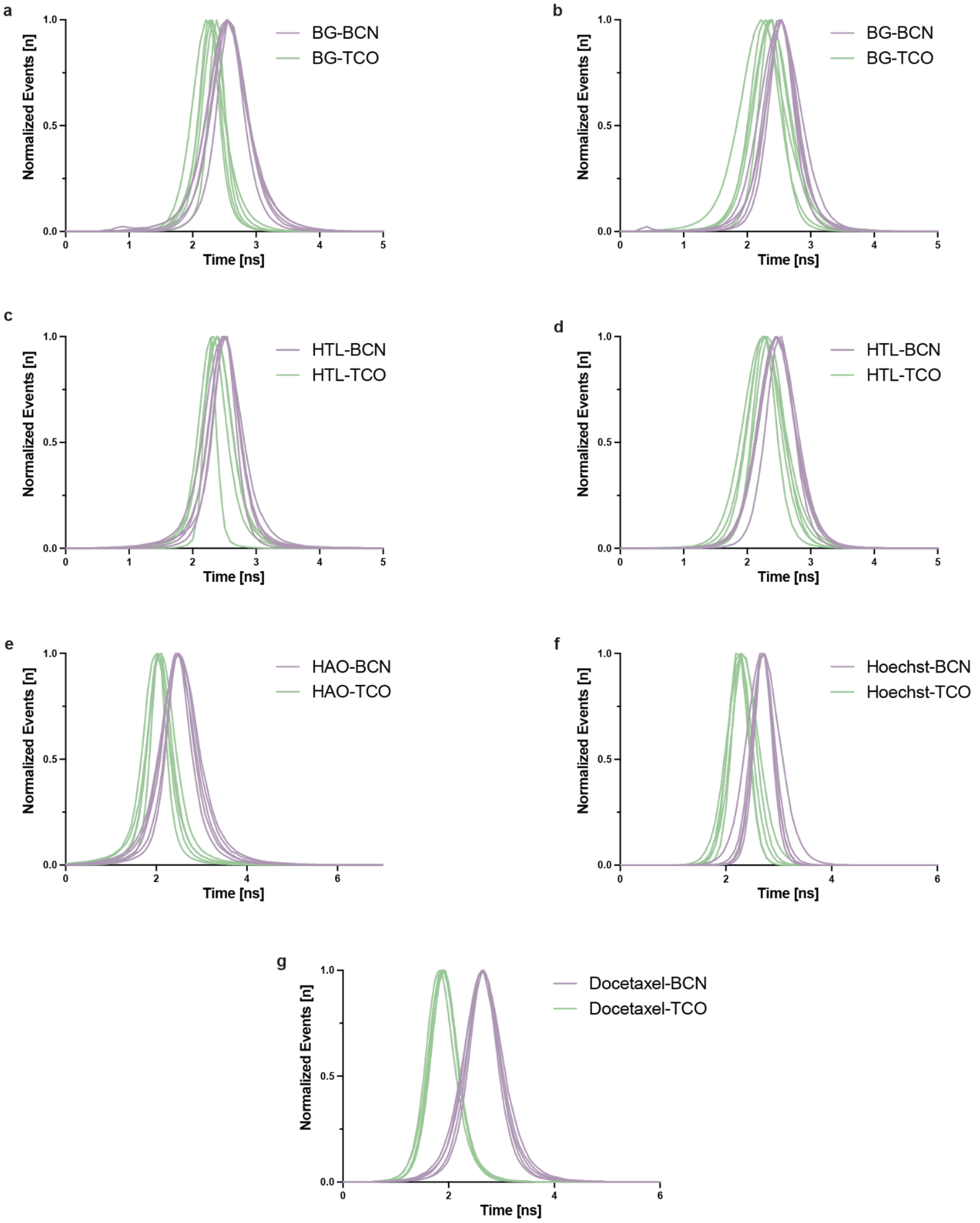

**Supplementary Fig. 9.** Normalized Fast FLIM histograms of **a,** COX8A-SNAPTag; **b,** Sec61β-SNAPTag; **c,** TOMM20-HaloTag; **d,** Sec61β-HaloTag; **e,** HAO; **f,** Hoechst; and **g,** Docetaxel. Self-labeling proteins were incubated with their respective BCN or TCO ligands (BG-BCN, BG-TCO, HTL-BCN, or HTL-TCO), then treated with HD555. Small molecule probes were functionalized with either BCN or TCO, then treated with HD555.

### Supplementary Tables

**Supplementary Table 1. Lifetime data for HD555 with bioorthogonal reaction partners in multiple contexts.**

| | | **Average intensity weighted lifetime** (ns) | **Average amplitude weighted lifetime** (ns) | **Phasor plot lifetime** (ns) | *n* |
|---|---|---|---|---|---|
| *In Vitro* | | | | | |
| | BCN | – | 2.934 ± 0.033 | 2.384 | 3 |
| | TCO | – | 2.001 ± 0.042 | 1.691 | 3 |
| | SpH | – | 2.435 ± 0.059 | 1.953 | 3 |
| *Self-labelling enzymes* | | | | | |
| **Halo-TOMM20** | HTL-BCN | 2.806 ± 0.057 | 2.438 ± 0.198 | 2.603 | 26 |
| | HTL-TCO | 2.580 ± 0.032 | 2.004 ± 0.107 | 2.221 | 30 |
| **Halo-Sec61β** | HTL-BCN | 2.753 ± 0.021 | 2.371 ± 0.112 | 2.566 | 27 |
| | HTL-TCO | 2.513 ± 0.026 | 1.989 ± 0.101 | 2.180 | 25 |
| **Cox8A-SNAP** | BG-BCN | 2.917 ± 0.044 | 2.452 ± 0.135 | 2.583 | 23 |
| | BG-TCO | 2.509 ± 0.103 | 1.656 ± 0.261 | 2.040 | 21 |
| **SNAP-Sec61β** | BG-BCN | 2.874 ± 0.035 | 2.448 ± 0.105 | 2.605 | 34 |
| | BG-TCO | 2.441 ± 0.134 | 1.763 ± 0.225 | 2.174 | 26 |
| *Small-molecule probes* | | | | | |
| | HAO-BCN | 2.819 ± 0.086 | 1.941 0.118 | 2.238 | 37 |
| | HAO-TCO | 2.226 ± 0.078 | 1.243 0.086 | 1.627 | 51 |
| | Hoechst-BCN | 3.134 ± 0.032 | 2.961 ± 0.031 | 2.905 | 32 |
| | Hoechst-TCO | 2.649 ± 0.112 | 1.502 ± 0.241 | 2.095 | 34 |
| | Docetaxel-BCN | 3.033 ± 0.035 | 2.812 ± 0.038 | 2.799 | 38 |
| | Docetaxel-TCO | 2.182 ± 0.087 | 1.190 ± 0.129 | 1.763 | 31 |

### Synthesis

#### General Considerations

All reactions were performed using flame-dried or oven-dried glassware under an atmosphere of anhydrous $N_2$ unless indicated otherwise. Microwave vials were purchased from Biotage; 4 mL vials refer to standard 1-dram glass vials and scintillation vials refer to standard 20 mL glass vials. Chemicals used for synthesis were purchased from commercial manufacturers (Combi-block Inc., Sigma-Aldrich®, BroadPharm®, ThermoScientific Inc., TCI America™). *N,N*-diisopropylethylamine (DIPEA), dry ethanol, dry tetrahydrofuran (THF), dry *N,N*-dimethylformamide (DMF), dry dichloromethane (DCM), and dry 1,4-dioxane were dispensed from a Sigma-Aldrich Sure/Seal™ container under a nitrogen atmosphere. Flash column chromatography was performed on a Teledyne ISCO CombiFlash® NextGen 300+ using RediSep® Silica (40–60 µm) or Alumina (40–60 µm) prepackaged columns. Preparative HPLC was performed on a Waters LC Prep 150 system equipped with a Waters 2707 autosampler, Waters Fraction Collector III and a reversed-phase C18 column (CSH C18 19 x 150 mm OBD Column 5 µm). Reversed-phase HPLC purifications used a gradient of water with 0.1% (v/v) trifluoroacetic acid (solvent A) and acetonitrile with 0.1% (v/v) trifluoroacetic acid (solvent B) at a flow rate of 20 mL/min.

$^{1}$H NMR and $^{13}$C NMR spectra were recorded using a Bruker NEO-500, NEO-501, AV-600, or AV-700 (frequencies listed in spectra). NMR spectra were analyzed using MestreLab MesReNova 15.0.1. Chemical shift values (δ) are reported in parts per million (ppm) relative to the solvent residual signals (reported individually for each compound). Reaction monitoring by LC/MS was carried out on a Low-resolution Waters Acquity UPLC-MS equipped with a dual atmospheric pressure chemical ionization / electrospray ionization mass spectrometry detector (SQD2), 200-800 nm UV/vis detector, a 300-1000 nm fluorescence detector, and an autosampler. Separations took place on a reverse-phase C18 column (Acquity UPLC HSS T3 C18 column, 1.8 mm particle size, 2.1 x 30 mm) using the following method: A linear gradient of 10-90% Solvent D over 3.5 min at a flow rate of 0.5 mL/min (Solvent C: 0.1% v/v formic acid in water and Solvent D: 0.1% v/v formic acid in acetonitrile). High-resolution mass spectrometry (HRMS) was performed on an Agilent 6530 Accurate-Mass Q-TOF LC/MS equipped with a reversed-phase column (YMC-Triart C18, 150 mm × 10 mm S-5 µm, 12 nm) using electrospray ionization in positive mode.

### 1. Synthesis of HTL-BCN and HTL-TCO

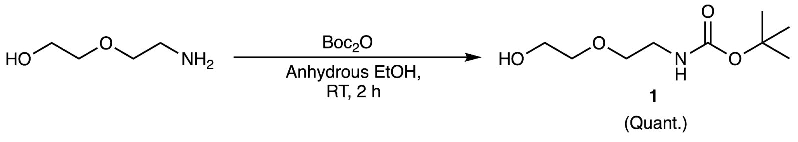

tert-butyl (2-(2-hydroxyethoxy)ethyl)carbamate **(1)**

The procedure was adapted from previously published methods[1]. To a 100 mL round-bottom flask equipped with a stir bar was added 2-(2-aminoethoxy)ethanol (1 g, 9.511 mmol, 1.0 equiv.) in anhydrous ethanol (0.5 M, 19.0 mL) and cooled to 0 °C in an ice bath. Di-tert-butyl dicarbonate (2.076 g, 9.511 mmol, 1.0 equiv.) was added dropwise and the mixture was stirred at 0 °C for 20 min. The reaction mixture was then allowed to warm to room temperature and stirred for an additional 2 h. After completion, the solvent was evaporated under reduced pressure. The resultant residue was diluted with water, and the organic material was extracted with DCM (4 × 20 mL). The combined organic layers were washed with brine, dried over anhydrous sodium sulfate ($Na_2SO_4$), filtered, and concentrated *in vacuo*. Residual solvent was removed under a stream of nitrogen overnight, providing the title compound as a colorless oil (1.95 g). The crude product was sufficiently pure and used directly in subsequent steps without further purification. Characterization agreed with previously reported spectra. $^1$H NMR (500 MHz, $CDCl_3$) δ 4.89 (s, 1H), 3.81 – 3.68 (m, 2H), 3.56 (dt, *J* = 12.7, 4.9 Hz, 4H), 3.33 (t, *J* = 5.2 Hz, 2H), 1.45 (s, 9H). $^{13}$C NMR (126 MHz, $CDCl_3$) δ 156.07, 79.46, 72.14, 70.32, 61.82, 40.42, 28.41. HRMS (ES+) calcd for ($C_9H_{19}NNaO_4^+$) $[M+Na]^+$ 228.12063, found 228.12173.

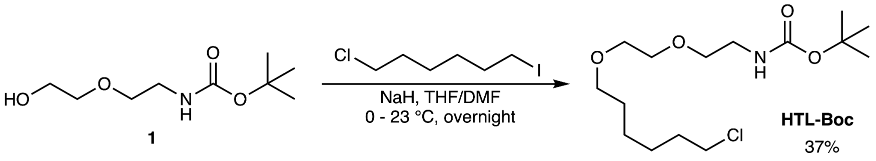

*tert*-butyl (2-(2-((6-chlorohexyl)oxy)ethoxy)ethyl)carbamate **(HTL-Boc)**

The procedure was adapted from previously published methods[1]. To a 5 mL microwave vial equipped with a stir bar was added **1** (0.500 g, 2.436 mmol), anhydrous THF (0.69 M, 3.53 mL), and DMF (1.390 M, 1.75 mL). The solution was stirred at 0 °C until equilibrated. Sodium hydride (60% dispersion in oil, 0.117 g, 2.923 mmol, 1.2 equiv.) was quickly added under nitrogen. Safety note: The addition of sodium hydride generates hydrogen gas. The vial was capped, sealed, and stirred for an additional 30 min. 1-chloro-6-iodohexane (0.841 g, 3.410 mmol, 1.4 equiv.) was added, and the reaction mixture was allowed to warm to room temperature and stirred for 18 h. The reaction was quenched with saturated aqueous $NH_4Cl$ and the organic material was extracted with DCM (3 x 50 mL). The combined organic layers were washed with brine, dried (anhydr. $Na_2SO_4$), filtered, and concentrated *in vacuo*. Purification by flash column chromatography (silica, 0–50% ethyl acetate in hexanes) yielded **HTL-Boc** (0.303 g, 37%) as a clear oil (Rf = 0.25, 70:30 hexanes/ethyl acetate, stains black with PMA). Characterization agreed with previously reported spectra. $^1$H NMR (500 MHz, $CDCl_3$) δ 4.99 (s, 1H), 3.60 (dd, *J* =

5.7, 3.0 Hz, 2H), 3.58 – 3.50 (m, 6H), 3.46 (t, *J* = 6.6 Hz, 2H), 3.34 – 3.28 (m, 2H), 1.77 (p, *J* = 6.9 Hz, 2H), 1.61 (p, *J* = 6.8 Hz, 2H), 1.46 (d, *J* = 8.5 Hz, 2H), 1.44 (s, 9H), 1.42 – 1.32 (m, 2H). $^{13}$C NMR (126 MHz, $CDCl_3$) δ 156.12, 79.32, 71.42, 70.42, 70.35, 70.17, 45.18, 40.49, 32.67, 31.07, 29.57, 28.56, 26.83, 25.56. HRMS (ES+) calcd for ($C_{15}H_{30}ClNNaO_4^+$) $[M+Na]^+$ 346.17556, found 346.17647.

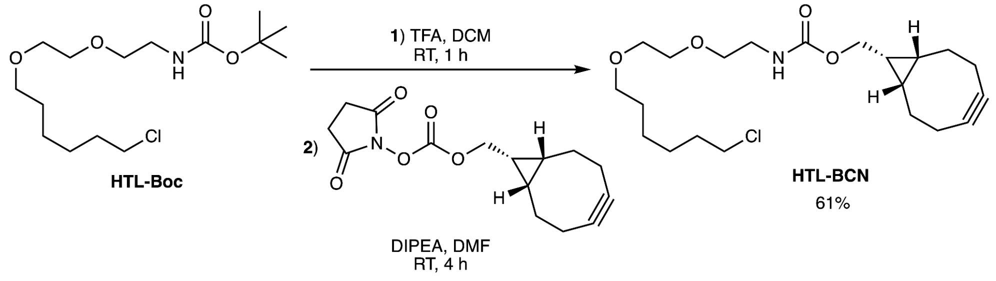

((1*R*,8*S*,9*s*)-bicyclo[6.1.0]non-4-yn-9-yl)methyl (2-(2-((6-chlorohexyl)oxy)ethoxy)ethyl) carbamate **(HTL-BCN)**

To a 4 mL vial equipped with a stir bar was added **HTL-Boc** (50 mg, 0.154 mmol), DCM (1.54 mL), and TFA (0.118 mL, 1.544 mmol, 10 equiv.). The reaction was stirred for 1 h (monitored by TLC and LCMS). The reaction mixture was concentrated *in vacuo*, and the deprotected amine was used without further purification. In a separate 4 mL vial, BCN-NHS ester (54 mg, 0.185 mmol, 1.2 equiv.) was dissolved in DMF (0.1 M, 1.54 mL), and DIPEA (0.249 g, 1.930 mmol, 12.5 equiv.) was added. The deprotected chloroalkane intermediate was dissolved separately in DMF (1.54 mL), and the BCN-NHS ester solution was quickly added. The reaction was stirred at room temperature and monitored by TLC (basic alumina, 75:25 hexanes/ethyl acetate; stains: $KMnO_4$, $I_2$) and LCMS until completion (4 h). Upon completion, the reaction was concentrated *in vacuo* and purified by flash column chromatography (basic alumina, 0–50% ethyl acetate in hexanes). The title compound was isolated as a colorless oil (37 mg, 61%).

$^1$H NMR (400 MHz, CDCl3) δ 5.18 (s, 1H), 4.13 (d, J = 8.1 Hz, 2H), 3.63 – 3.53 (m, 4H), 3.54 (s, 2H), 3.55 – 3.44 (m, 3H), 3.44 (d, J = 6.7 Hz, 2H), 3.36 (q, J = 5.4 Hz, 2H), 2.24 (dqd, J = 15.4, 9.7, 3.6 Hz, 5H), 1.76 (p, J = 7.0 Hz, 2H), 1.60 (p, J = 6.9 Hz, 3H), 1.50 – 1.27 (m, 5H), 0.99 – 0.86 (m, 2H). $^{13}$C NMR (126 MHz, CDCl3) δ 156.12, 79.32, 71.42, 70.42, 70.35, 70.17, 45.18, 40.49, 32.67, 31.07, 29.57, 28.56, 26.83, 25.56. HRMS (ES+) calcd for ($C_{21}H_{34}ClNO_4$) $[M+K^+]$ 438.18079, found 438.37069.

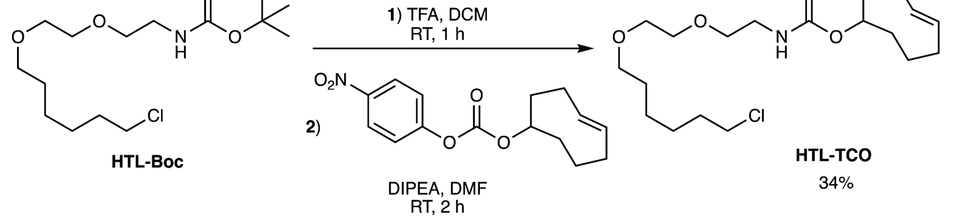

(*E*)-cyclooct-4-en-1-yl (2-(2-((6-chlorohexyl)oxy)ethoxy)ethyl)carbamate **(HTL-TCO)**

To a 4 mL vial equipped with a stir bar was added **HTL-Boc** (50 mg, 0.154 mmol) dissolved in DCM (1.54 mL) and TFA (0.118 mL, 1.544 mmol, 10 equiv.). The reaction was stirred for 1 h (monitoring and completion verified by TLC and LCMS). The reaction mixture was concentrated *in vacuo*, and the deprotected amine was used without further purification. To a separate 4 mL vial was added TCO-PNB ester (54 mg, 0.185 mmol, 1.2 equiv.) dissolved in DMF (0.1 M, 1.54 mL) and DIPEA (0.249 g, 1.930 mmol, 12.5 equiv.). The deprotected chloroalkane intermediate was dissolved separately in DMF (1.54 mL), and the TCO-PNB ester solution was quickly added. The reaction was stirred at room temperature and monitored by TLC (silica, 50:50 hexanes/ethyl acetate; stains: $KMnO_4$, $I_2$) and LCMS until completion (2 h). Upon completion, the reaction was concentrated and purified by column chromatography (basic alumina, 0–50% ethyl acetate in hexanes). The title compound was isolated as a yellowish oil (17.9 mg, 34%). $^1$H NMR (500 MHz, $CDCl_3$) δ 5.61 – 5.44 (m, 2H), 5.07 (t, *J* = 6.0 Hz, 1H), 4.32 (dd, *J* = 11.0, 5.9 Hz, 1H), 3.62 – 3.56 (m, 2H), 3.58 – 3.49 (m, 6H), 3.44 (t, *J* = 6.6 Hz, 2H), 3.37 – 3.30 (m, 2H), 2.39 – 2.27 (m, 3H), 2.04 – 1.98 (m, 1H), 2.00 – 1.85 (m, 3H), 1.81 – 1.71 (m, 3H), 1.74 – 1.67 (m, 1H), 1.66 – 1.40 (m, 5H), 1.43 – 1.31 (m, 2H), 1.26 (s, 1H). $^{13}$C NMR (126 MHz, $CDCl_3$) δ 156.12, 79.32, 71.42, 70.42, 70.35, 70.17, 45.18, 40.49, 32.67, 31.07, 29.57, 28.56, 26.83, 25.56. HRMS (ES+) calcd for ($C_{19}H_{34}ClNNaO_4$) $[M+Na]^+$ 398.20686, found 398.20199.

### 2. Synthesis of BG-BCN and BG-TCO

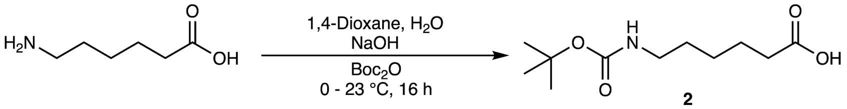

**2**

92%

6-((tert-butoxycarbonyl)amino)hexanoic acid **(2)**

To a 500 mL round-bottom flask equipped with a stir bar was added NaOH (1.1 eq, 189 mg, 4.738 mmol) dissolved in dioxane/water (90.0 mL, 2:1). 6-aminocaproic acid (1 eq, 565 mg, 4.307 mmol) was then added, and the reaction vessel was cooled to 0 °C. Di-tert-butyl dicarbonate (1.1 eq, 1.04 g, 4.738 mmol) was added to the flask in portions. The reaction mixture was stirred for 30 min, then warmed up to ambient temperature over 18 h. The mixture was concentrated, redissolved in water (50 mL), and then washed with ethyl acetate (3 × 50.0 mL). The aqueous layer was then acidified to pH 2 with aqueous 0.1 M HCl. The organic material was extracted with ethyl acetate (3 × 50 mL). Organic layers were combined and dried with Na2SO4, filtered, and concentrated *in vacuo* to yield the title compound (922 mg, 92%) as a colourless oil (crystallizes as a white solid upon drying and cooling). Spectra agree with previously published data[2]. $^1$H NMR (500 MHz, MeOD) δ 3.03 (t, *J* = 6.6 Hz, 2H), 2.33 – 2.26 (m, 2H), 1.61 (h, *J* = 6.4 Hz, 2H), 1.51 – 1.45 (m, 3H), 1.43 (s, 8H), 1.35 (h, *J* = 7.0 Hz, 2H). $^{13}$C NMR (126 MHz, $CDCl_3$) δ 156.12, 79.32, 71.42, 70.42, 70.35, 70.17, 45.18, 40.49, 32.67, 31.07, 29.57, 28.56, 26.83, 25.56. HRMS (ES+) calcd for ($C_{11}H_{21}NNaO_4^+$) $[M+Na]^+$ 254.13628, found 254.13735.

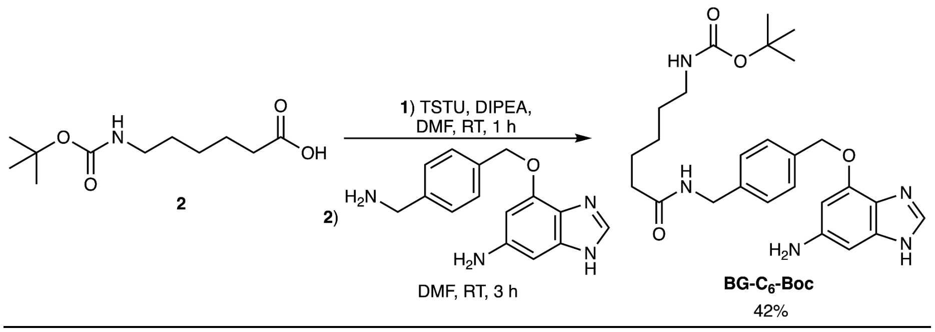

<u>*tert*-butyl (6-((4-(((2-amino-9*H*-purin-6-yl)oxy)methyl)benzyl)amino)-6-oxohexyl)carbamate **(BG-C$_6$-Boc)**</u>

To a 4 mL vial equipped with a stir bar was added Boc-aminocaprioic acid (**2**, 0.250 g, 1.082 mmol, 1.0 equiv.) dissolved in DMF (2.4 mL) and DIPEA (0.885 mL, 5.083 mmol, 4.7 equiv.). The mixture was stirred and equilibrated. TSTU (0.391 g, 1.298 mmol, 1.2 equiv.) was added to the vial. Complete conversion to the NHS-ester was observed after 1 h via LCMS. 6-((4-(aminomethyl)benzyl)oxy)-9*H*-purin-2-amine hydrochloride (0.409 g, 1.514 mmol, 1.4 equiv.) was then dissolved separately in DMF (2.4 mL, 0.226 M for total reaction volume), added dropwise to the NHS-ester solution, and stirred at room temperature for 3 h. Reaction progress and completion was monitored by LCMS. After completion, the mixture was concentrated *in vacuo*, redissolved in 50:50 acetonitrile/water containing 0.1% formic acid, and purified via reverse-phase HPLC (5–95% Solvent B over 20 min). The fractions were combined and lyophilized, affording the title compound (0.220 g, 42%) as a white solid. $^{1}$H NMR (500 MHz, MeOD) δ 7.85 (s, 0H), 7.53 – 7.48 (m, 1H), 7.34 – 7.28 (m, 1H), 5.55 (s, 1H), 4.38 (s, 1H), 3.03 (t, $J$ = 7.0 Hz, 1H), 2.25 (t, $J$ = 7.5 Hz, 1H), 1.65 (p, $J$ = 7.6 Hz, 1H), 1.54 – 1.45 (m, 1H), 1.44 (s, 4H), 1.40 – 1.30 (m, 1H). $^{13}$C NMR (126 MHz, MeOD) δ 174.61, 160.29, 157.16, 138.71, 135.52, 128.30, 127.28, 78.41, 67.25, 47.89, 42.42, 39.79, 35.57, 29.24, 27.39, 26.05, 25.33. HRMS (ES+) calcd for ($C_{24}H_{34}N_7O_4^+$) $[M+H]^+$ 484.26668, found 484.26757.

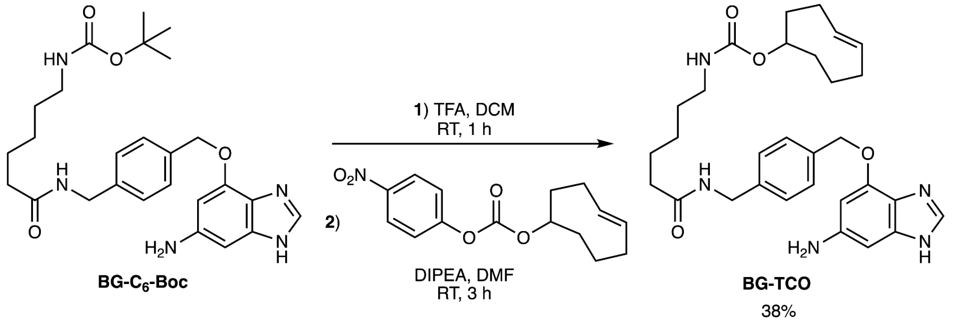

<u>(E)-cyclooct-4-en-1-yl (6-((4-(((2-amino-9H-purin-6-yl)oxy)methyl)benzyl)amino)-6-oxohexyl)carbamate **(BG–TCO)**</u>

Prepared similarly to **HTL-BCN**—adapted procedure below:
To a 4 mL vial equipped with a stir bar was added **BG-C$_6$-Boc** (35 mg, 0.072 mmol, 1 eq.) dissolved in DCM (1.54 mL) and TFA (0.118 mL, 1.544 mmol). The reaction was stirred for 1 h

(complete deprotection monitored by TLC and LCMS). The reaction mixture was concentrated *in vacuo*, and the deprotected amine was used without further purification. To a separate 4 mL vial was added TCO-PNB ester (21 mg, 0.072 mmol, 1 equiv.) dissolved in DMF (0.226 M, 0.320 mL) and DIPEA (0.076 mL, 0.434 mmol, 6 equiv.). The deprotected chloroalkane intermediate was dissolved separately in DMF (0.320 mL), and the TCO-PNB ester solution was quickly added. The reaction was stirred at room temperature and full conversion (3 h) was verified by LCMS. After completion, the mixture was concentrated *in vacuo*, redissolved in 50:50 acetonitrile/water containing 0.1% formic acid, and purified via reverse-phase HPLC (5–95% Solvent B over 20 min). The fractions were combined and lyophilized affording the title compound (15 mg, 38%) as a white solid. $^{1}$H NMR (500 MHz, MeOD) δ 8.34 (s, 1H), 7.54 – 7.49 (m, 2H), 7.35 – 7.30 (m, 2H), 5.64 (s, 2H), 5.57 (ddd, *J* = 15.1, 10.2, 4.6 Hz, 1H), 5.44 (ddd, *J* = 16.0, 11.0, 3.6 Hz, 1H), 4.37 (s, 2H), 4.36 – 4.24 (m, 1H), 3.04 (t, *J* = 6.9 Hz, 2H), 2.35 – 2.29 (m, 2H), 2.24 (t, *J* = 7.5 Hz, 2H), 1.92 (dddd, *J* = 23.3, 15.4, 8.2, 3.3 Hz, 3H), 1.76 – 1.51 (m, 4H), 1.47 (p, *J* = 7.2 Hz, 2H), 1.37 – 1.29 (m, 2H). $^{13}$C NMR (126 MHz, MeOD) δ 176.05, 162.54, 162.25, 161.10, 158.72, 158.04, 153.46, 143.78, 141.06, 136.07, 135.22, 133.72, 130.25, 128.79, 118.99, 116.67, 107.92, 81.54, 70.95, 49.45, 49.28, 43.71, 42.21, 41.47, 39.63, 36.92, 35.15, 33.48, 32.10, 30.68, 30.63, 27.38, 26.67, 24.21. HRMS (ES+) calcd for ($C_{28}H_{37}N_7O_4$) $[M+H]^+$ 536.29798, found 536.28416.

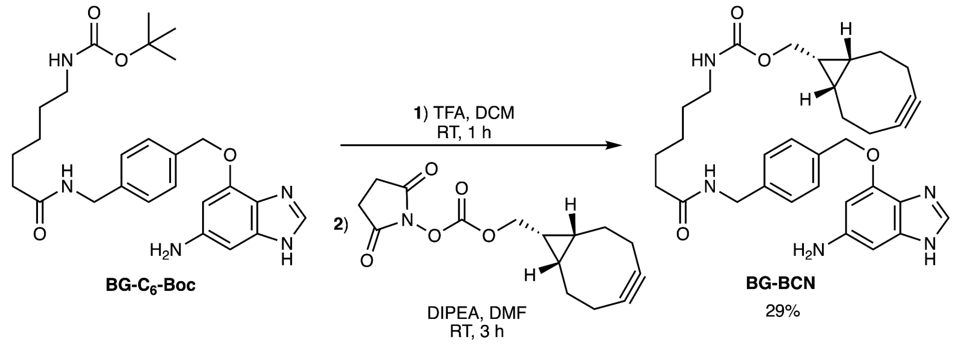

<u>((1*R*,8*S*,9*s*)-bicyclo[6.1.0]non-4-yn-9-yl)methyl (6-((4-(((2-amino-9*H*-purin-6-yl)oxy)methyl) benzyl) amino)-6-oxohexyl) carbamate **(BG–BCN)**</u>

Prepared similarly to **HTL-BCN**—adapted procedure below:

To a 4 mL vial equipped with a stir bar was added **BG-$C_6$-Boc** (35 mg, 0.072 mmol, 1 eq.) dissolved in DCM (1.54 mL) and TFA (0.118 mL, 1.544 mmol). The reaction was stirred until complete deprotection was observed (1 h) via TLC and LCMS. The reaction mixture was concentrated *in vacuo*, and the deprotected amine was used without further purification. To a separate 4 mL vial was added BCN-NHS ester (21 mg, 0.072 mmol, 1 equiv.) dissolved in DMF (0.226 M, 0.320 mL) and DIPEA (0.076 mL, 0.434 mmol, 6 equiv.). The deprotected chloroalkane intermediate was dissolved separately in DMF (0.320 mL), and the BCN-NHS ester solution was quickly added. The reaction was stirred at room temperature and monitored by LCMS until completion (3 h). After completion, the mixture was concentrated *in vacuo*, redissolved in 50:50 acetonitrile/water containing 0.1% formic acid, and purified via reverse-phase HPLC (5–95% Solvent B over 20 min). The fractions were combined and lyophilized affording the title compound (12 mg, 29%) as a white solid. $^1$H NMR (500 MHz, MeOD) δ 8.32 (s, 1H), 7.54 – 7.48 (m, 2H), 7.36 – 7.30 (m, 2H), 5.64 (s, 2H), 4.37 (s, 2H), 4.13 (dd, *J* = 12.9, 8.2 Hz, 2H), 3.07 (t, *J* = 7.0 Hz, 3H), 2.68 – 2.44 (m, 1H), 2.38 – 2.09 (m, 6H), 2.05 – 1.75 (m, 2H), 1.71 – 1.54 (m, 5H), 1.50 (p, *J* = 7.1 Hz, 3H), 1.35 (dd, *J* = 11.8, 4.5 Hz, 3H), 1.36 – 1.27 (m, 1H), 1.27 – 1.17 (m, 1H), 1.03 – 0.65 (m, 2H). $^{13}$C NMR (126 MHz, MeOD) δ 174.64, 159.72, 152.38, 142.10, 139.65, 133.96, 128.82, 127.39, 98.08, 69.42, 62.15, 61.79, 48.23, 48.06, 47.88, 45.04, 42.32, 40.14, 39.72, 35.54, 29.23, 28.75, 28.16, 25.98, 25.28, 23.24, 20.51, 19.96, 19.46, 18.32, 17.87, 17.55. HRMS (ES+) calcd for ($C_{30}H_{37}N_7O_4$) $[M+H]^+$ 560.29798, found 560.28841.

**Synthesis of Hexyl Acridine Orange (HAO) derivatives**

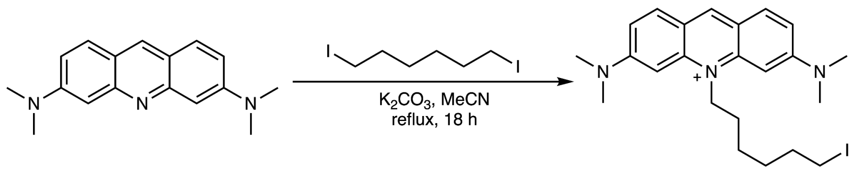

<u>3,6-bis(dimethylamino)-10-(6-iodohexyl)acridin-10-ium **(HAO-iodide)**</u>

Acridine Orange free base was prepared according to the procedure published in Zheng et al[3]. To a 25 mL microwave vial equipped with a stir bar was added Acridine Orange free base (0.100 g, 0.377 mmol, 1.0 equiv.), potassium carbonate (0.062 g, 0.452 mmol, 1.2 equiv.), and anhydrous acetonitrile (7.54 mL, 0.05 M). The mixture was stirred for 15 min to equilibrate. Diiodo-hexane (1.274 g, 0.621 mL, 3.768 mmol, 10.0 equiv.) was then added to the mixture dropwise. The reaction mixture was heated to 85 °C and stirred for 18 h. The reaction mixture was then filtered and concentrated *in vacuo*. The compound was purified by flash column chromatography (neutral alumina, 0–10% methanol in DCM), with an isocratic hold upon elution of the product. The title compound (124 mg, 68%) was isolated as a waxy orange solid. $^1$H NMR (600 MHz, $CDCl_3$) δ 8.71 (s, 1H), 7.90 (d, *J* = 9.2 Hz, 2H), 7.03 (dd, *J* = 9.3, 2.1 Hz, 2H), 6.63 (d, *J* = 2.2 Hz, 2H), 4.90 (t, *J* = 8.1 Hz, 2H), 3.33 (s, 12H), 3.24 (t, *J* = 6.8 Hz, 2H), 1.98 (p, *J* = 7.6 Hz, 2H), 1.87 (h, *J* = 7.1 Hz, 2H), 1.75 (p, *J* = 7.6 Hz, 2H), 1.60 (d, *J* = 9.9 Hz, 6H). $^{13}$C NMR (151 MHz, $CDCl_3$) δ 155.70, 143.09, 142.80, 133.40, 117.30, 114.23, 93.15, 48.46, 41.45, 33.13, 30.51, 26.26, 26.21, 7.78. HRMS (ES+) calcd for ($C_{23}H_{31}IN_3^+$) $[M]^+$ 476.15572, found 476.15167.

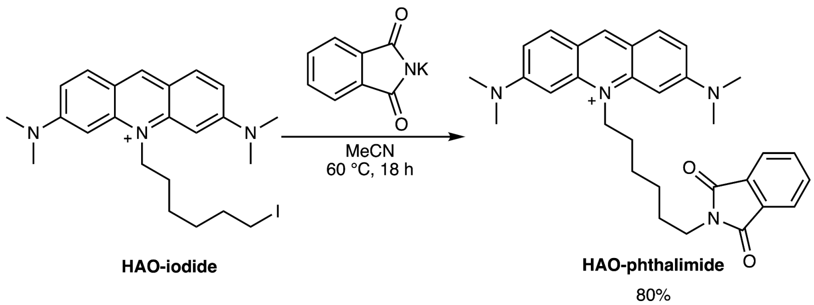

3,6-bis(dimethylamino)-10-(6-(1,3-dioxoisoindolin-2-yl)hexyl)acridin-10-ium **(HAO-phthalimide)**

To a 25 mL microwave vial equipped with a stir bar was added **HAO-iodide** (0.382 g, 0.802 mmol, 1.0 equiv.) and dried for 18 h *under vacuum.* Potassium phthalimide (0.594 g, 3.207 mmol, 4.0 equiv.) and anhydrous acetonitrile (8.02 mL, 0.1 M) were then added, and the reaction mixture was stirred at 60 °C for 18 h. The reaction mixture was cooled and concentrated *in vacuo*. The crude was resuspended in DCM and the organic layer was washed with water. The organic layer was then dried over anhydrous $Na_2SO_4$, gravity filtered, and concentrated *in vacuo*. The crude was purified using flash column chromatography (neutral alumina, 0–5% methanol/DCM). The title compound was isolated as an orange solid (0.40 g, 80% yield). $^1$H NMR (500 MHz, $CDCl_3$) δ 8.70 (s, 1H), 7.87 (d, *J* = 9.3 Hz, 2H), 7.76 (dd, *J* = 5.4, 3.1 Hz, 2H), 7.68 (dd, *J* = 5.5, 3.0 Hz, 2H), 7.00 (dd, *J* = 9.3, 2.1 Hz, 2H), 6.49 (d, *J* = 2.2 Hz, 2H), 4.73 (t, *J* = 8.0 Hz, 2H), 3.64 (t, *J* = 7.0 Hz, 2H), 3.28 (s, 11H), 1.91 (d, *J* = 16.0 Hz, 2H), 1.74 – 1.63 (m, 5H), 1.42 (qd, *J* = 9.7, 4.5 Hz, 2H). $^{13}$C NMR (151 MHz, $CDCl_3$) δ 155.70, 143.09, 142.80, 133.40, 117.30, 114.23, 93.15, 48.46, 41.45, 33.13, 30.51, 26.26, 26.21, 7.78. HRMS (ES+) calcd for ($C_{31}H_{35}N_4O_2^+$) $[M]^+$ 495.27545, found 495.27591.

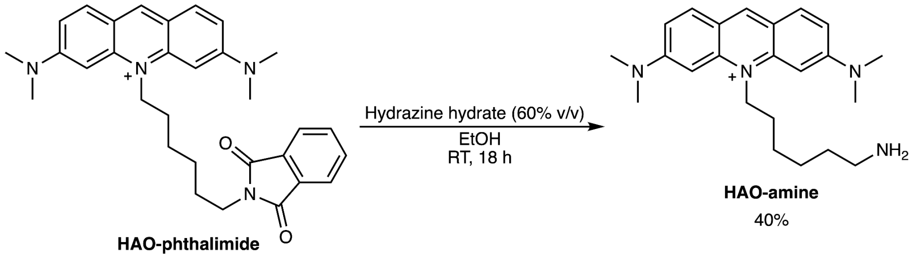

10-(6-aminohexyl)-3,6-bis(dimethylamino)acridin-10-ium **(HAO-amine)**

To a 25 mL round-bottom flask equipped with a stir bar was added **HAO-phthalimide** (0.120 g, 0.242 mmol) dissolved in a minimal amount of ethanol (<5 mL), and the solution was stirred for 5 min. Hydrazine hydrate—60% v/v—(0.048 g, 0.079 mL, 0.968 mmol, 4.0 equiv.) was then added to the flask. The reaction was stirred overnight (18 h), and completion was indicated by the formation of a white precipitate; product formation was confirmed by LCMS. The reaction mixture was filtered and purified directly via reverse-phase HPLC (0-50% Solvent B over 30 min), yielding the title compound (36 mg, 40%) as an orange solid. $^1$H NMR (600 MHz, MeOD) δ 8.66 (s, 1H), 7.91 (d, *J* = 9.3 Hz, 2H), 7.28 (dd, *J* = 9.3, 2.2 Hz, 2H), 6.68 (d, *J* = 2.4 Hz, 2H), 4.74 – 4.69 (m, 2H), 3.33 (s, 12H), 2.95 (t, *J* = 7.7 Hz, 2H), 2.08 – 2.00 (m, 2H), 1.77 – 1.66 (m,

4H), 1.57 (dddd, $J$ = 14.8, 9.4, 6.6, 3.5 Hz, 2H). $^{13}$C NMR (151 MHz, MeOD) δ 157.43, 144.35, 144.13, 134.39, 118.58, 115.57, 93.64, 48.34, 40.79, 40.57, 28.57, 27.63, 27.32, 26.97. HRMS (ES+) calcd for ($C_{23}H_{33}N_4^+$) $[M]^+$ 365.26997, found 365.27094.

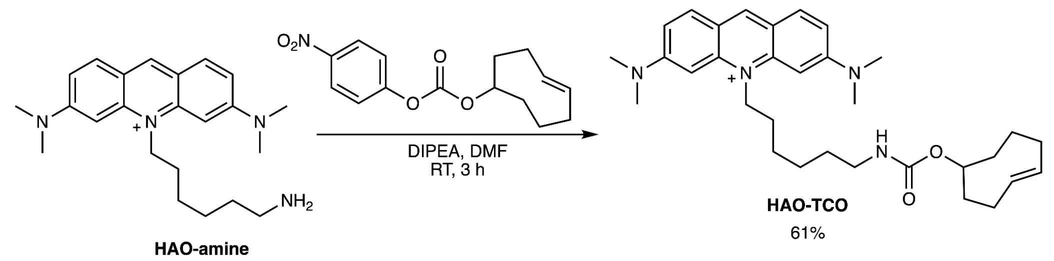

<u>(*E*)-10-(6-(((cyclooct-3-en-1-yloxy)carbonyl)amino)hexyl)-3,6-bis(dimethylamino)acridin-10-ium **(HAO-TCO)**</u>

To a 4 mL vial equipped with a stir bar was added **HAO-amine** (0.010 g, 0.021 mmol, 1.0 equiv.) dissolved in anhydrous DMF (0.209 mL) and DIPEA (11 µL, 0.063 mmol, 3.0 equiv.). The mixture was stirred and equilibrated for 5 min. TCO-PNB (8 mg, 0.027 mmol, 1.3 equiv.) was added to the reaction in one portion. The reaction was stirred at room temperature for 3 h—completion was confirmed by LCMS. Upon completion, the mixture was concentrated *in vacuo* and purified via reverse-phase HPLC (5-95% Solvent B over 25 min), yielding the title compound (8 mg, 61%) as an orange solid. $^1$H NMR (600 MHz, MeOD) δ 8.68 (s, 1H), 7.93 (d, $J$ = 9.3 Hz, 2H), 7.30 (dd, $J$ = 9.3, 2.2 Hz, 2H), 6.70 (d, $J$ = 2.3 Hz, 2H), 5.71 – 5.52 (m, 1H), 5.44 (ddd, $J$ = 15.7, 11.1, 3.6 Hz, 1H), 4.70 (d, $J$ = 16.7 Hz, 1H), 4.29 (d, $J$ = 8.3 Hz, 0H), 3.34 (s, 11H), 3.14 – 3.04 (m, 2H), 2.29 (qd, $J$ = 10.9, 4.8 Hz, 2H), 2.07 – 1.98 (m, 3H), 1.98 – 1.90 (m, 3H), 1.85 (s, 1H), 1.74 – 1.63 (m, 5H), 1.60 – 1.49 (m, 6H). $^{13}$C NMR (151 MHz, MeOD) δ 157.43, 144.35, 144.13, 134.39, 118.58, 115.57, 93.64, 48.34, 40.79, 40.57, 28.57, 27.63, 27.32, 26.97. HRMS (ES+) calcd for ($C_{32}H_{45}N_4O_2$+) $[M]^+$ 517.3538, found 517.34403.

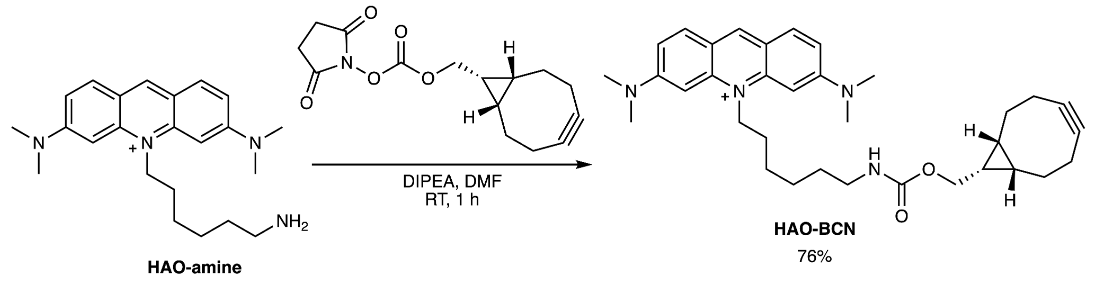

<u>10-(6-(((((1*R*,8*S*,9*s*)-bicyclo[6.1.0]non-4-yn-9-yl)methoxy)carbonyl)amino)hexyl)-3,6-bis(dimethylamino)acridin-10-ium **(HAO-BCN)**</u>

To a 4 mL vial equipped with a stir bar was added **HAO-amine** (0.010 g, 0.021 mmol, 1.0 equiv.) dissolved in anhydrous DMF (0.209 mL) and DIPEA (11 µL, 0.063 mmol, 3.0 equiv.). The mixture was stirred and equilibrated for 5 min. BCN-NHS ester (8 mg, 0.027 mmol, 1.3 equiv.) was added to the reaction in one portion. The reaction was stirred at room temperature for 1 h—completion was confirmed by LCMS. Upon completion, the mixture was concentrated *in vacuo* and purified via reverse-phase HPLC (5-95% acetonitrile gradient in water; **No TFA added** over 25 min), yielding the title compound (10 mg, 76%) as an orange solid. $^{1}H$ NMR (600 MHz, MeOD) δ 8.68 (s, 1H), 7.93 (d, *J* = 9.2 Hz, 2H), 7.32 – 7.27 (m, 2H), 6.70 (s, 2H), 4.76 – 4.63 (m, 2H), 4.12 (d, *J* = 8.2 Hz, 2H), 3.35 (s, 1H), 3.34 (s, 11H), 3.13 (q, *J* = 8.4 Hz, 2H), 2.91 (d, *J* = 13.2 Hz, 1H), 2.33 – 2.13 (m, 5H), 2.10 (d, *J* = 15.4 Hz, 2H), 2.02 (p, *J* = 7.9 Hz, 2H), 1.69 (p, *J* = 7.5 Hz, 2H), 1.61 – 1.51 (m, 4H), 1.41 – 1.24 (m, 2H), 0.90 (t, *J* = 9.6 Hz, 2H). $^{13}C$ NMR (151 MHz, MeOD) δ 156.04, 142.93, 142.76, 132.98, 117.20, 114.20, 98.08, 92.24, 62.15, 39.91, 39.42, 29.42, 28.76, 26.08, 25.91, 25.49, 20.48, 19.96, 17.56. HRMS (ES+) calcd for ($C_{34}H_{45}N_4O_2^{+}$) $[M]^{+}$ 541.3538, found 541.34378.

### Synthesis of Hoechst derivatives

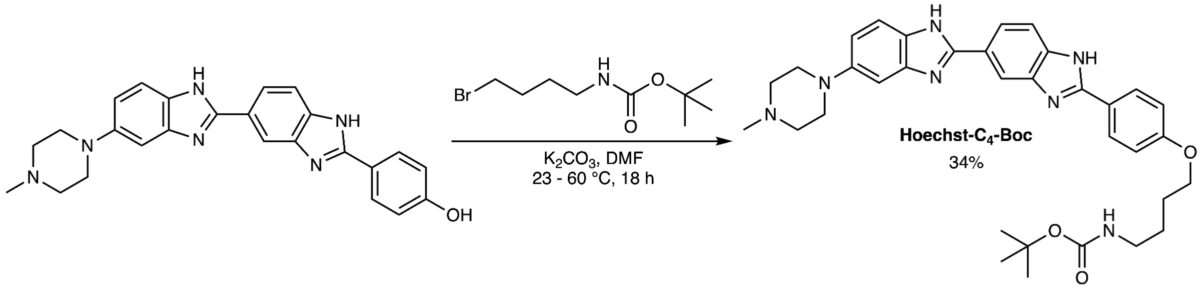

<u>*tert*-butyl (4-(4-(5-(4-methylpiperazin-1-yl)-1*H*,1'*H*-[2,5'-bibenzo[*d*]imidazol]-2'-yl)phenoxy)butyl)carbamate **(Hoechst-C$_4$-Boc)**</u>

**Hoechst-C$_4$-Boc** was prepared according to the procedure published in Lukinavičius, G. *et al*[4]. Hoechst 33258 hydrochloride salt was treated to isolate Hoechst free base—the free base was used to prepare **Hoechst-C$_4$-Boc**.

To a 5 mL microwave vial equipped with a stir bar was added Hoechst free base (0.100 g, 0.236 mmol, 1.0 equiv.) and $K_2CO_3$ (0.088 g, 0.636 mmol, 2.7 equiv.). The vial was temporarily sealed (suba seal septa), evacuated, and filled with $N_2$. Anhydrous DMF (1.18 mL, 0.2 M) was added, and the mixture was stirred for 30 min to equilibrate. 4-(Boc-amino)butyl bromide (0.077 g, 0.306 mmol, 1.3 equiv.) was added to the vial in portions. The vial was then sealed and purged with nitrogen. The reaction was heated to 60 °C and stirred for 18 h. Reaction completion was confirmed by LCMS. Upon completion, the crude was dissolved in 5:5:1 acetonitrile:water:acetic acid, then purified by reverse-phase HPLC (5-95% Solvent B over 15 min), affording the title compound as a TFA salt (75 mg, 34%).$^1$H NMR (500 MHz, MeOD) δ 8.38 – 8.34 (m, 1H), 8.11 – 8.02 (m, 3H), 7.91 (d, *J* = 8.6 Hz, 1H), 7.69 (d, *J* = 9.0 Hz, 1H), 7.36 (dd, *J* = 9.1, 2.3 Hz, 1H), 7.29 (d, *J* = 2.3 Hz, 1H), 7.12 (d, *J* = 8.6 Hz, 2H), 4.05 (t, *J* = 6.3 Hz, 2H), 3.94 (d, *J* = 13.1 Hz, 2H), 3.69 (d, *J* = 11.7 Hz, 2H), 3.34 (d, *J* = 9.6 Hz, 2H), 3.21 (d, *J* = 13.2 Hz, 2H), 3.12 (t, *J* = 6.9 Hz, 2H), 3.01 (s, 3H), 1.81 (dq, *J* = 12.0, 6.6 Hz, 2H), 1.65 (p, *J* = 7.1 Hz, 2H), 1.45 (s, 9H). $^{13}$C NMR (126 MHz, MeOD) δ 164.70, 162.30, 162.00, 158.62, 154.80, 150.74, 149.70, 138.49, 136.32, 134.60, 130.88, 128.07, 125.07, 120.53, 119.53, 118.85, 117.42, 116.75, 116.45, 115.65, 114.93, 100.91, 79.94, 69.34, 54.57, 49.63, 49.45, 49.28, 48.38, 43.57, 41.00, 28.80, 27.59, 27.46. HRMS (ES+) calcd for ($C_{34}H_{42}N_7O_3$) $[M+H]^+$ 596.33436, found 596.33423.

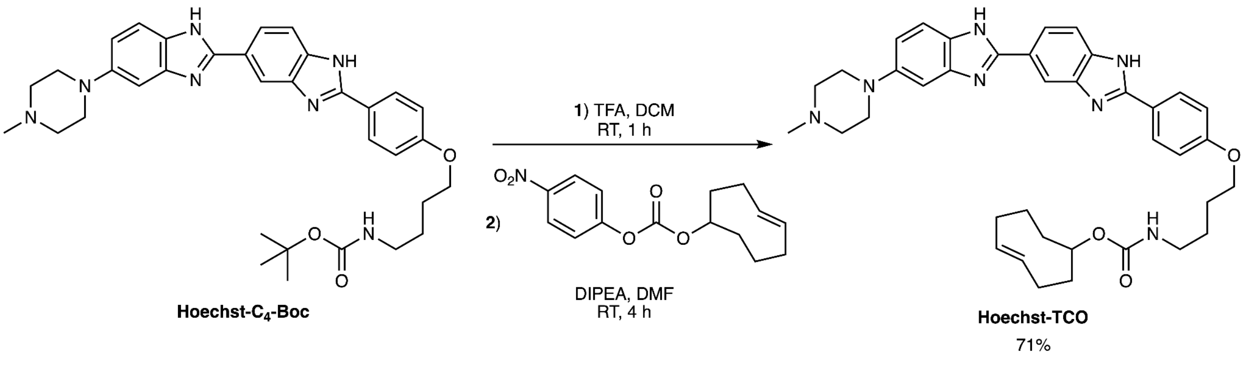

(*E*)-cyclooct-4-en-1-yl (4-(4-(5-(4-methylpiperazin-1-yl)-1*H*,1'*H*-[2,5'-bibenzo[*d*]imidazol]-2'-yl)phenoxy)butyl)carbamate **(Hoechst-TCO)**

To a 4 mL vial equipped with a stir bar was added **Hoechst-C$_4$-Boc** (0.020 g, 0.021 mmol) dissolved in DCM (0.213 mL) and TFA (0.020 mL, 0.267 mmol, 12.5 equiv.). The reaction mixture was stirred at room temperature for 1 h (complete deprotection confirmed by LCMS). Upon completion, the reaction mixture was concentrated *in vacuo*. The deprotected amine intermediate was dissolved in DMF (0.213 mL, 0.1 M), DIPEA (46 µL, 0.267 mmol, 12.5 equiv.) was added, and the solution was stirred for 5 min to equilibrate. TCO-PNB ester (7 mg, 0.026 mmol, 1.2 equiv.) was dissolved in DMF (0.213 mL) and added dropwise. The reaction was stirred at room temperature and monitored by LCMS until completion (4 h). The mixture was concentrated *in vacuo*, dissolved in 50:50 water/acetonitrile containing 0.1% formic acid, and purified via reverse-phase HPLC (5-95% Solvent B over 15 min) to yield the title compound as a yellow solid (15 mg, 71%). $^1$H NMR (600 MHz, MeOD) δ 8.42 – 8.39 (m, 1H), 8.13 – 8.08 (m, 2H), 8.06 (dd, *J* = 8.5, 1.8 Hz, 1H), 7.91 (dd, *J* = 8.5, 0.7 Hz, 1H), 7.72 (d, *J* = 9.0 Hz, 1H), 7.39 (dd, *J* = 9.0, 2.3 Hz, 1H), 7.31 (d, *J* = 2.2 Hz, 1H), 7.19 – 7.14 (m, 2H), 5.60 (ddd, *J* = 21.1, 10.1, 5.3 Hz, 1H), 5.47 (ddd, *J* = 15.9, 11.0, 3.6 Hz, 1H), 4.34 – 4.28 (m, 1H), 4.11 (t, *J* = 6.3 Hz, 2H), 3.95 (s, 2H), 3.69 (s, 2H), 3.35 (s, 2H), 3.17 (dt, *J* = 8.2, 4.1 Hz, 2H), 3.02 (s, 3H), 2.33 (dh, *J* = 15.5, 4.9 Hz, 3H), 1.99 (s, 1H), 2.00 – 1.88 (m, 2H), 1.84 (dq, *J* = 8.9, 6.4 Hz, 2H), 1.78 – 1.64 (m, 4H), 1.59 (ddd, *J* = 15.8, 12.1, 5.7 Hz, 1H). $^{13}$C NMR (126 MHz, MeOD) δ 164.70, 162.30, 162.00, 158.62, 154.80, 150.74, 149.70, 138.49, 136.32, 134.60, 130.88, 128.07, 125.07, 120.53, 119.53, 118.85, 117.42, 116.75, 116.45, 115.65, 114.93, 100.91, 79.94, 69.34, 54.57, 49.63, 49.45, 49.28, 48.38, 43.57, 41.00, 28.80, 27.59, 27.46. . HRMS (ES+) calcd for ($C_{38}H_{45}N_7O_3$) $[M+H]^+$ 648.36566, found 648.35196.

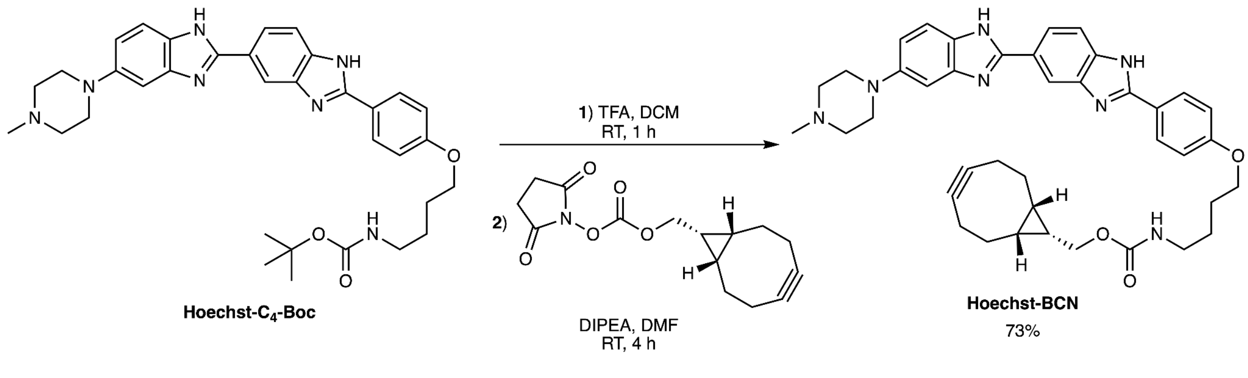

((1*R*,8*S*,9*s*)-bicyclo[6.1.0]non-4-yn-9-yl)methyl (4-(4-(5-(4-methylpiperazin-1-yl)-1*H*,1'*H*-[2,5'-bibenzo[*d*]imidazol]-2'-yl)phenoxy)butyl)carbamate **(Hoechst-BCN)**

To a 4 mL vial equipped with a stir bar was added **Hoechst-C$_4$-Boc** (0.022 g, 0.023 mmol) dissolved in DCM (0.250 mL) and TFA (0.022 mL, 0.293 mmol, 12.5 equiv.). The reaction mixture was stirred at room temperature for 1 h (complete deprotection confirmed by LCMS).

Upon completion, the reaction mixture was concentrated *in vacuo*. The deprotected amine intermediate was dissolved in DMF (0.235 mL, 0.1 M), DIPEA (51 μL, 0.293 mmol, 12.5 equiv.) was added, and the solution was stirred for 5 min to equilibrate. BCN-NHS ester (8 mg, 0.028 mmol, 1.2 equiv.) was dissolved in DMF (0.235 mL) and added dropwise. The reaction was stirred at room temperature and monitored by LCMS until completion (4 h). The mixture was concentrated *in vacuo*, dissolved in 50:50 water/acetonitrile containing 0.1% formic acid, and purified via reverse-phase HPLC (5-95% Solvent B over 20 min) to yield the title compound as a yellow solid (18 mg, 73%). $^{1}$H NMR (600 MHz, MeOD) δ 8.34 (d, *J* = 1.7 Hz, 1H), 8.09 – 8.02 (m, 3H), 7.88 (d, *J* = 8.5 Hz, 1H), 7.68 (d, *J* = 9.0 Hz, 1H), 7.35 (dd, *J* = 9.0, 2.2 Hz, 1H), 7.28 (d, *J* = 2.2 Hz, 1H), 7.11 (d, *J* = 8.5 Hz, 2H), 4.14 (d, *J* = 8.1 Hz, 2H), 4.06 (t, *J* = 6.4 Hz, 2H), 3.99 – 3.86 (m, 2H), 3.68 (s, 2H), 3.64 (d, *J* = 7.7 Hz, 2H), 3.43 – 3.32 (m, 2H), 3.19 (t, *J* = 6.9 Hz, 4H), 3.01 (s, 3H), 2.29 – 2.18 (m, 7H), 2.14 (ddt, *J* = 15.9, 4.5, 2.2 Hz, 4H), 1.86 – 1.77 (m, 3H), 1.72 – 1.64 (m, 3H), 1.64 – 1.53 (m, 4H), 1.38 (p, *J* = 8.7 Hz, 1H), 1.27 (tt, *J* = 9.2, 7.7 Hz, 1H), 0.97 – 0.84 (m, 3H). $^{13}$C NMR (126 MHz, MeOD) δ 164.70, 162.30, 162.00, 158.62, 154.80, 150.74, 149.70, 138.49, 136.32, 134.60, 130.88, 128.07, 125.07, 120.53, 119.53, 118.85, 117.42, 116.75, 116.45, 115.65, 114.93, 100.91, 79.94, 69.34, 54.57, 49.63, 49.45, 49.28, 48.38, 43.57, 41.00, 28.80, 27.59, 27.46. HRMS (ES+) calcd for ($C_{40}H_{45}N_7O_3$) $[M+H]^+$ 672.36566, found 672.35274.

**Synthesis of Docetaxel Ligands**

Synthesis of Docetaxel ligands was adapted from Beliu, G. *et al*[5].

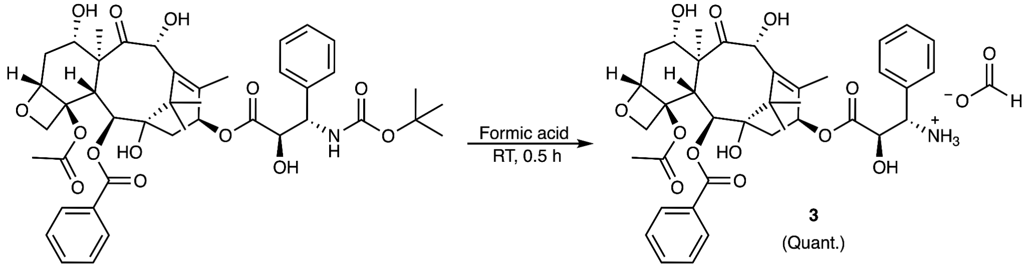

### 3’-Aminodocetaxel

No inert conditions required. To a 4 mL vial equipped with a stir bar was added Docetaxel (100 mg, 0.124 mmol) dissolved in formic acid (2 mL, 0.060 M). The reaction was stirred at room temperature for 0.5 h and monitored by LCMS (complete deprotection confirmed by LCMS). The reaction mixture was concentrated *in vacuo*, redissolved in water, and lyophilized. The title compound was isolated as a white solid (quantitative yield) and used without further purification.

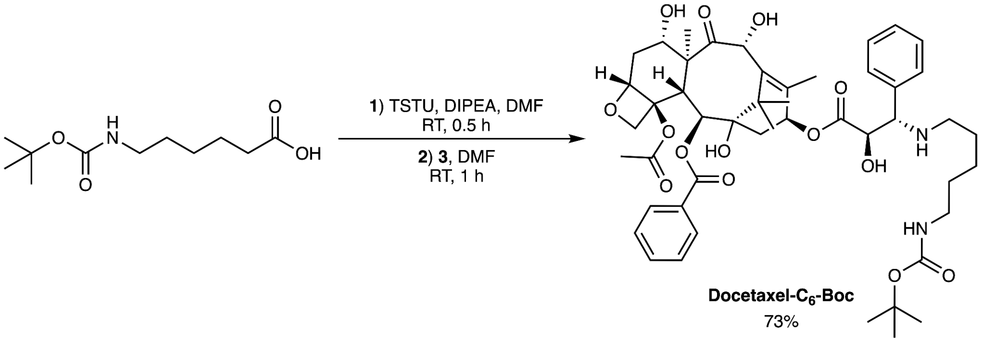

### Docetaxel-$C_6$-Boc

To a 5 mL microwave vial equipped with a stir bar was added 6-((tert-butoxycarbonyl)amino)hexanoic acid (**2**) (0.019 g, 0.084 mmol, 1.2 equiv.), TSTU (0.027 g, 0.091 mmol, 1.3 equiv.), DMF (0.702 mL, 0.100 M), and DIPEA (53 µL, 0.302 mmol, 4.3 equiv.). The reaction mixture was stirred at room temperature for 0.5 h. Formation of the NHS-ester intermediate was confirmed by LCMS. To a separate 4 mL vial was added **3’-aminodocetaxel** (0.053 g, 0.070 mmol, 1.0 equiv.) dissolved in DMF (250 µL). The **3’-aminodocetaxel** solution was then added dropwise to the NHS-ester mixture and stirred at room temperature for 1 h (complete conversion confirmed by LCMS). The reaction mixture was concentrated *in vacuo*, redissolved in 50:50 acetonitrile/water containing 0.1% formic acid, and purified by reverse-phase HPLC (5–95% Solvent B over 25 min), providing the title compound (43 mg, 71% yield) as a white solid. $^1$H NMR (500 MHz, MeOD) δ 8.14 – 8.06 (m, 2H), 7.70 – 7.62 (m, 1H), 7.60 – 7.53 (m, 2H), 7.45 – 7.35 (m, 4H), 7.28 (qt, *J* = 6.5, 1.7 Hz, 1H), 6.16 (td, *J* = 9.1, 1.7 Hz, 1H), 5.65 (d, *J* = 7.2 Hz, 1H), 5.52 – 5.42 (m, 1H), 5.27 (s, 1H), 4.99 (dd, *J* = 9.7, 2.2 Hz, 1H), 4.59

(dd, $J$ = 7.7, 4.5 Hz, 1H), 4.26 – 4.18 (m, 2H), 4.19 (d, $J$ = 8.4 Hz, 1H), 3.87 (d, $J$ = 7.2 Hz, 1H), 2.98 (t, $J$ = 7.1 Hz, 2H), 2.44 (ddd, $J$ = 14.4, 9.7, 6.5 Hz, 1H), 2.33 (d, $J$ = 4.1 Hz, 3H), 2.32 – 2.14 (m, 3H), 2.05 – 1.93 (m, 1H), 1.90 (t, $J$ = 1.8 Hz, 3H), 1.83 (ddd, $J$ = 13.9, 11.3, 2.3 Hz, 1H), 1.70 (s, 3H), 1.68 – 1.53 (m, 3H), 1.44 (d, $J$ = 7.4 Hz, 1H), 1.42 (s, 10H), 1.31 (dtt, $J$ = 14.9, 7.2, 4.1 Hz, 2H), 1.19 (s, 3H), 1.13 (d, $J$ = 3.8 Hz, 3H). $^{13}$C NMR (126 MHz, MeOD) δ 209.67, 174.50, 173.03, 170.40, 166.27, 159.25, 158.94, 157.12, 138.73, 137.81, 136.58, 133.13, 130.03, 129.79, 128.29, 128.25, 127.47, 127.01, 84.51, 80.93, 78.41, 77.68, 76.19, 75.02, 74.24, 73.45, 71.26, 71.13, 57.46, 55.46, 48.21, 48.04, 47.87, 46.46, 43.13, 39.76, 39.10, 36.09, 35.49, 35.41, 29.22, 27.38, 26.01, 25.67, 25.47, 25.34, 21.81, 20.26, 13.00, 9.05. HRMS (ES+) calcd for ($C_{49}H_{64}N_2NaO_{15}$) $[M+Na]^+$ 943.41989, found 943.42020.

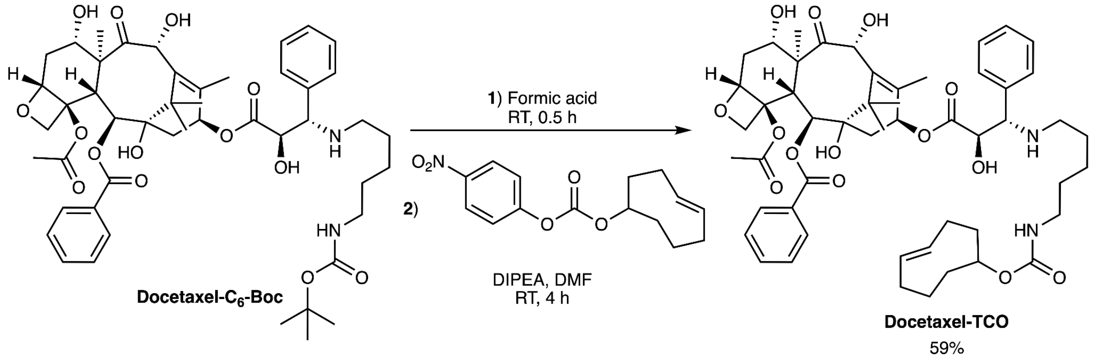

### Docetaxel-TCO

To a 4 mL vial equipped with a stir bar was added **Docetaxel-C$_6$-Boc** (0.010 g (mass calc. for TFA salt), 0.010 mmol, 1 equiv.) dissolved in formic acid (0.5 mL). he reaction was stirred at room temperature for 0.5 h (complete deprotection confirmed by LCMS). The crude mixture was concentrated and dried overnight *in vacuo*. The deprotected amine intermediate was dissolved in DMF (97 µL, 0.1 M), DIPEA (21 µL, 0.121 mmol, 12.5 equiv.) was added, and the solution was equilibrated for 15 min. TCO-PNB ester (3 mg, 0.012 mmol, 1.2 equiv.) was dissolved in DMF (97 µL) and added to the reaction mixture dropwise. The reaction was stirred at room temperature for 4 h (complete conversion confirmed by LCMS). The reaction mixture was concentrated *in vacuo*. The crude was resuspended in 1:1:1 methanol/$H_2O$/acetonitrile, and purified via reverse phase HPLC (5-95% Solvent B over 25 min), yielding the title compound (5.6 mg, 59%) as a white solid. $^1$H NMR (600 MHz, MeOD) δ 8.14 – 8.09 (m, 2H), 7.69 – 7.63 (m, 1H), 7.56 (t, $J$ = 7.8 Hz, 2H), 7.44 – 7.34 (m, 4H), 7.28 (tt, $J$ = 6.9, 1.7 Hz, 1H), 6.16 (t, $J$ = 9.0 Hz, 1H), 5.65 (d, $J$ = 7.2 Hz, 1H), 5.58 (ddd, $J$ = 15.3, 9.9, 4.9 Hz, 1H), 5.51 – 5.42 (m, 2H), 5.27 (s, 1H), 4.99 (dd, $J$ = 9.7, 2.2 Hz, 1H), 4.58 (d, $J$ = 4.6 Hz, 1H), 4.28 (s, 1H), 4.25 – 4.17 (m, 3H), 3.88 (d, $J$ = 7.2 Hz, 1H), 3.00 (dt, $J$ = 10.8, 6.6 Hz, 2H), 2.44 (ddd, $J$ = 14.4, 9.7, 6.5 Hz, 1H), 2.34 (d, $J$ = 0.8 Hz, 3H), 2.33 – 2.21 (m, 7H), 2.05 – 1.96 (m, 2H), 1.96 – 1.88 (m, 5H), 1.83 (ddd, $J$ = 13.9, 11.3, 2.3 Hz, 1H), 1.70 (s, 3H), 1.68 – 1.62 (m, 1H), 1.59 (dd, $J$ = 16.2, 8.9 Hz, 3H), 1.48 – 1.39 (m, 2H), 1.35 – 1.26 (m, 2H), 1.19 (s, 3H), 1.13 (s, 3H). $^{13}$C NMR (151 MHz, MeOD) δ 209.73, 170.42, 166.31, 137.82, 136.62, 134.68, 133.14, 132.37, 130.06, 129.80, 128.30, 128.26, 127.47, 127.02, 84.54, 80.98, 80.15, 77.71, 76.23, 75.06, 74.26, 73.44, 71.28, 71.18, 57.50, 55.44, 46.49, 43.16, 40.83, 38.24, 36.12, 35.47, 33.77, 32.09, 30.71, 29.24,

25.96, 25.72, 25.29, 21.83, 20.29, 13.01, 9.07. HRMS (ES+) calcd for ($C_{53}H_{69}N_2O_{15}$) $[M+H]^+$ 973.46924, found 973.45095.

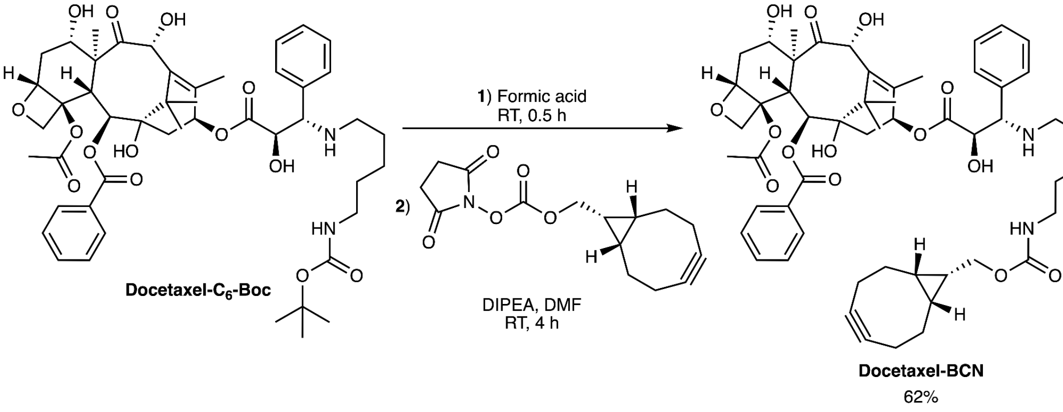

### Docetaxel-BCN

To a 4 mL vial equipped with a stir bar was added **Docetaxel-C$_6$-Boc** (0.010 g (mass calc. for TFA salt), 0.010 mmol, 1 equiv.) dissolved in formic acid (0.5 mL). The reaction was stirred at room temperature for 0.5 h (complete deprotection confirmed by LCMS). The crude mixture was concentrated and dried overnight *in vacuo*. The deprotected amine intermediate was dissolved in DMF (97 µL, 0.1 M), DIPEA (21 µL, 0.121 mmol, 12.5 equiv.) was added, and the solution was equilibrated for 15 min. BCN-NHS ester (3 mg, 0.012 mmol, 1.2 equiv.) was dissolved in DMF (97 µL) and added to the reaction mixture dropwise. The reaction was stirred at room temperature for 4 h. (complete conversion confirmed by LCMS). The reaction mixture was concentrated *in vacuo*. The crude was resuspended in 1:1:1 methanol/$H_2O$/acetonitrile, and purified via reverse phase HPLC (5-95% Solvent B over 25 min), yielding the title compound (6.0 mg, 59%) as a white solid. $^1H$ NMR (600 MHz, MeOD) δ 8.14 – 8.09 (m, 2H), 7.66 (ddt, *J* = 7.9, 6.9, 1.3 Hz, 1H), 7.60 – 7.53 (m, 2H), 7.44 – 7.36 (m, 4H), 7.31 – 7.25 (m, 1H), 6.16 (t, *J* = 8.9 Hz, 1H), 5.65 (d, *J* = 7.2 Hz, 1H), 5.46 (d, *J* = 4.6 Hz, 1H), 5.27 (s, 1H), 4.99 (dd, *J* = 9.8, 2.2 Hz, 1H), 4.59 (d, *J* = 4.5 Hz, 1H), 4.25 – 4.17 (m, 3H), 4.11 (d, *J* = 8.1 Hz, 2H), 3.88 (d, *J* = 7.1 Hz, 1H), 3.05 (t, *J* = 7.0 Hz, 2H), 2.44 (ddd, *J* = 14.4, 9.7, 6.5 Hz, 1H), 2.34 (s, 3H), 2.33 – 2.24 (m, 4H), 2.22 (ddt, *J* = 14.0, 10.3, 2.1 Hz, 5H), 2.20 – 2.13 (m, 3H), 2.08 – 1.99 (m, 2H), 1.90 (d, *J* = 1.4 Hz, 3H), 1.83 (ddd, *J* = 13.9, 11.3, 2.4 Hz, 1H), 1.70 (s, 3H), 1.62 (p, *J* = 7.4 Hz, 4H), 1.49 – 1.42 (m, 2H), 1.42 – 1.28 (m, 3H), 1.19 (s, 3H), 1.13 (s, 3H), 0.95 – 0.89 (m, 2H). $^{13}C$ NMR (151 MHz, MeOD) δ 211.12, 174.45, 171.83, 167.71, 140.18, 139.23, 138.03, 134.54, 131.47, 131.21, 129.70, 129.66, 128.87, 128.42, 99.53, 85.94, 82.39, 79.11, 77.63, 76.48, 75.67, 74.85, 72.68, 72.60, 63.57, 58.90, 56.85, 49.57, 47.90, 44.56, 41.53, 37.52, 36.89, 30.16, 27.36, 27.10, 26.70, 23.22, 21.93, 21.69, 21.38, 18.97, 14.41, 10.46. HRMS (ES+) calcd for ($C_{55}H_{69}N_2O_{15}^+$) $[M+H]^+$ 997.46924, found 997.44931.

#### Synthesis of HD555

HD555 was prepared according to the previously published procedure in Werther, P. *et al.*[6]

Characterization data matched with previously published data: $^1$H NMR (400 MHz, MeOD) δ 7.98 (dd, $J$ = 7.7, 1.2 Hz, 1H), 7.81 – 7.68 (m, 2H), 7.20 (d, $J$ = 9.5 Hz, 2H), 7.02 (dd, $J$ = 9.5, 2.5 Hz, 2H), 6.92 (d, $J$ = 2.5 Hz, 2H), 5.65 (s, 2H), 3.30 (s, 11H), 2.94 (s, 3H).

**References**

1. Singh, V., Wang, S. & Kool, E. T. Genetically Encoded Multispectral Labeling of Proteins with Polyfluorophores on a DNA Backbone. *J. Am. Chem. Soc.* **135**, 6184–6191 (2013).
2. Chen, W., Guo, C., Ding, H., Yang, X. & Zhang, K. Controlled Ring-Opening Polymerization of Macrocyclic Monomers Based on Ring-Opening/Ring-Closing Cascade Reaction. *J. Am. Chem. Soc.* jacs.3c10765 (2023) doi:10.1021/jacs.3c10765.
3. Zheng, S. *et al.* Long-term super-resolution inner mitochondrial membrane imaging with a lipid probe. *Nat. Chem. Biol.* **20**, 83–92 (2024).
4. Lukinavičius, G. *et al.* SiR–Hoechst is a far-red DNA stain for live-cell nanoscopy. *Nat. Commun.* **6**, 8497 (2015).
5. Beliu, G. *et al.* Bioorthogonal labeling with tetrazine-dyes for super-resolution microscopy. *Commun. Biol.* **2**, 261 (2019).
6. Werther, P. *et al.* Bio-orthogonal Red and Far-Red Fluorogenic Probes for Wash-Free Live-Cell and Super-resolution Microscopy. *ACS Cent. Sci.* **7**, 1561–1571 (2021).

**NMR Spectra**

[1]H - NMR (500 MHz, $CDCl_3$)

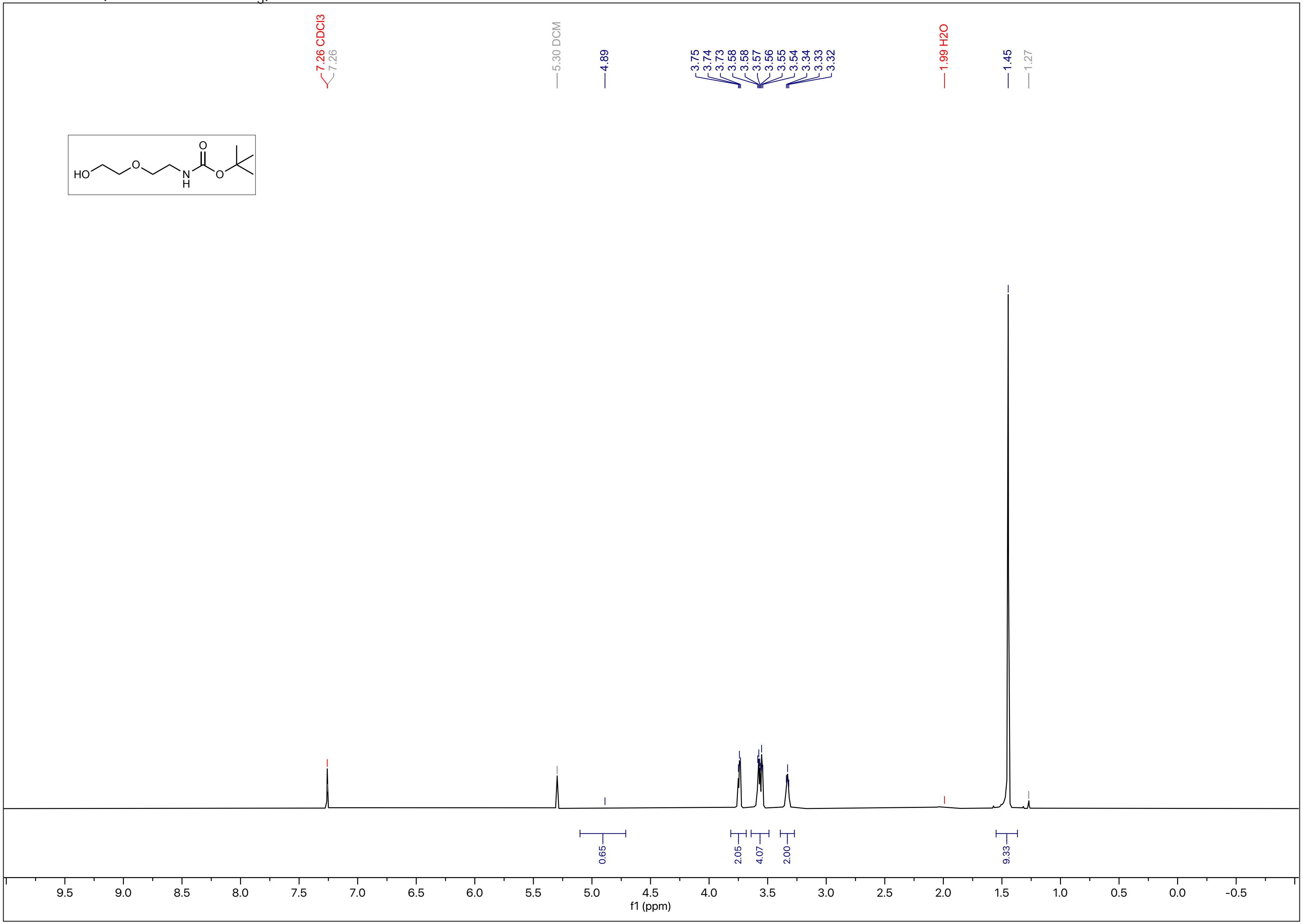

N-2

$^{13}C$ - NMR (126 MHz, $CDCl_3$)

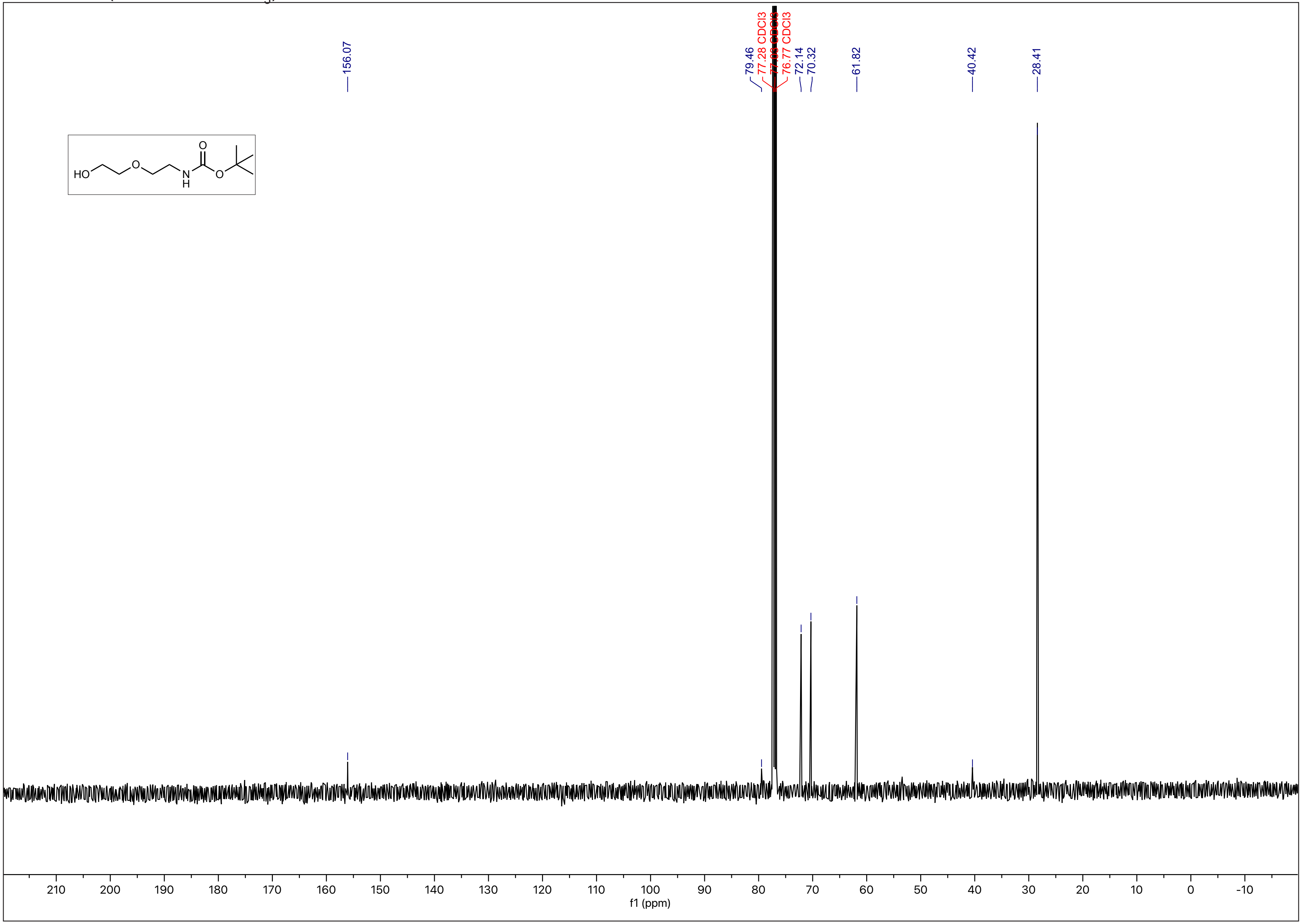

N-3

[1]H - NMR (500 MHz, $CDCl_3$)

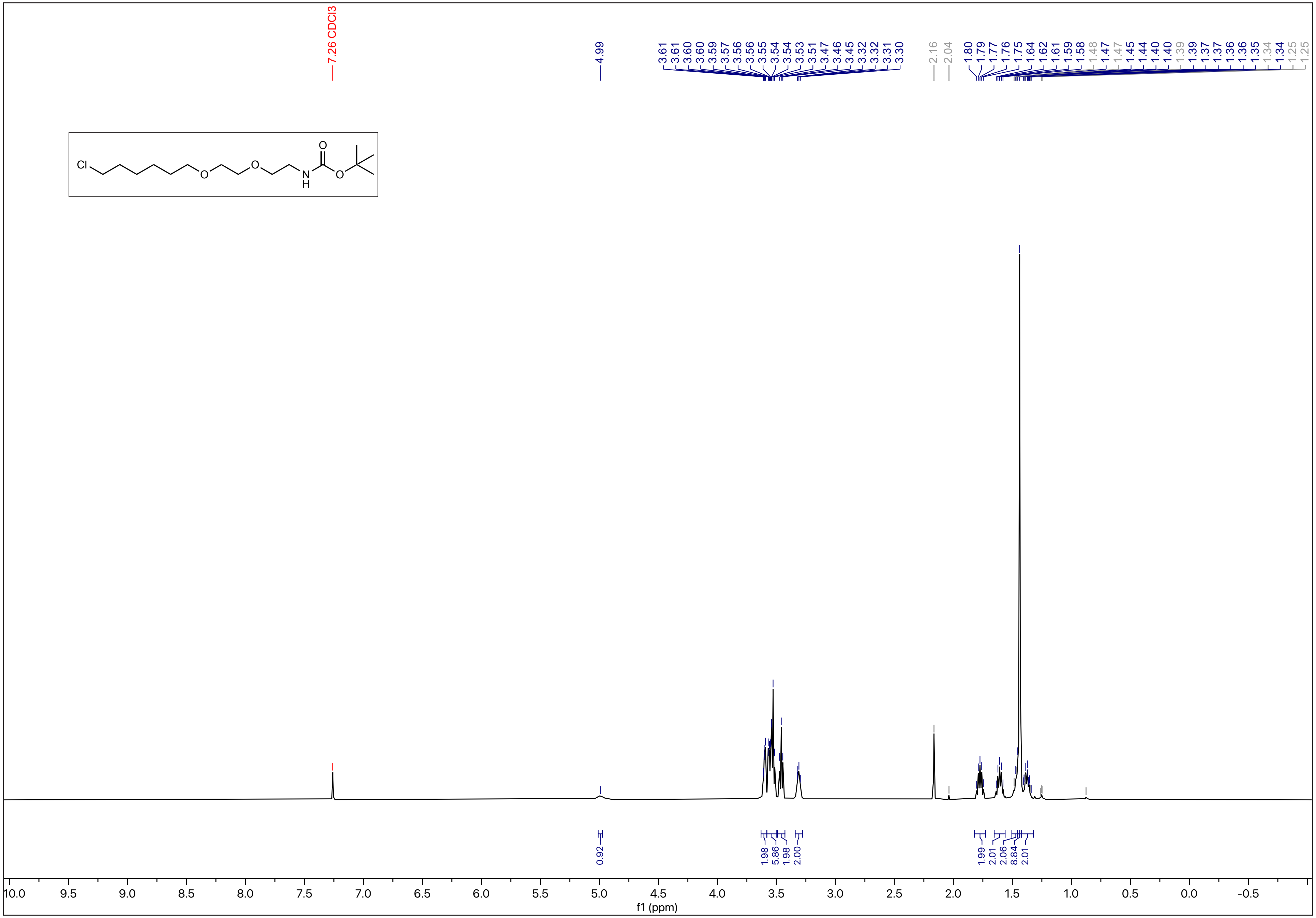

N-4

### $^{13}C$ - NMR (126 MHz, $CDCl_3$)

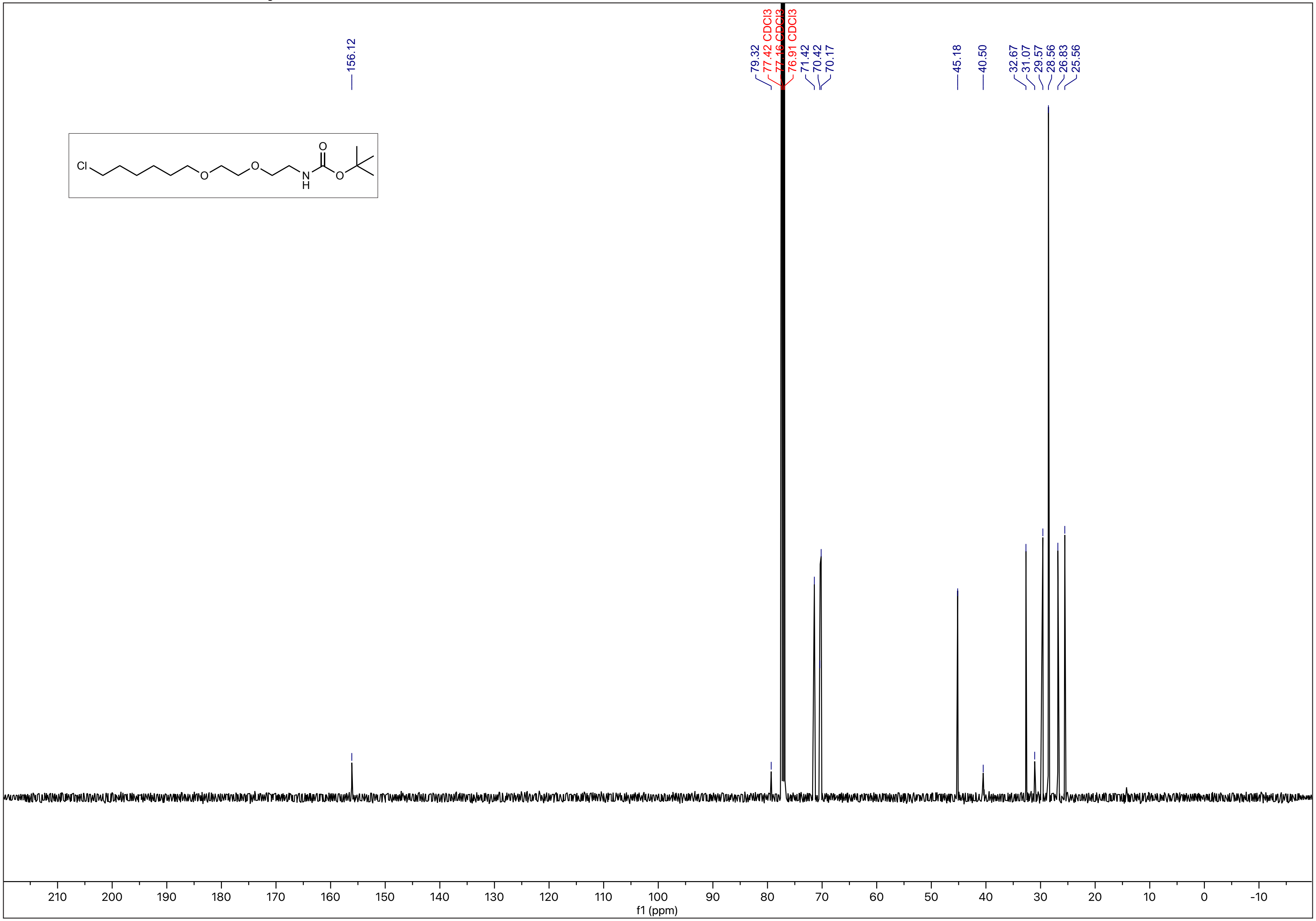

N-5

$^{1}H$ - NMR (400 MHz, $CDCl_3$)

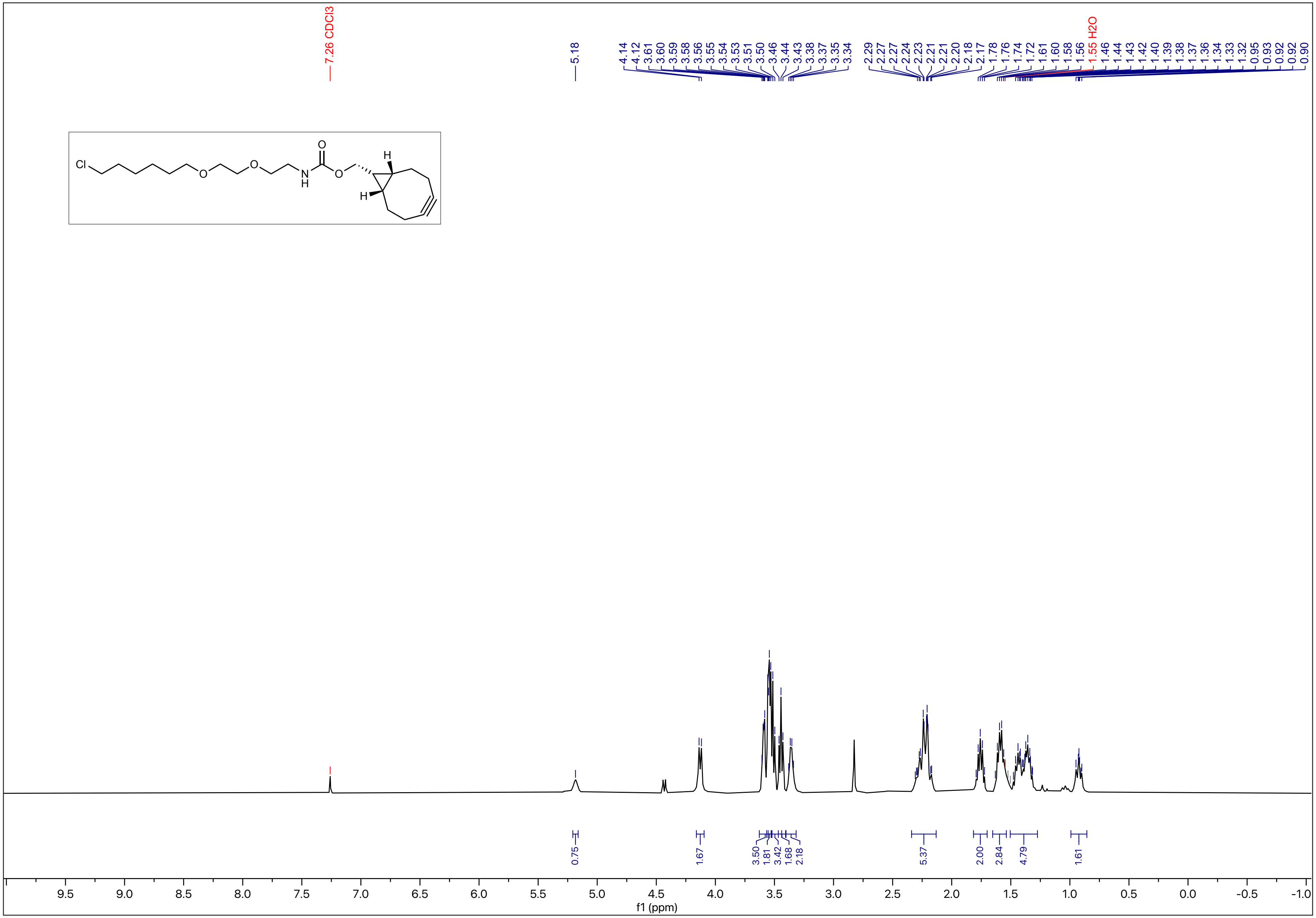

$^{13}C$ - NMR (101 MHz, $CDCl_3$)

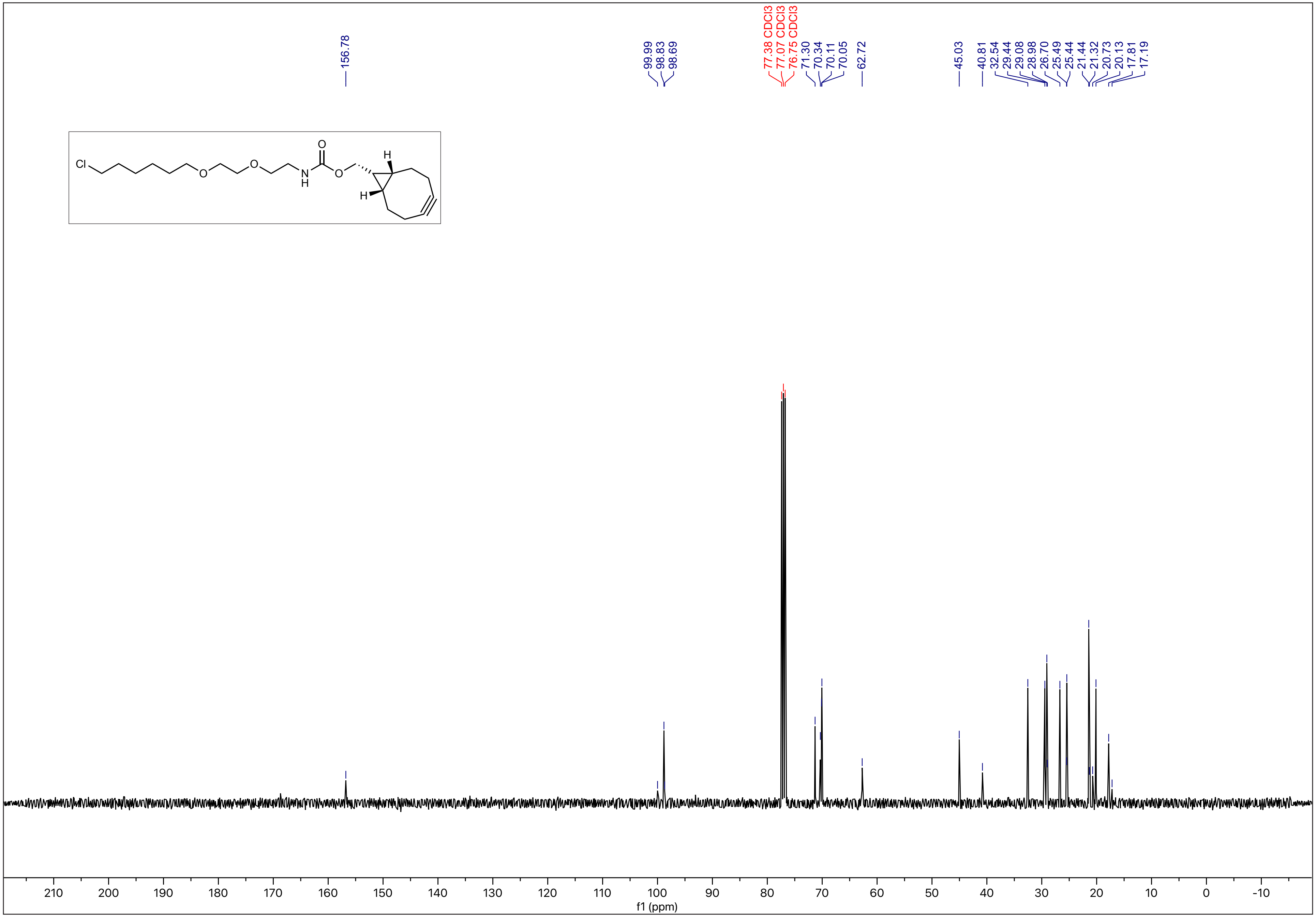

N-7

$^{1}H$ - NMR (500 MHz, $CDCl_3$)

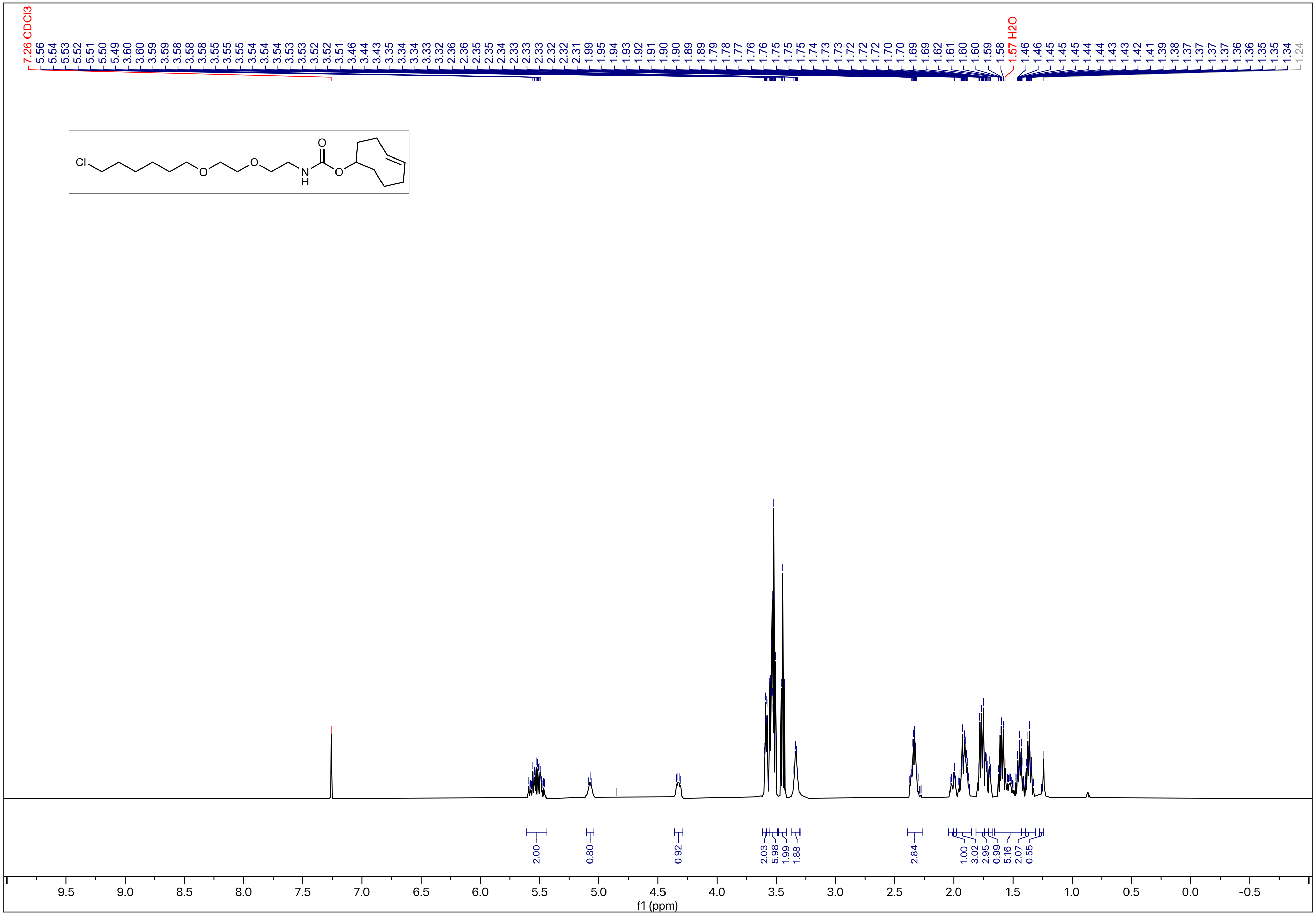

$^{13}C$ - NMR (126 MHz, $CDCl_3$)

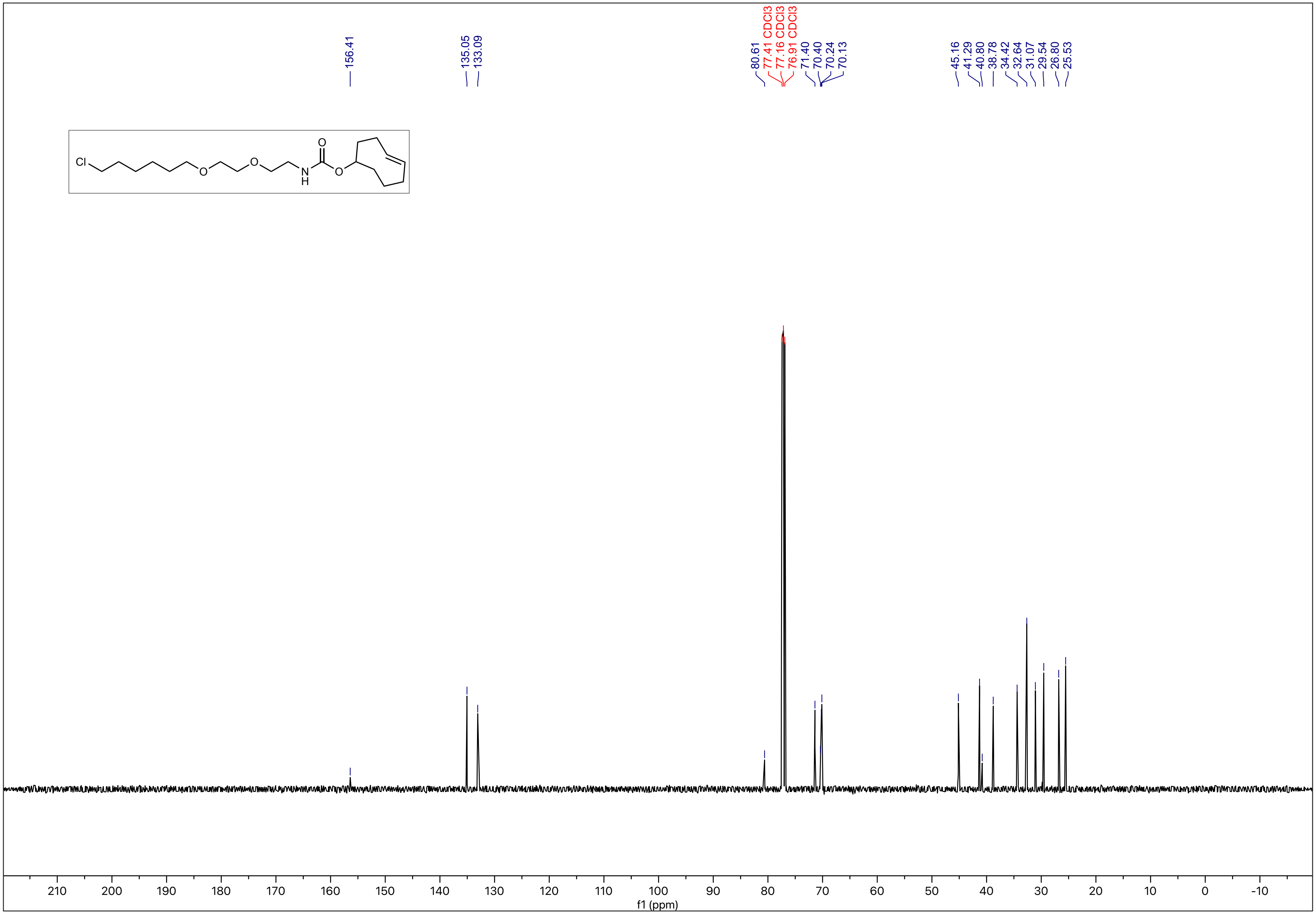

$^{1}H$ - NMR (500 MHz, MeOD)

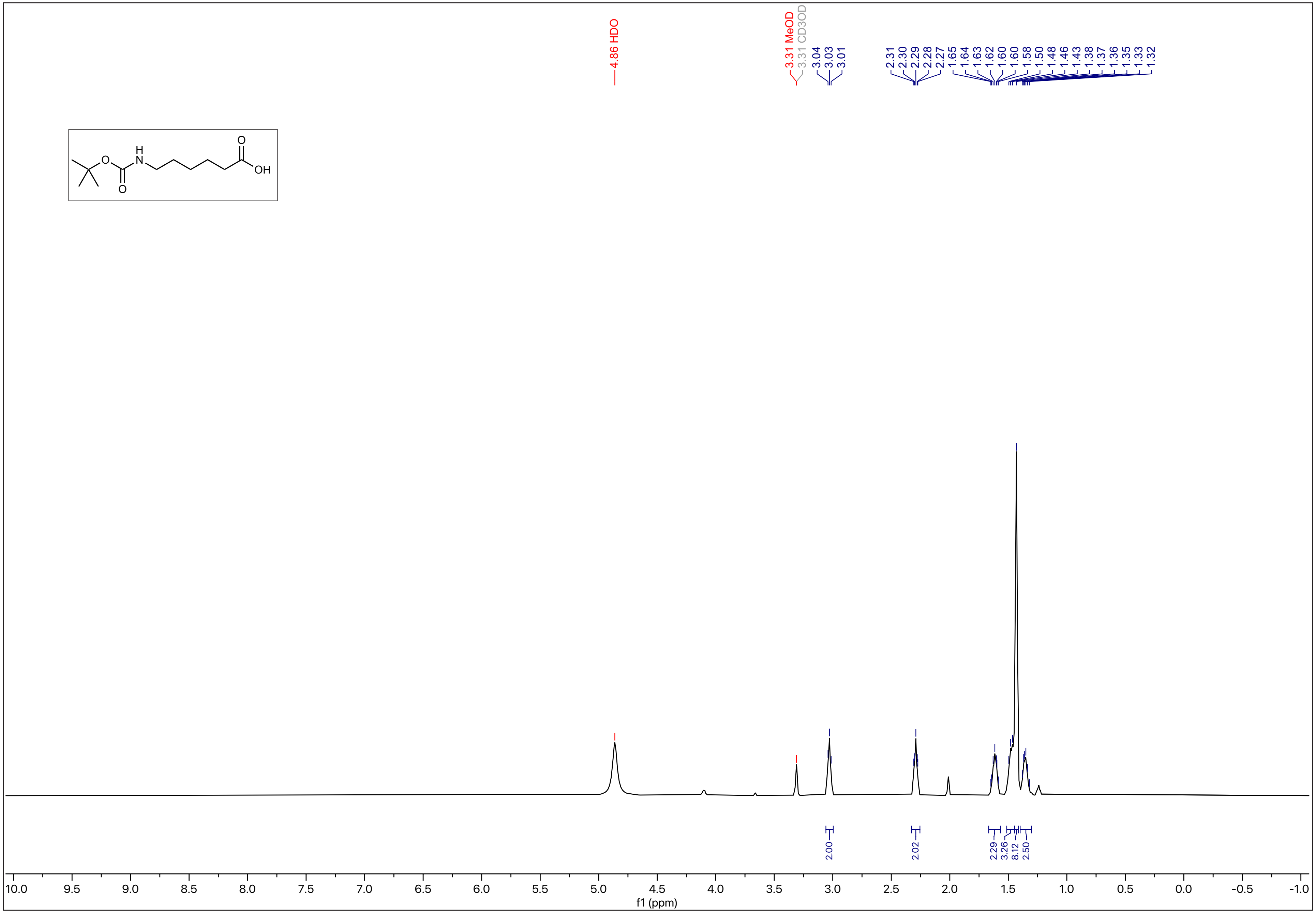

N-10

$^{13}C$ - NMR (126 MHz, MeOD)

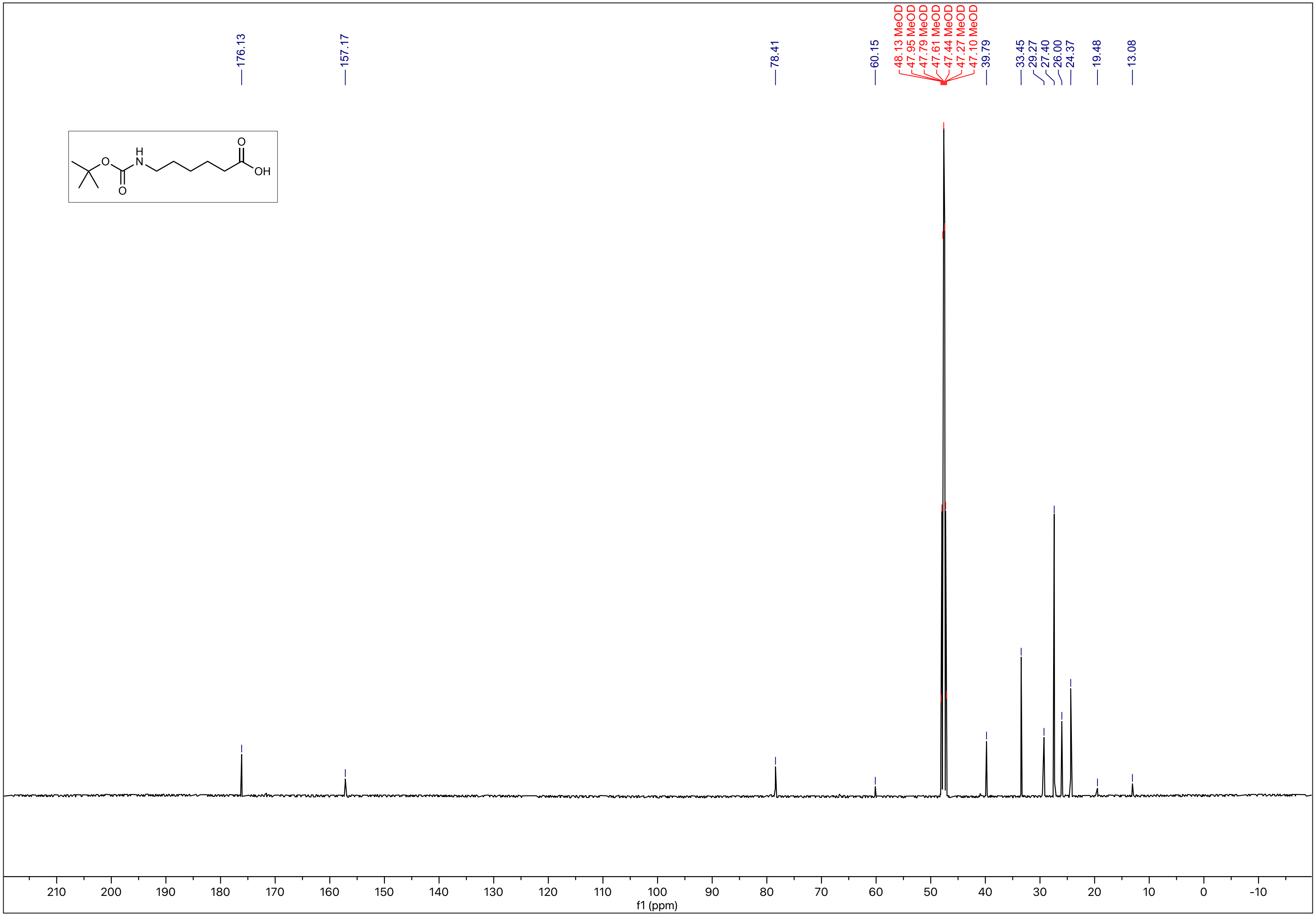

N-11

[1]H - NMR (500 MHz, MeOD)

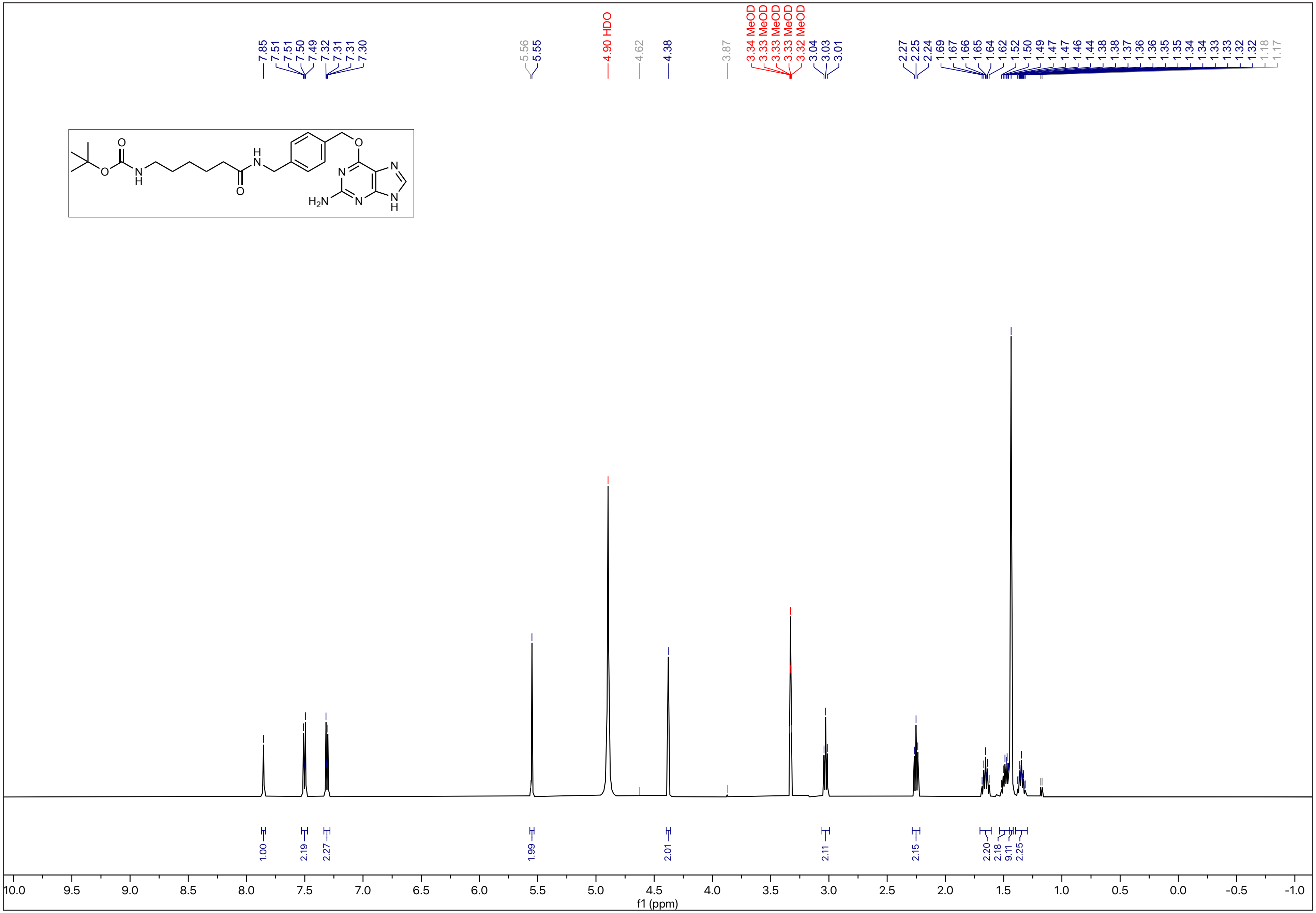

N-12

$^{13}C$ - NMR (126 MHz, MeOD)

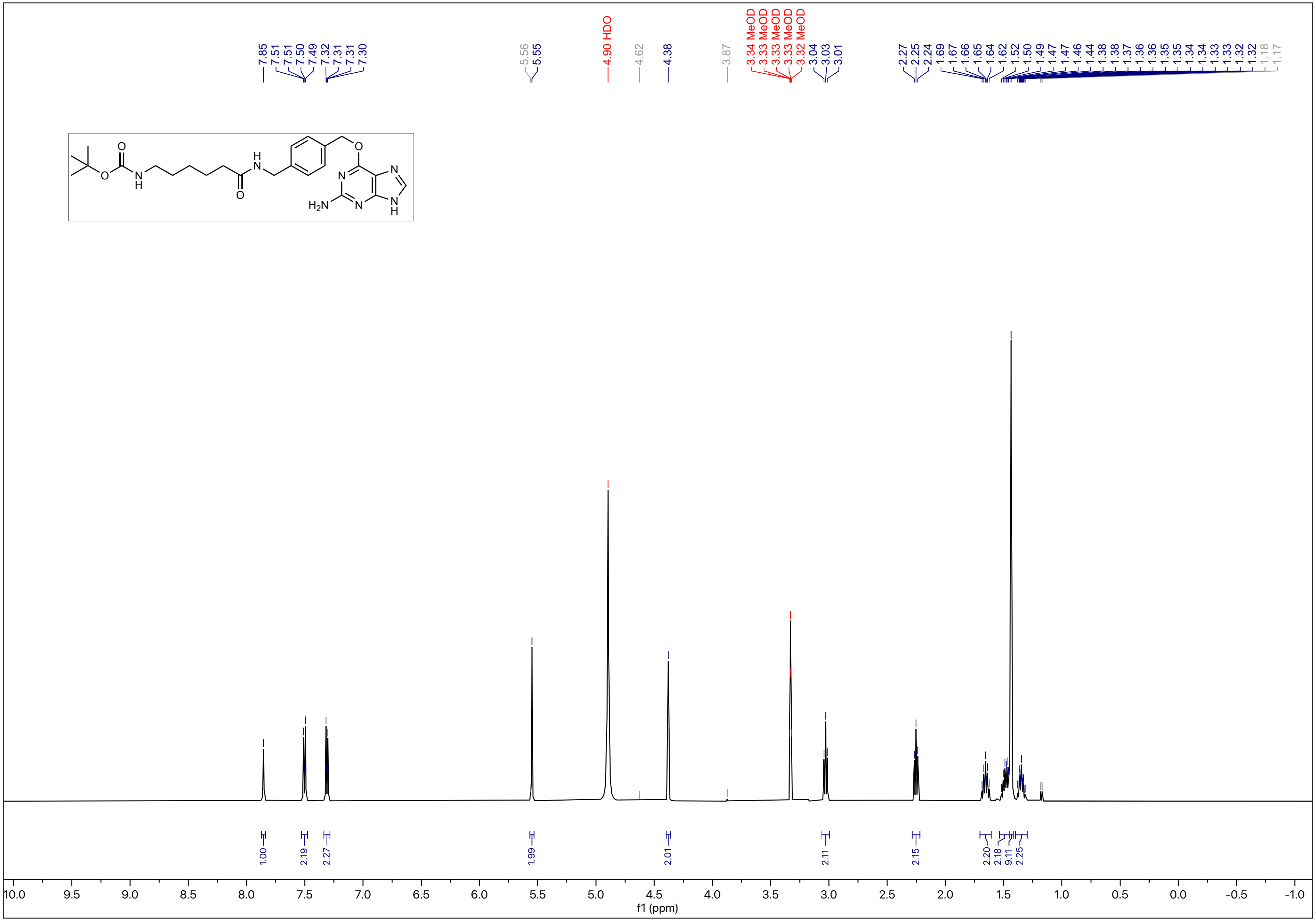

N-13

$^1H$ - NMR (500 MHz, MeOD)

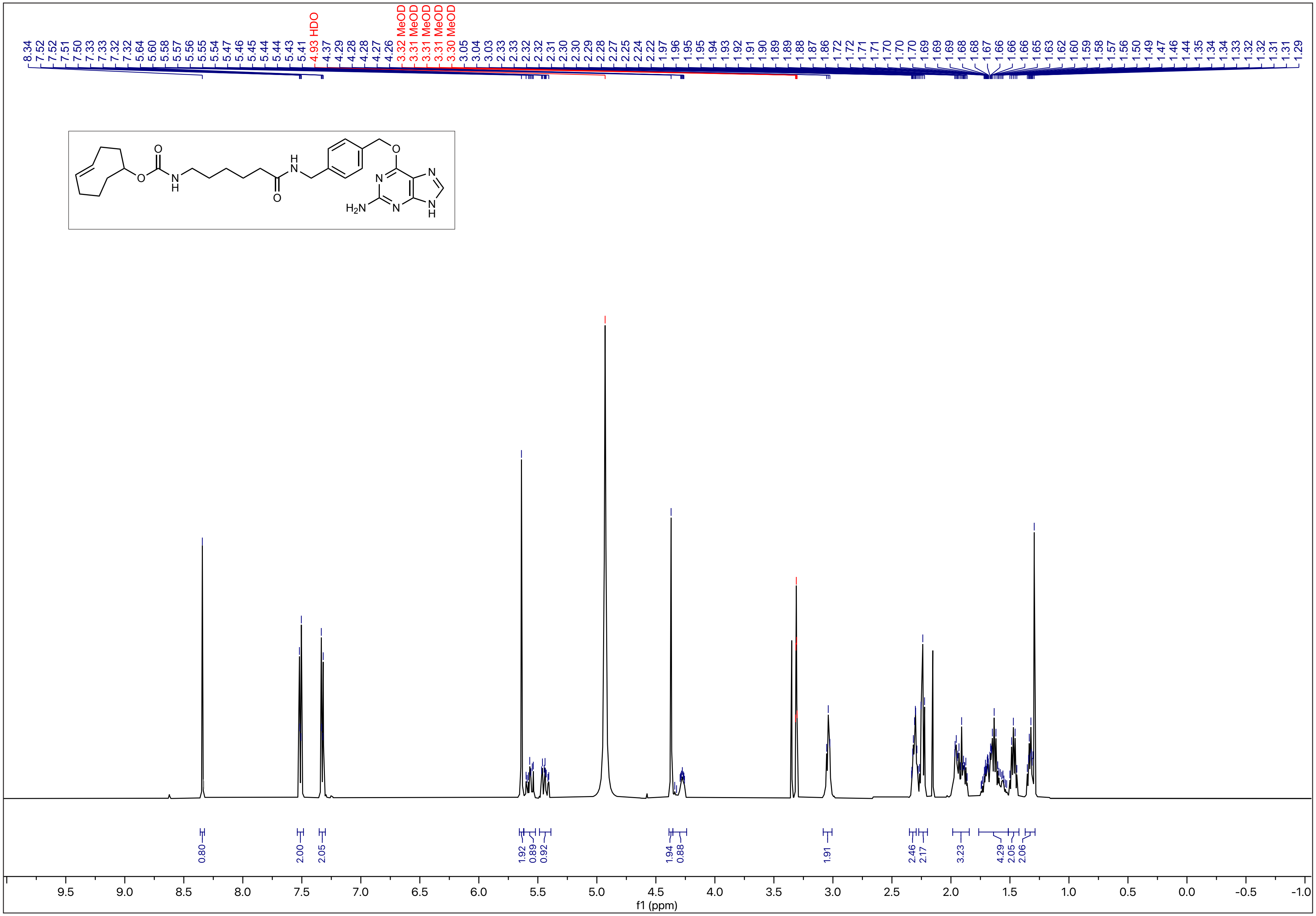

$^{13}C$ - NMR (126 MHz, MeOD)

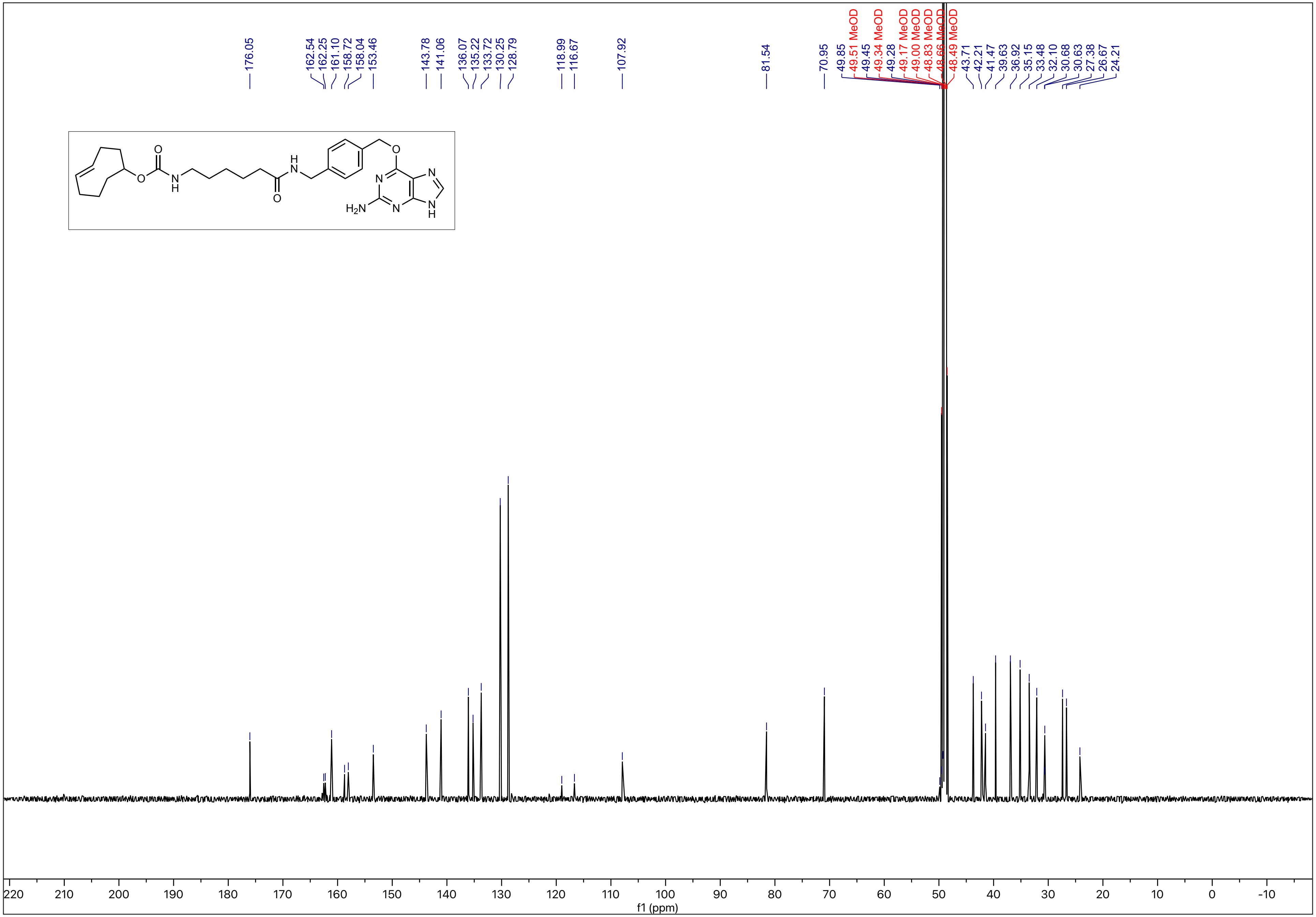

$^1H$ - NMR (500 MHz, MeOD)

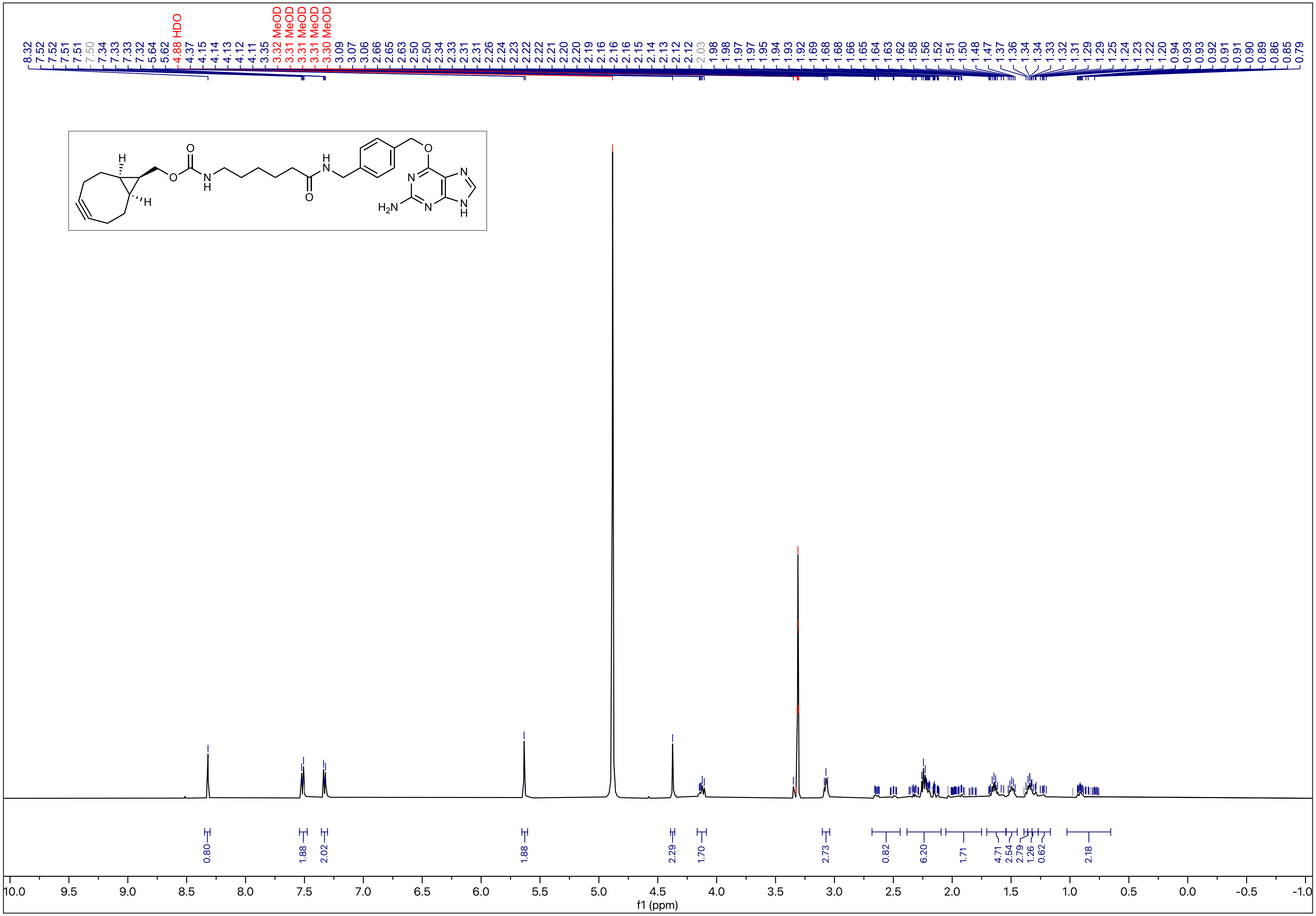

N-16

$^{13}C$ - NMR (126 MHz, MeOD)

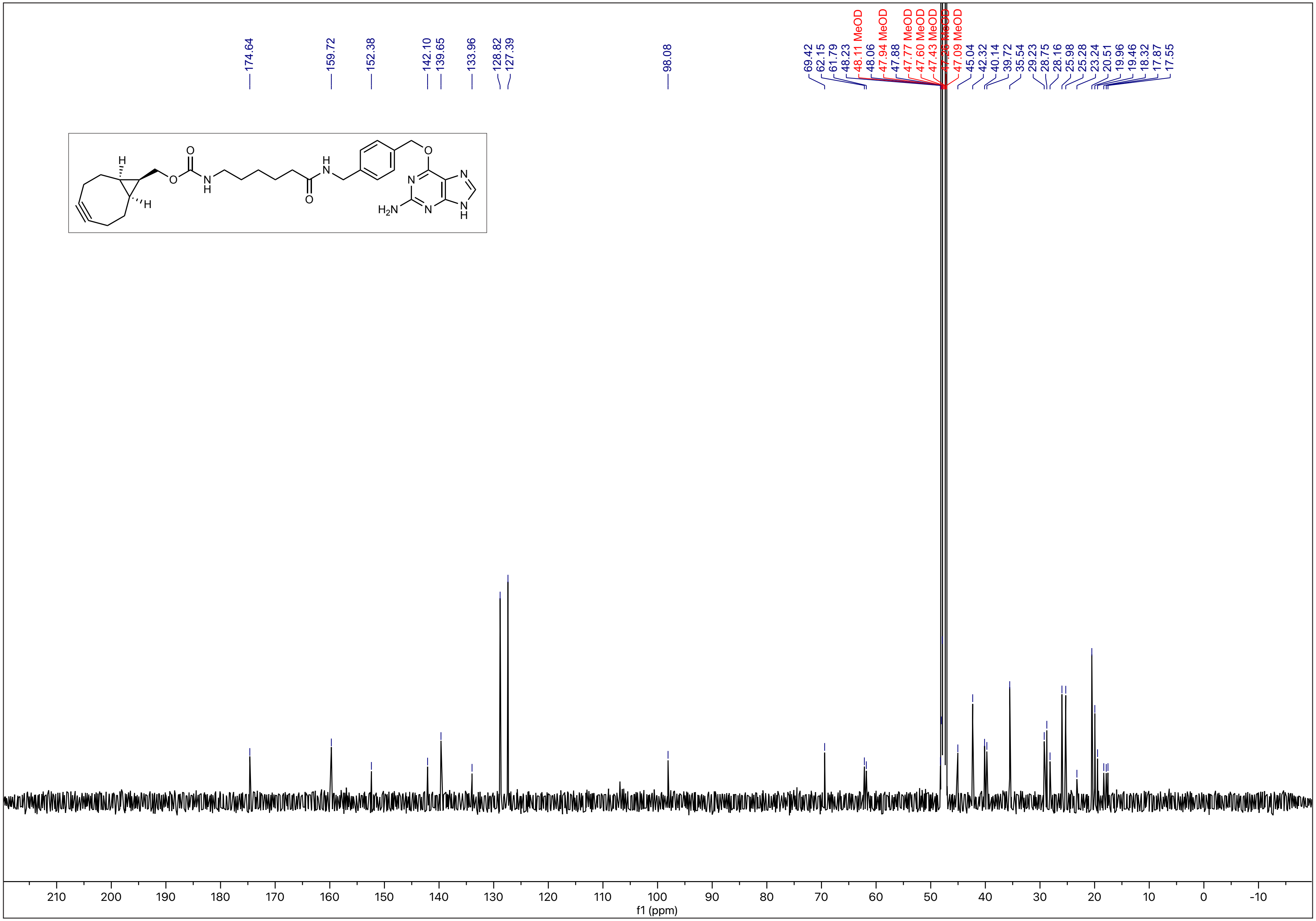

$^1H$ - NMR (600 MHz, $CDCl_3$)

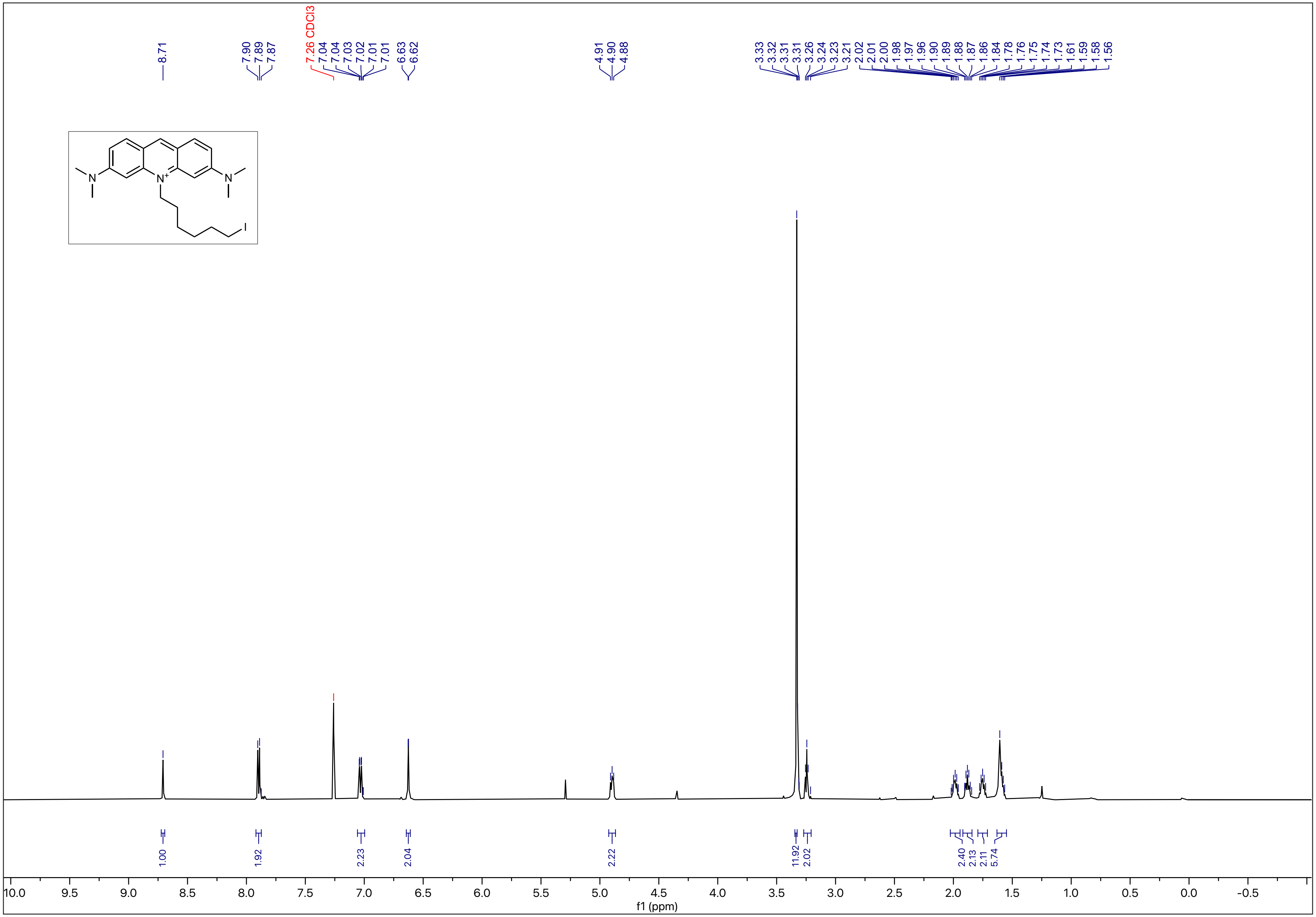

$^{13}C$ - NMR (151 MHz, $CDCl_3$)

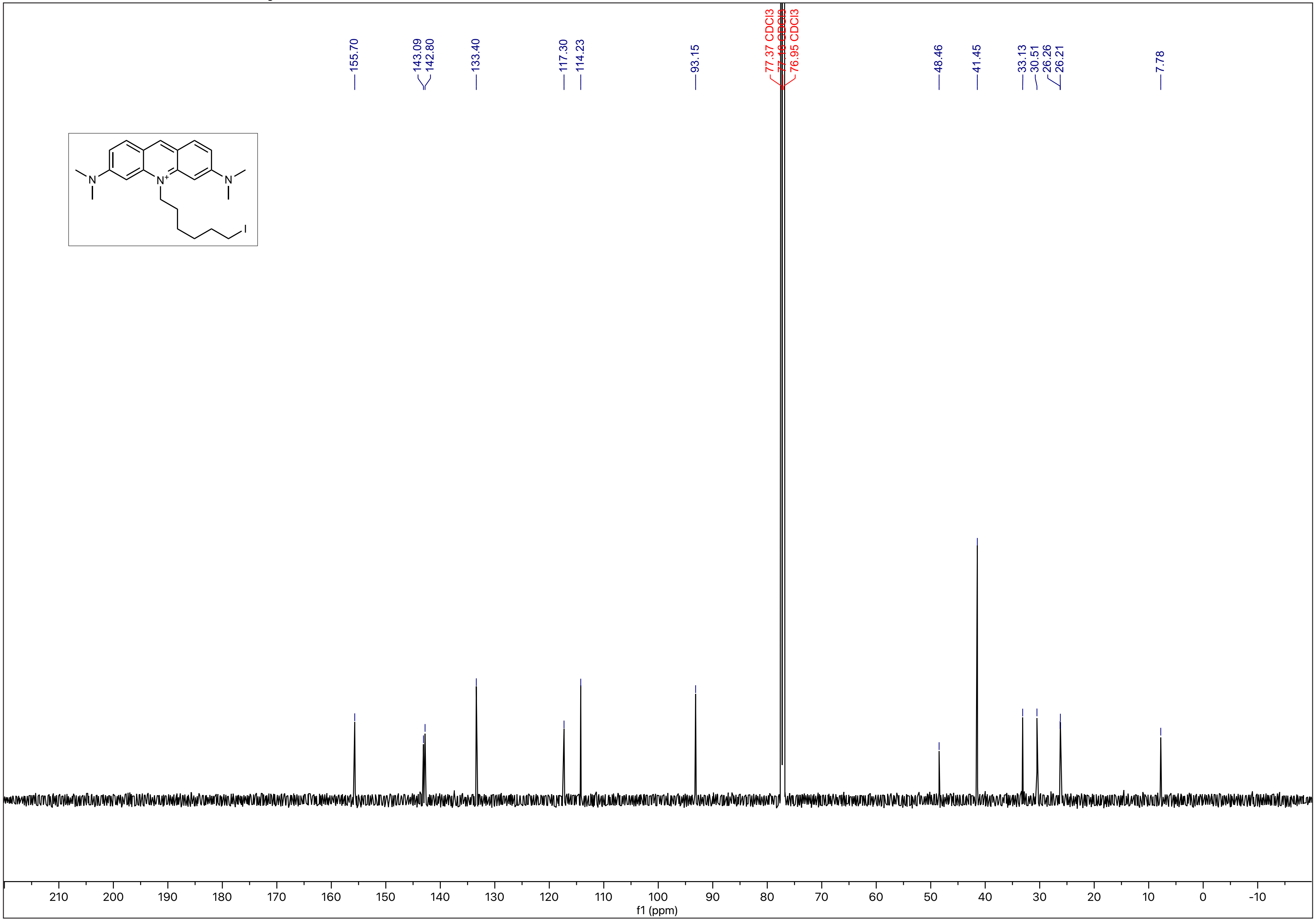

N-19

$^1$H - NMR (500 MHz, $CDCl_3$)

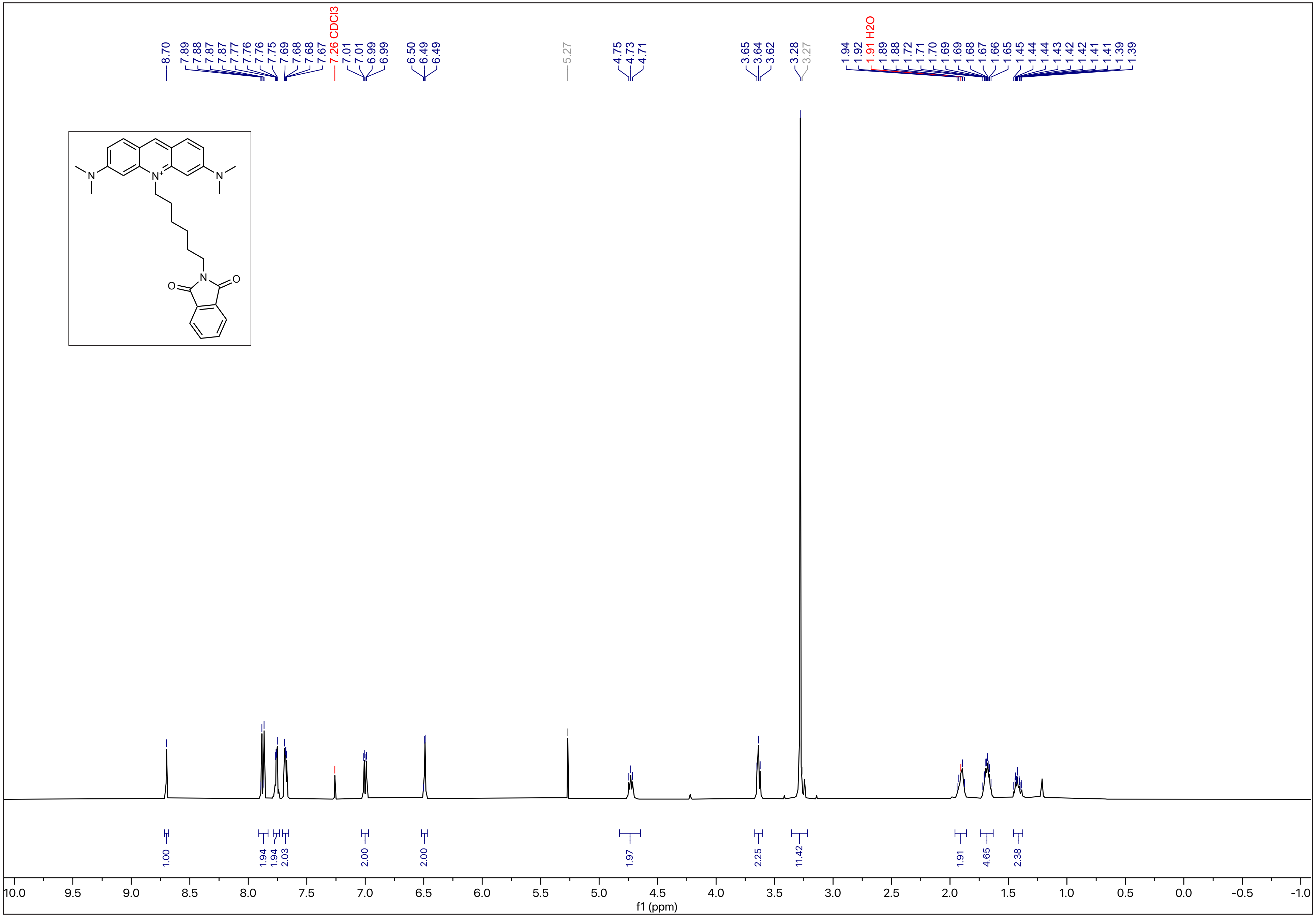

N-20

$^{13}C$ - NMR (126 MHz, $CDCl_3$)

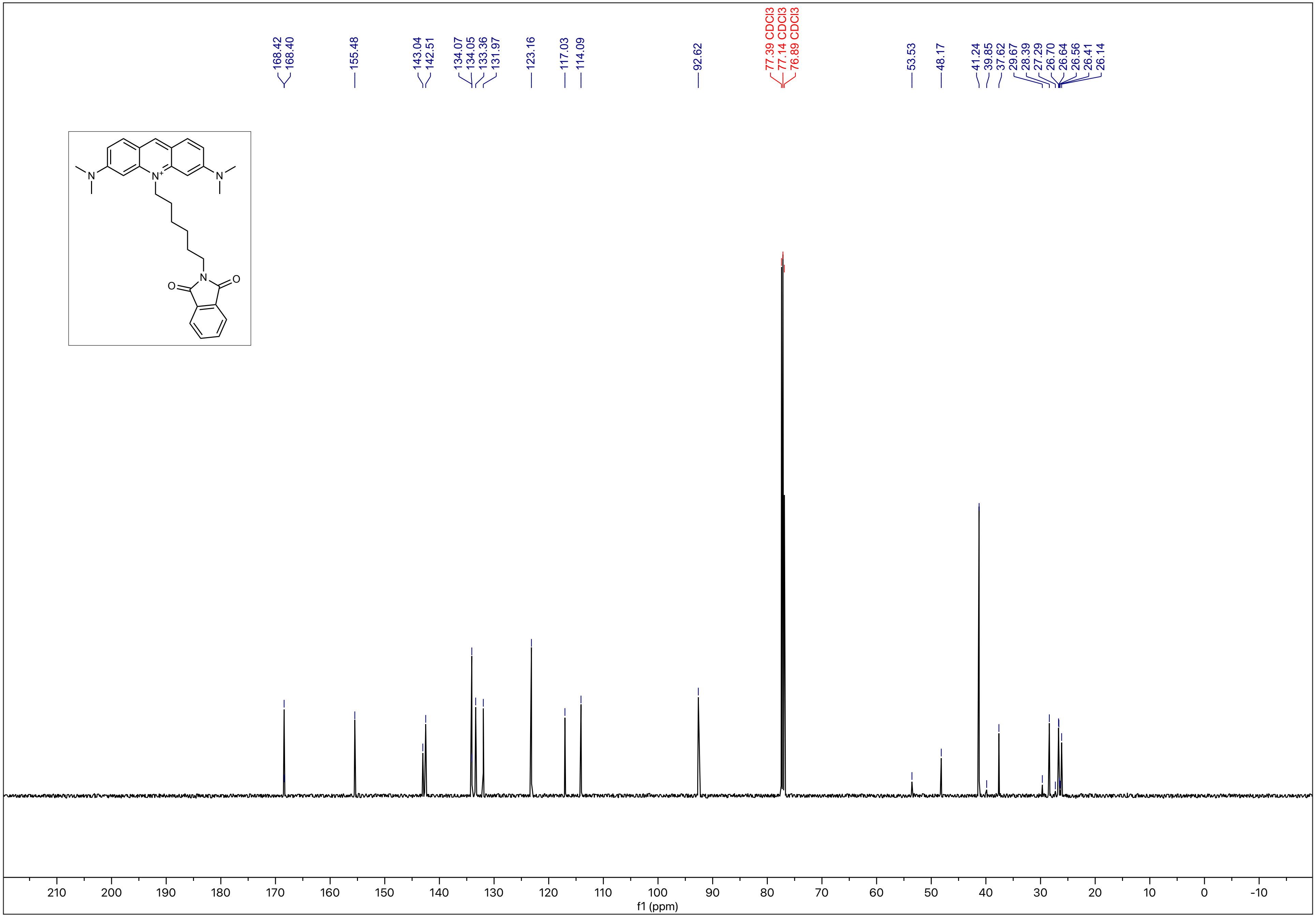

$^{1}H$ - NMR (600 MHz, MeOD)

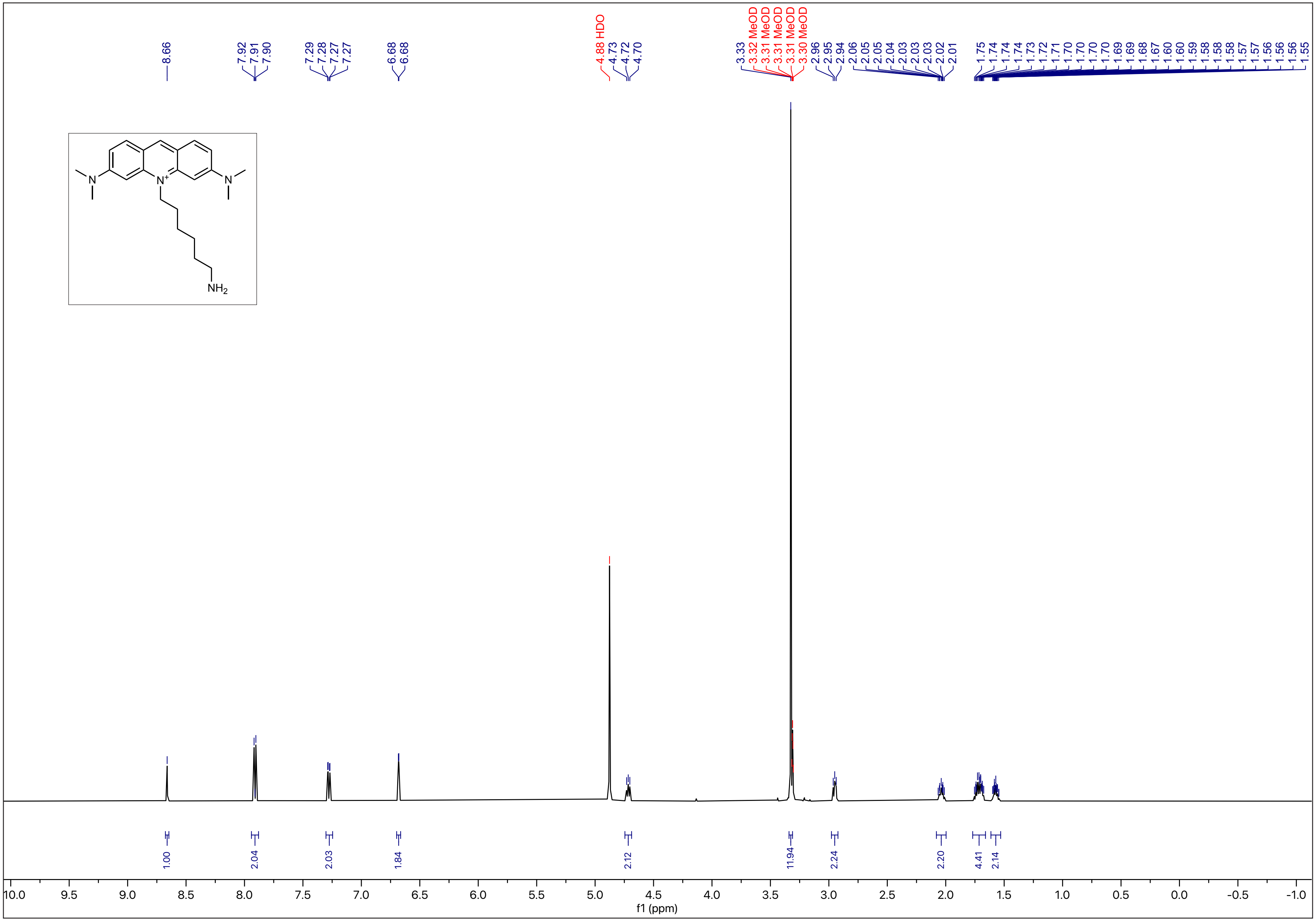

N-22

$^{13}C$ - NMR (151 MHz, MeOD)

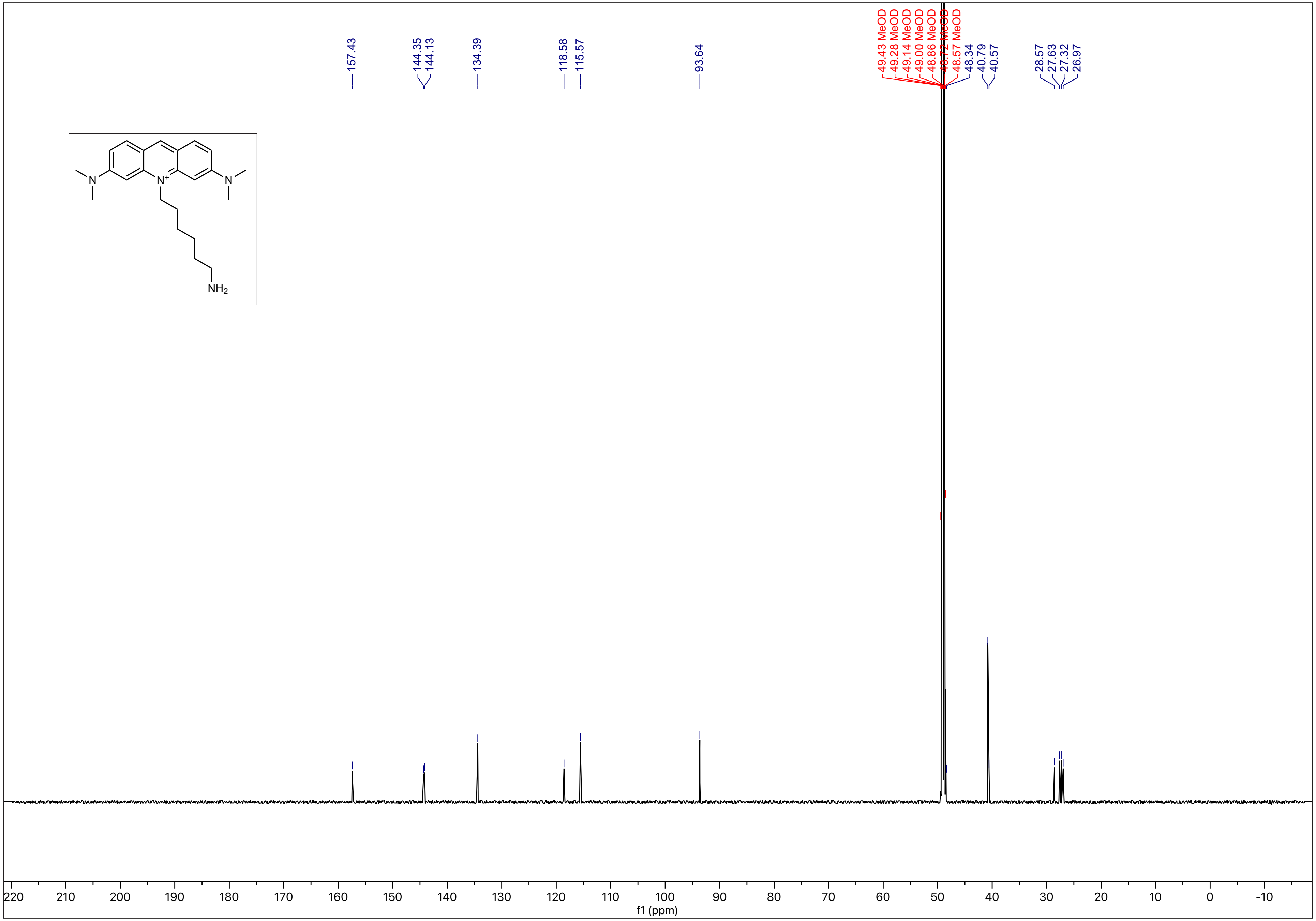

N-23

$^1$H - NMR (600 MHz, MeOD)

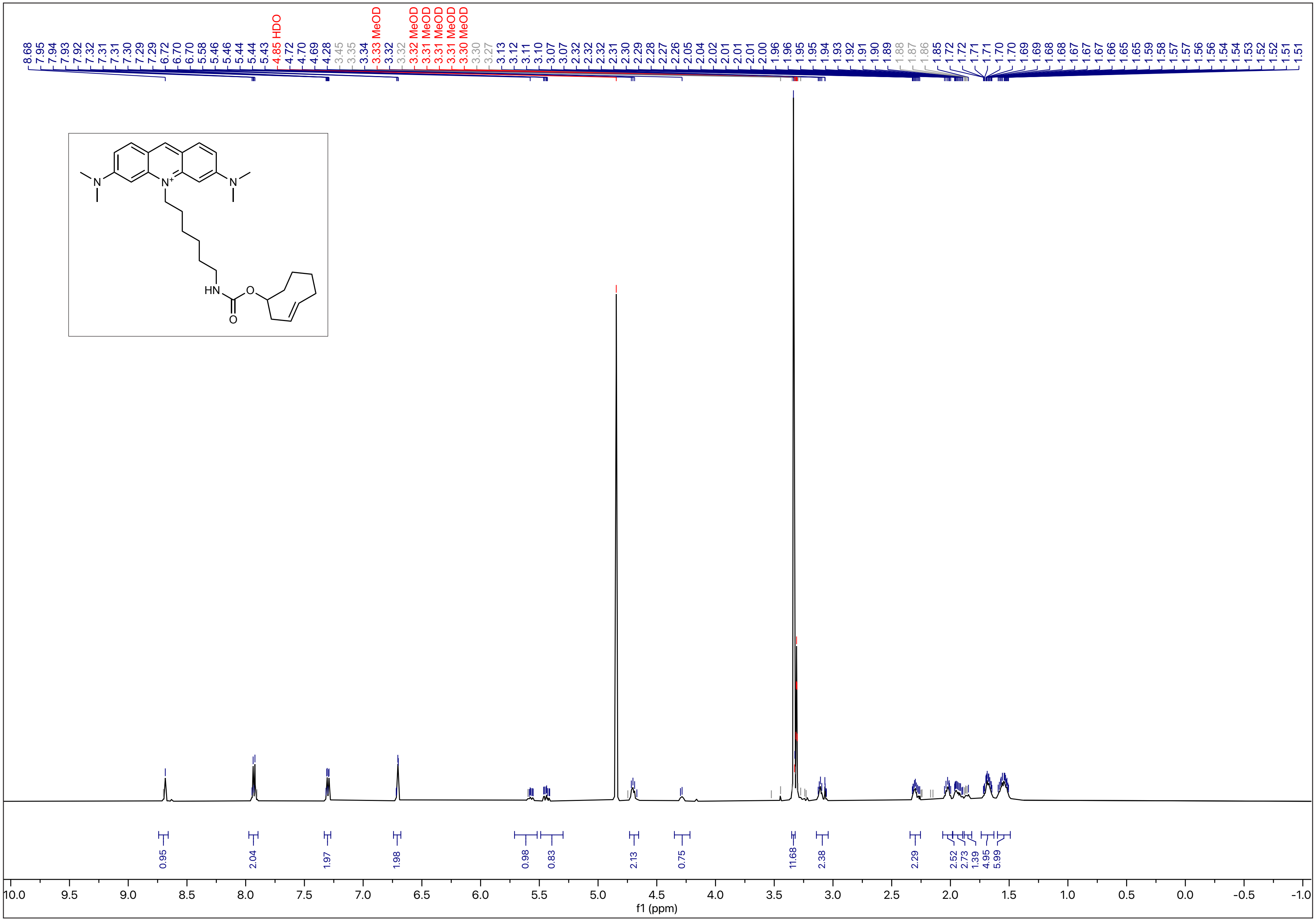

N-24

$^{13}C$ - NMR (151 MHz, MeOD)

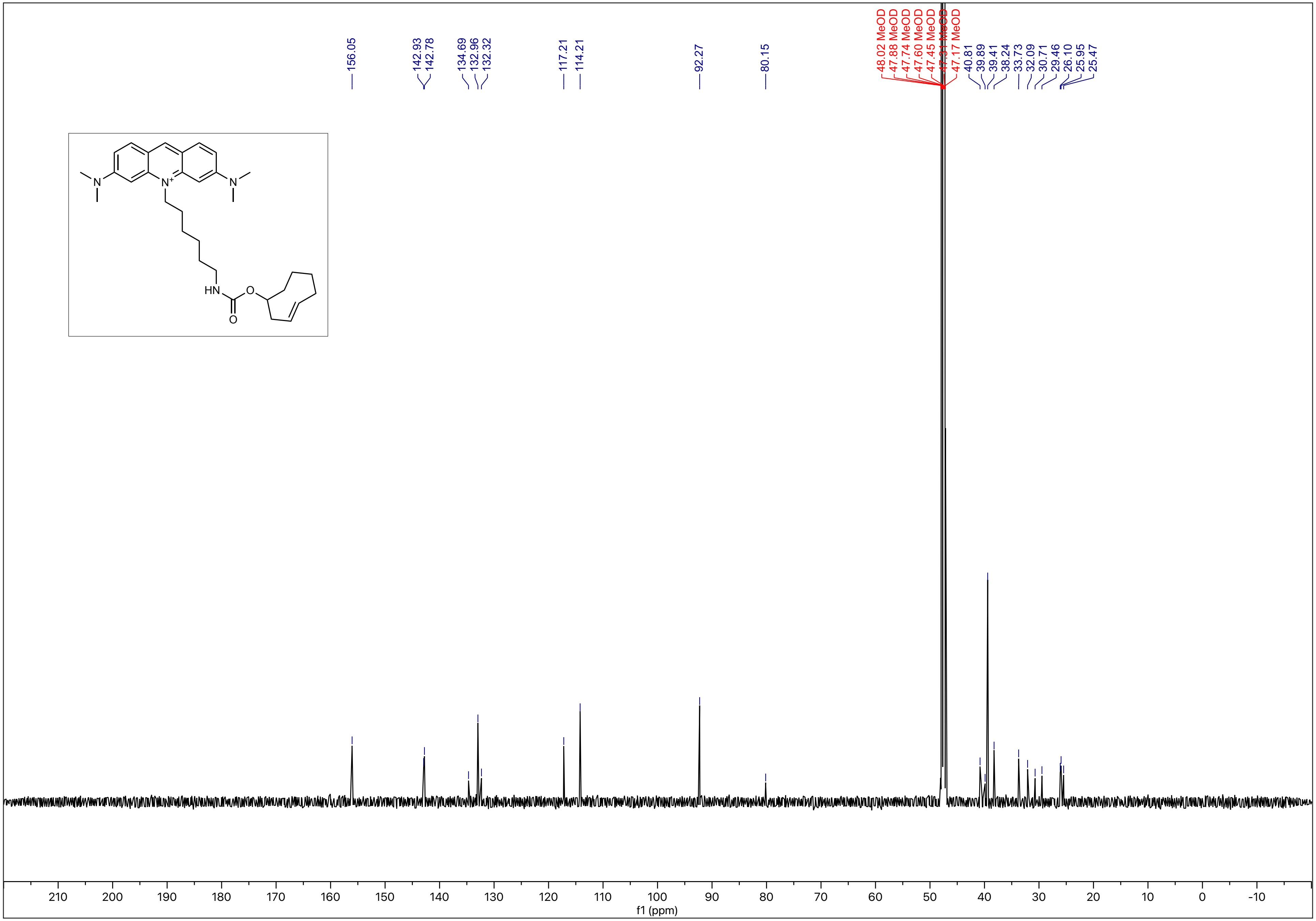

[1]H - NMR (600 MHz, MeOD)

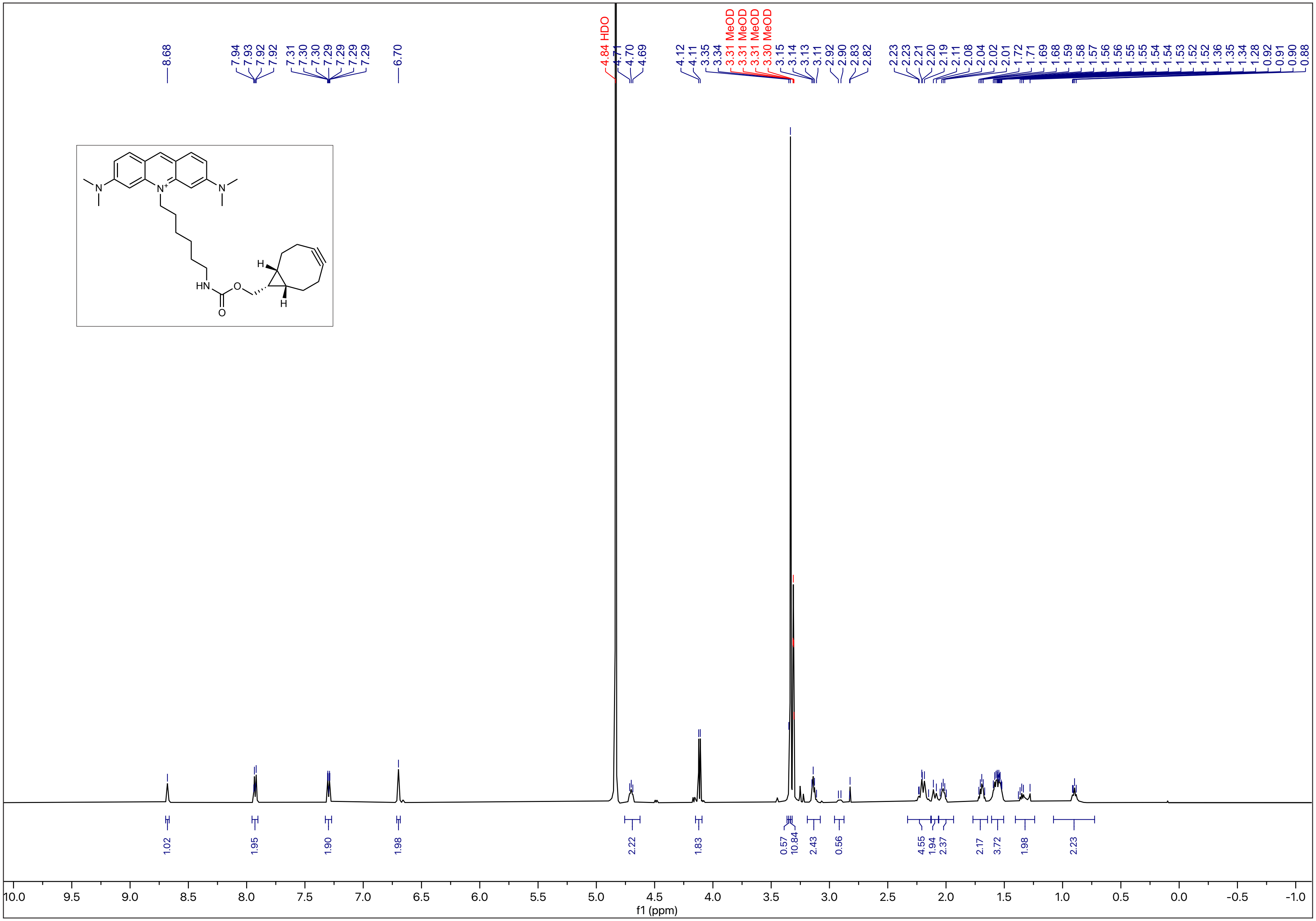

$^{13}C$ - NMR (151 MHz, MeOD)

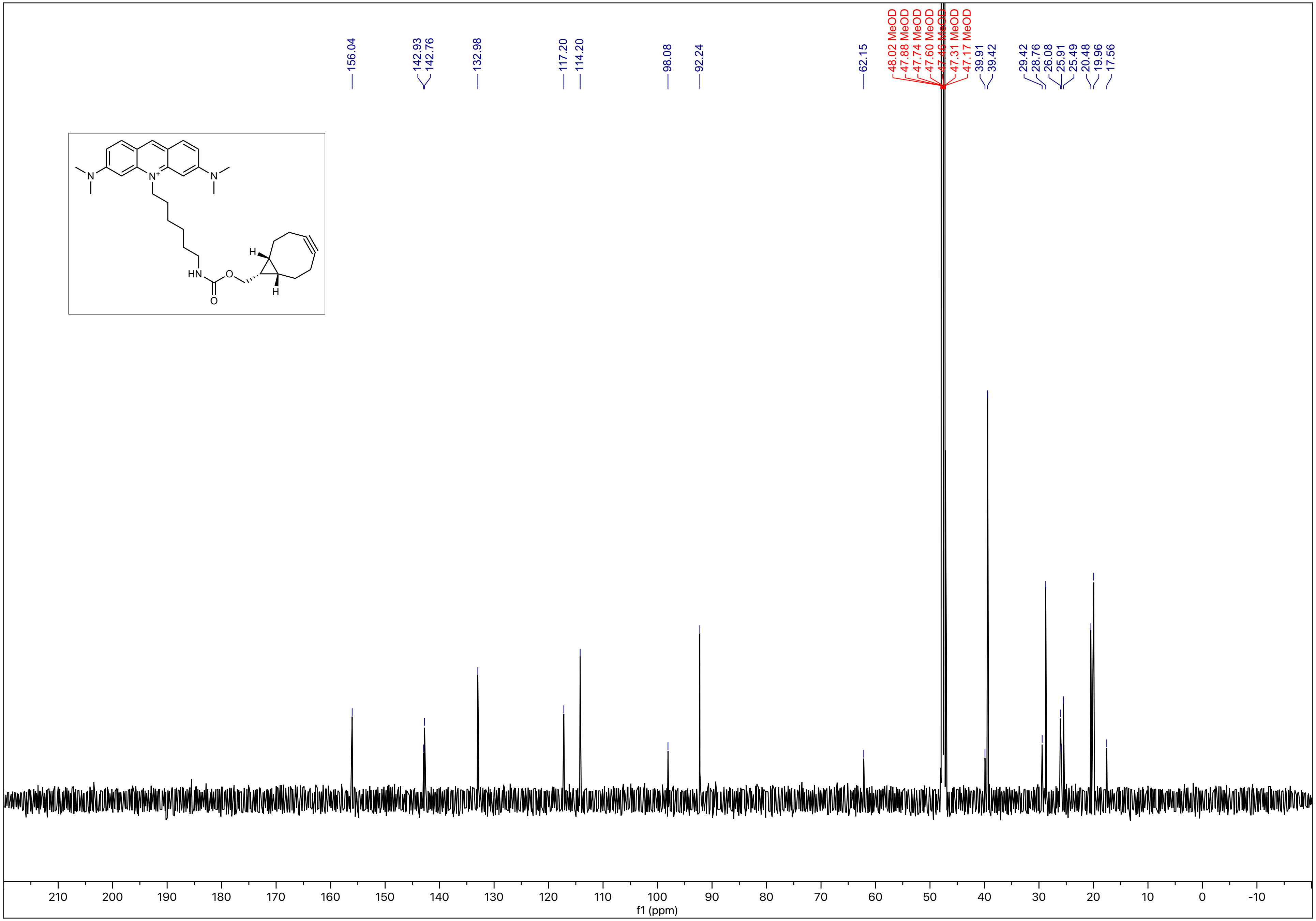

[1]H - NMR (500 MHz, MeOD)

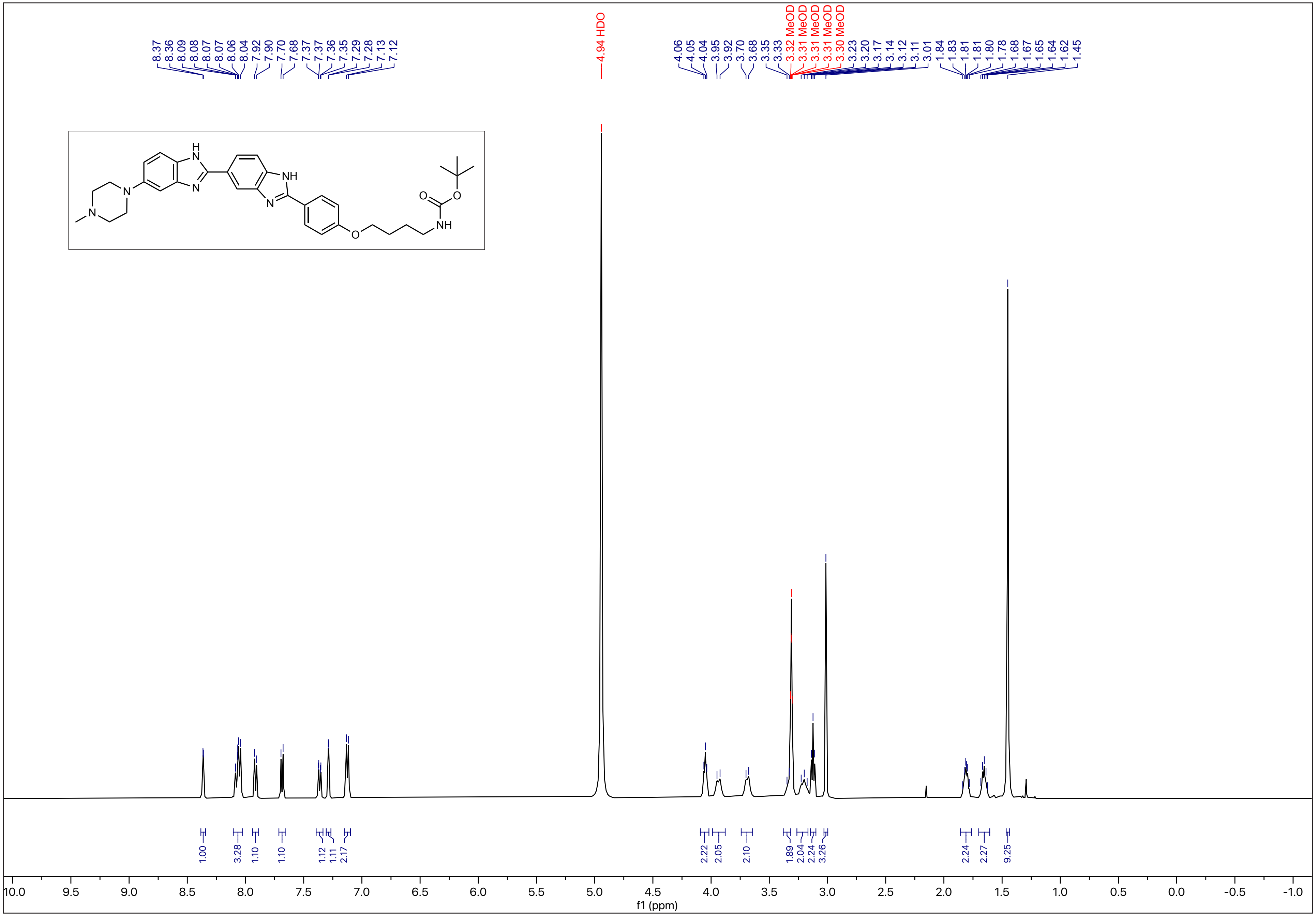

N-28

$^{13}C$ - NMR (126 MHz, MeOD)

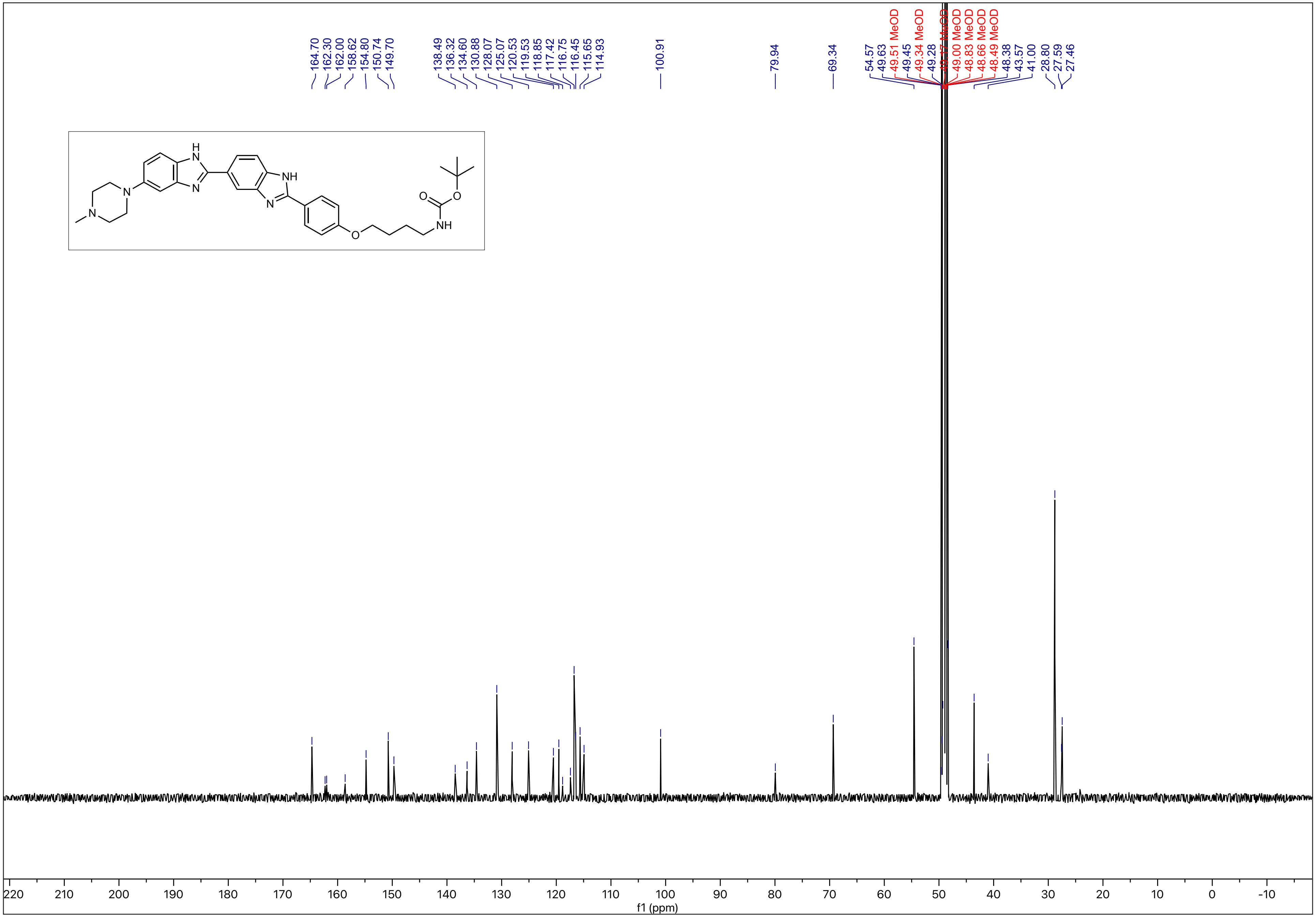

[1]H - NMR (600 MHz, MeOD)

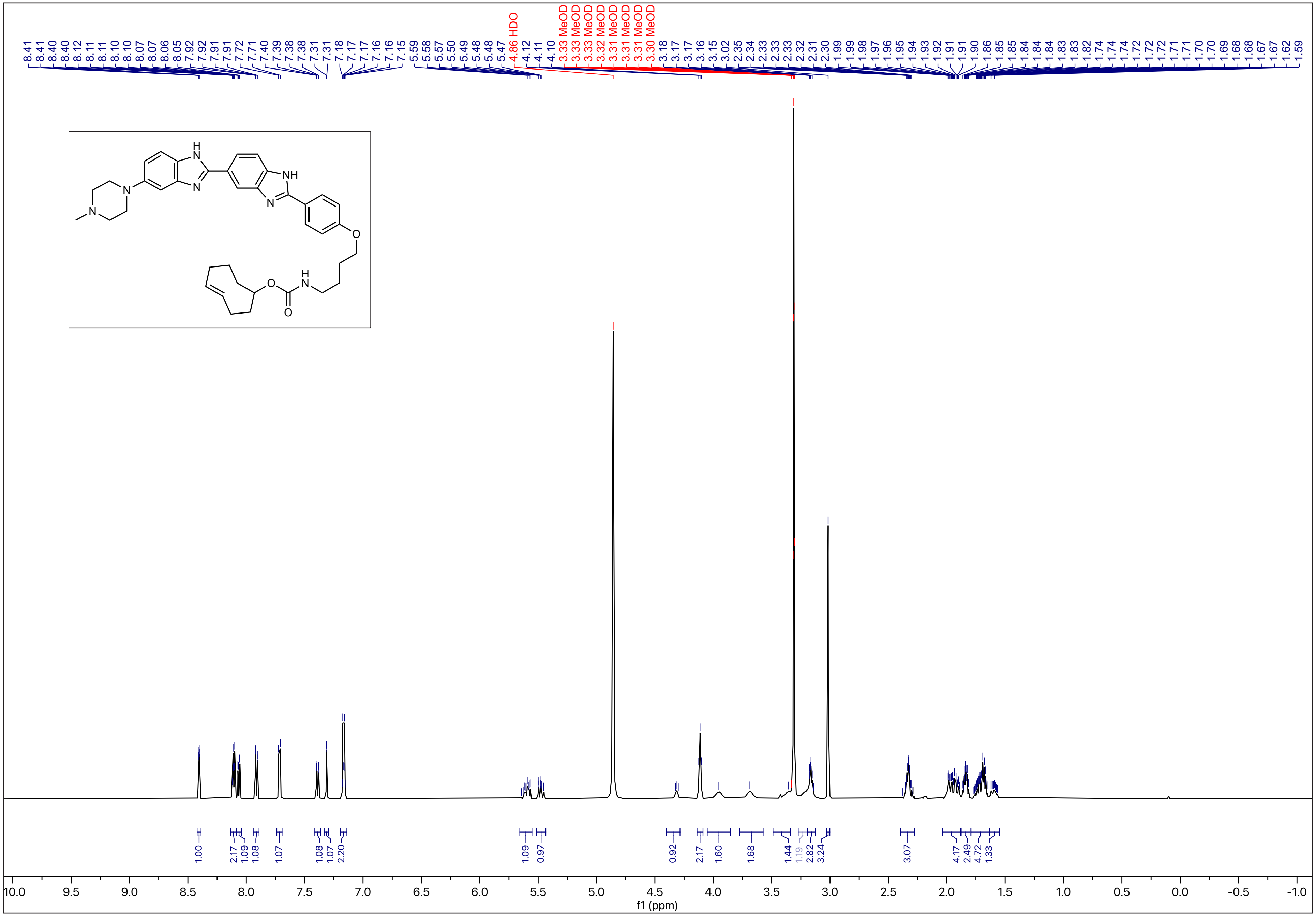

$^{13}$C - NMR (151 MHz, MeOD)

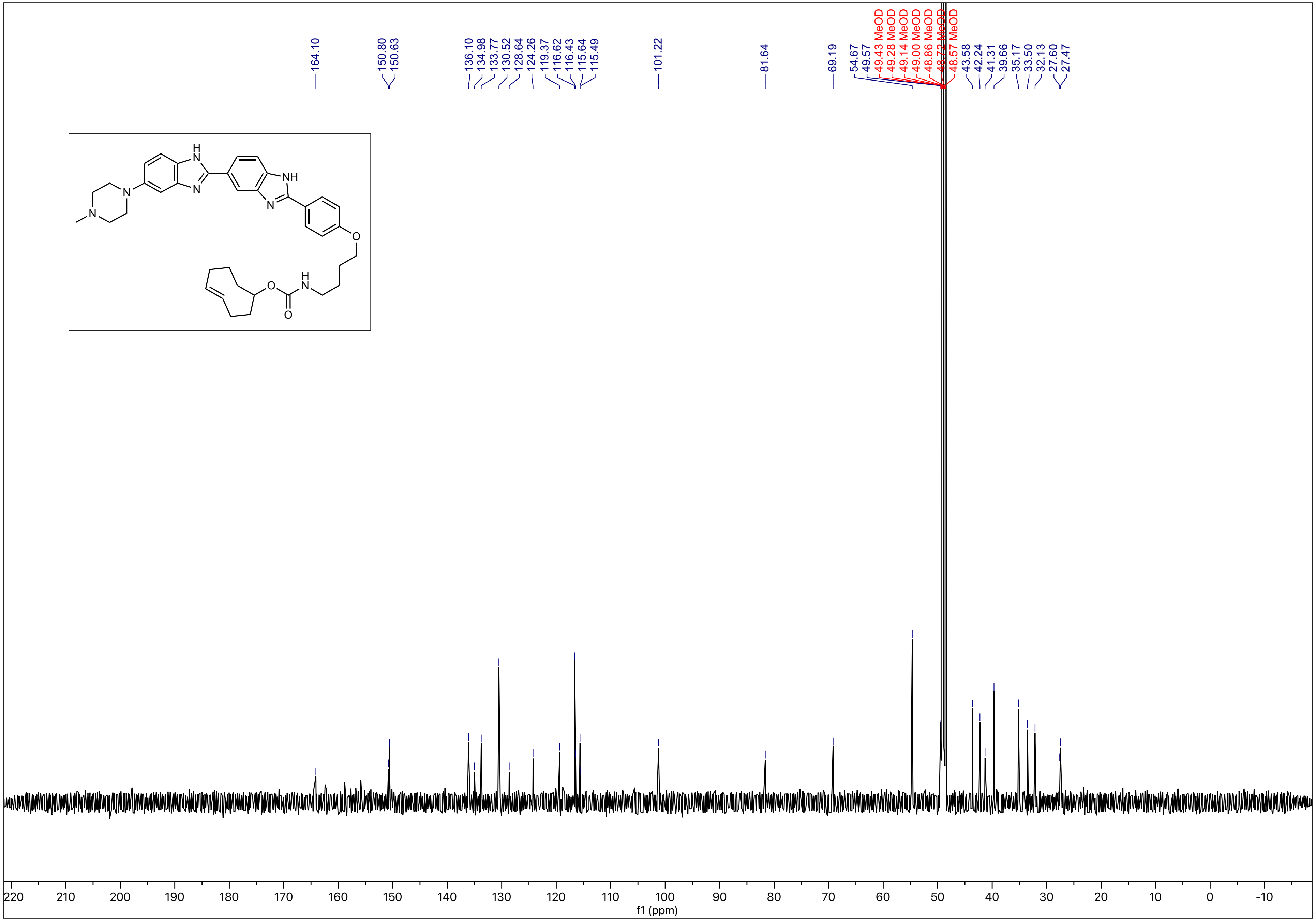

N-31

[1]H - NMR (600 MHz, MeOD)

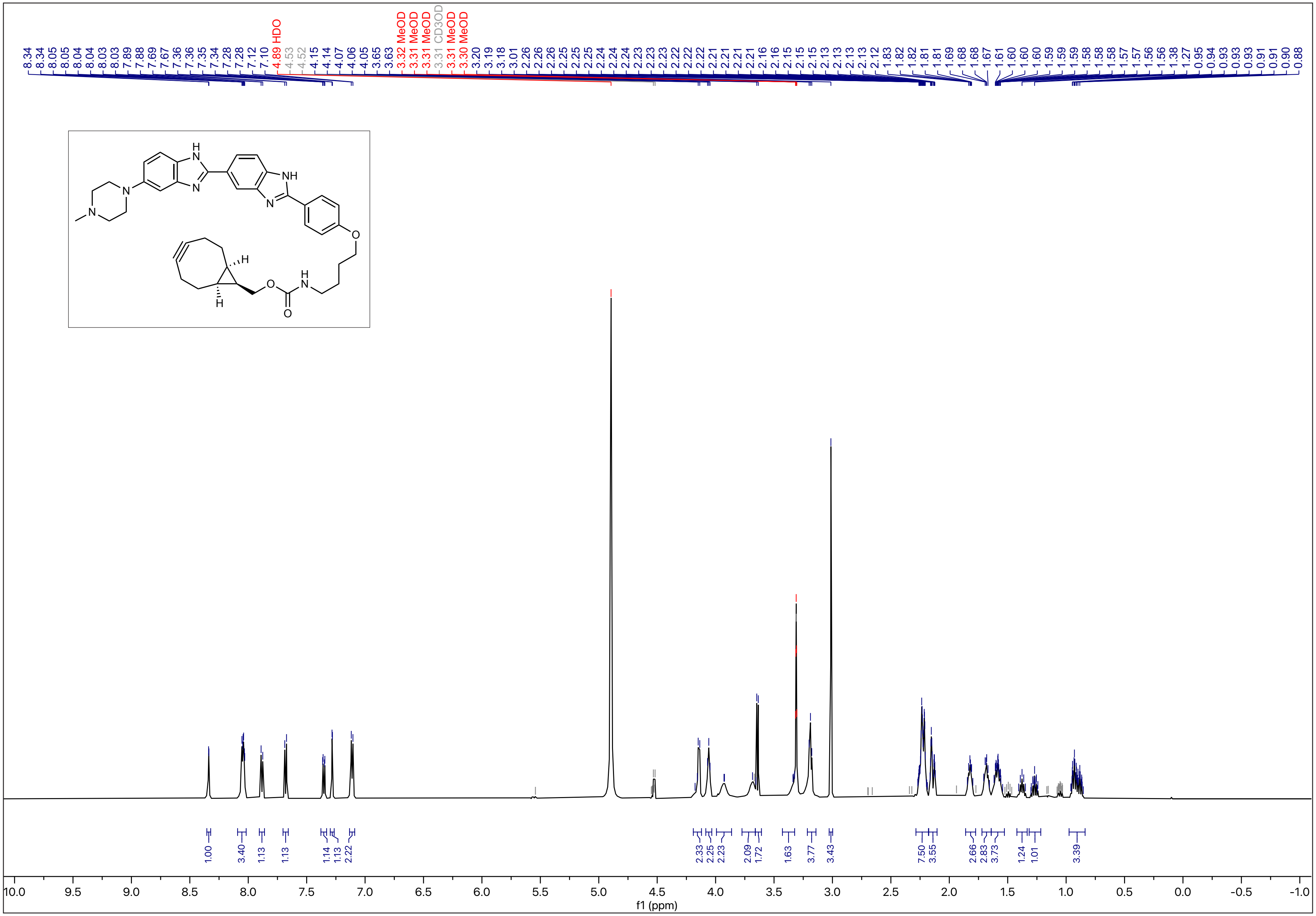

$^{13}C$ - NMR (151 MHz, MeOD)

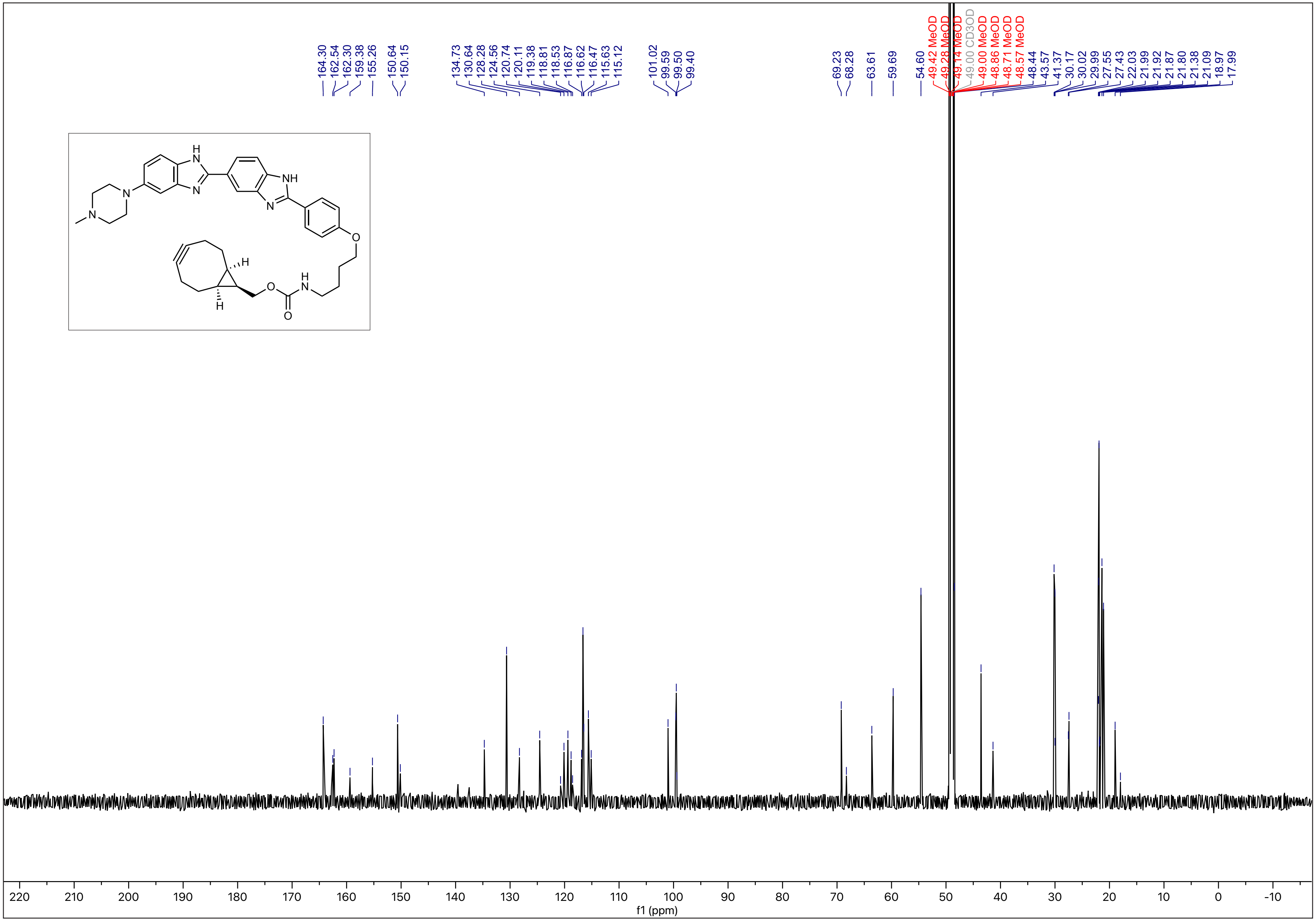

[1]H - NMR (500 MHz, MeOD)

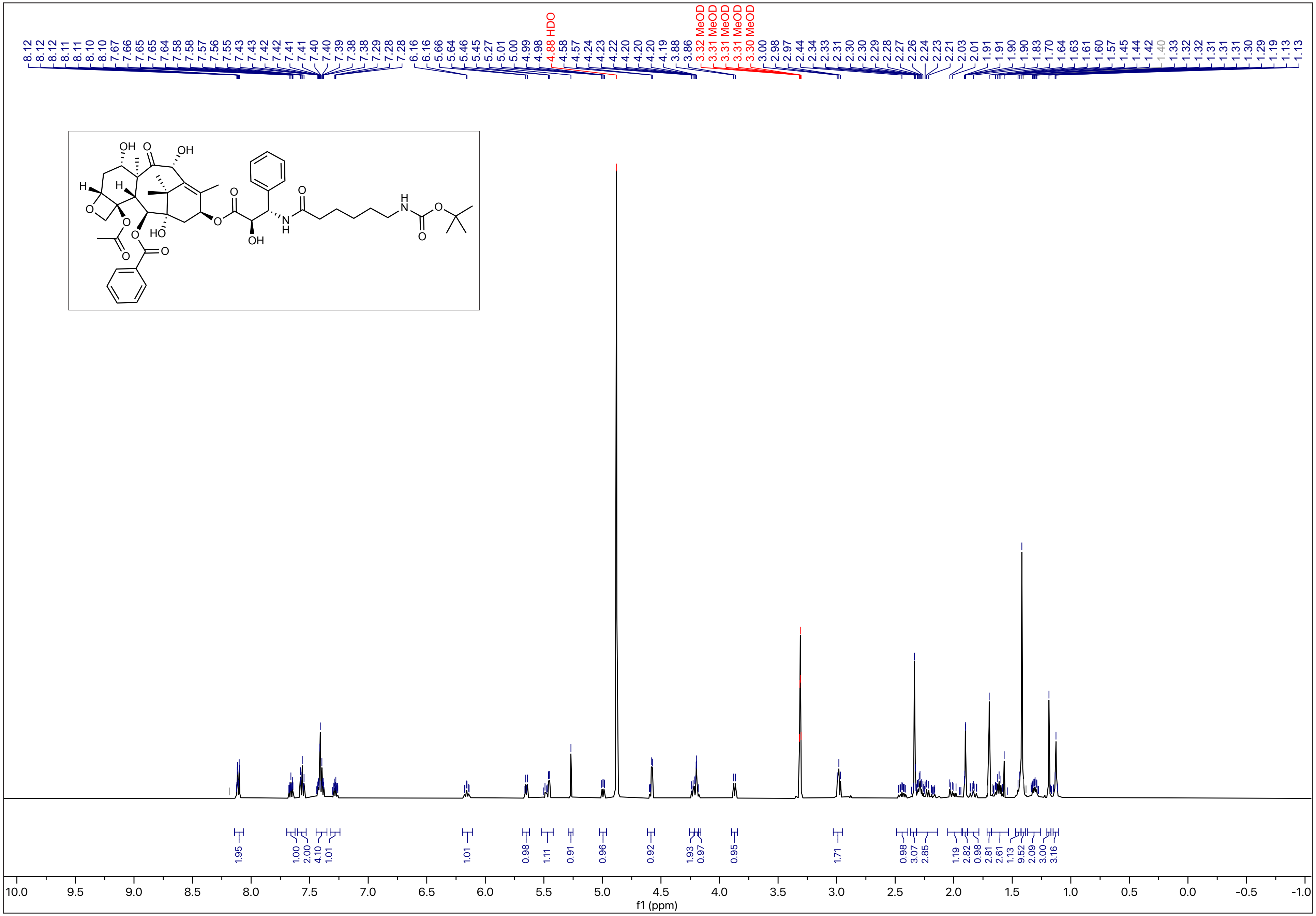

[13]C - NMR (126 MHz, MeOD)

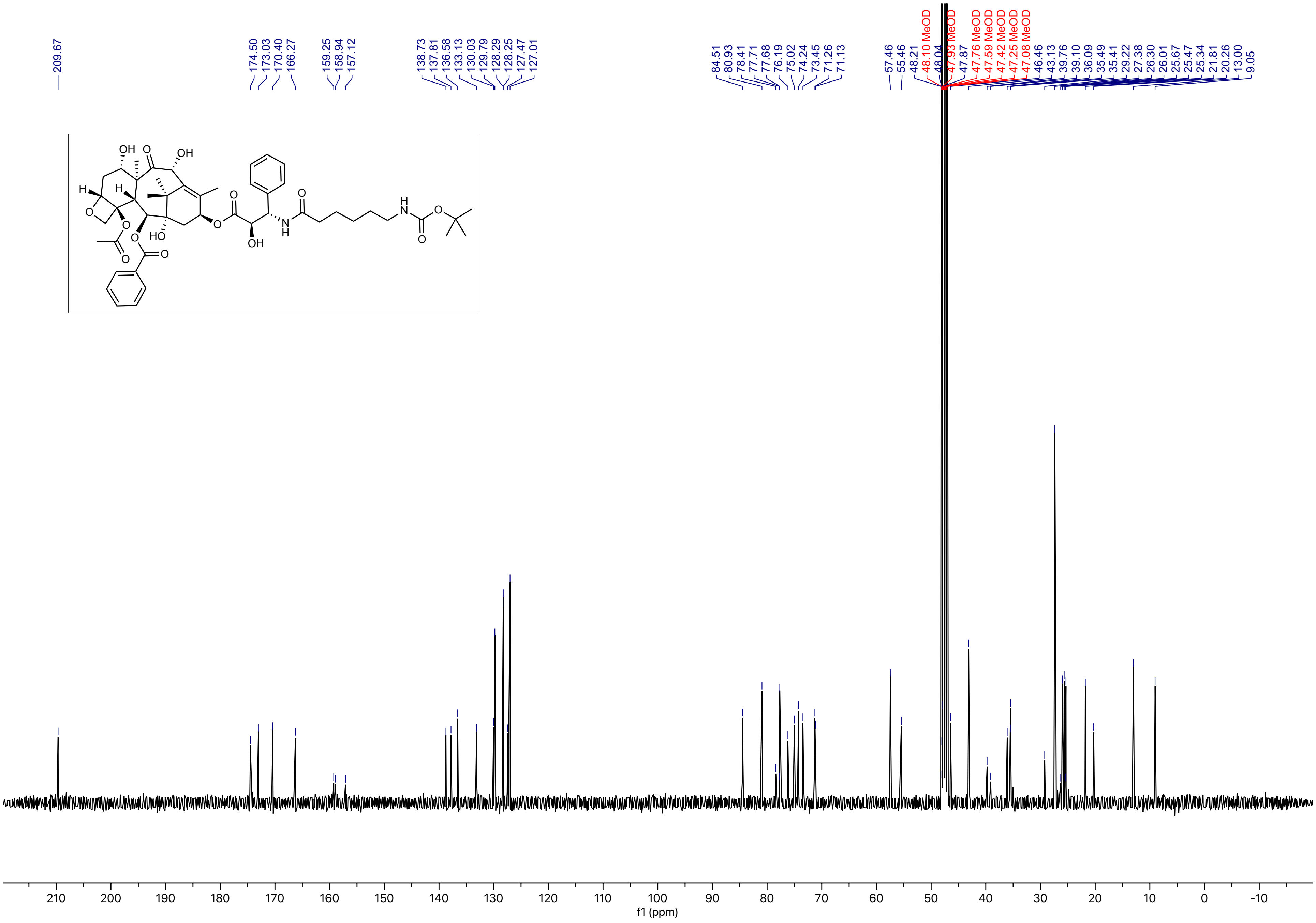

N-35

[1]H - NMR (600 MHz, MeOD)

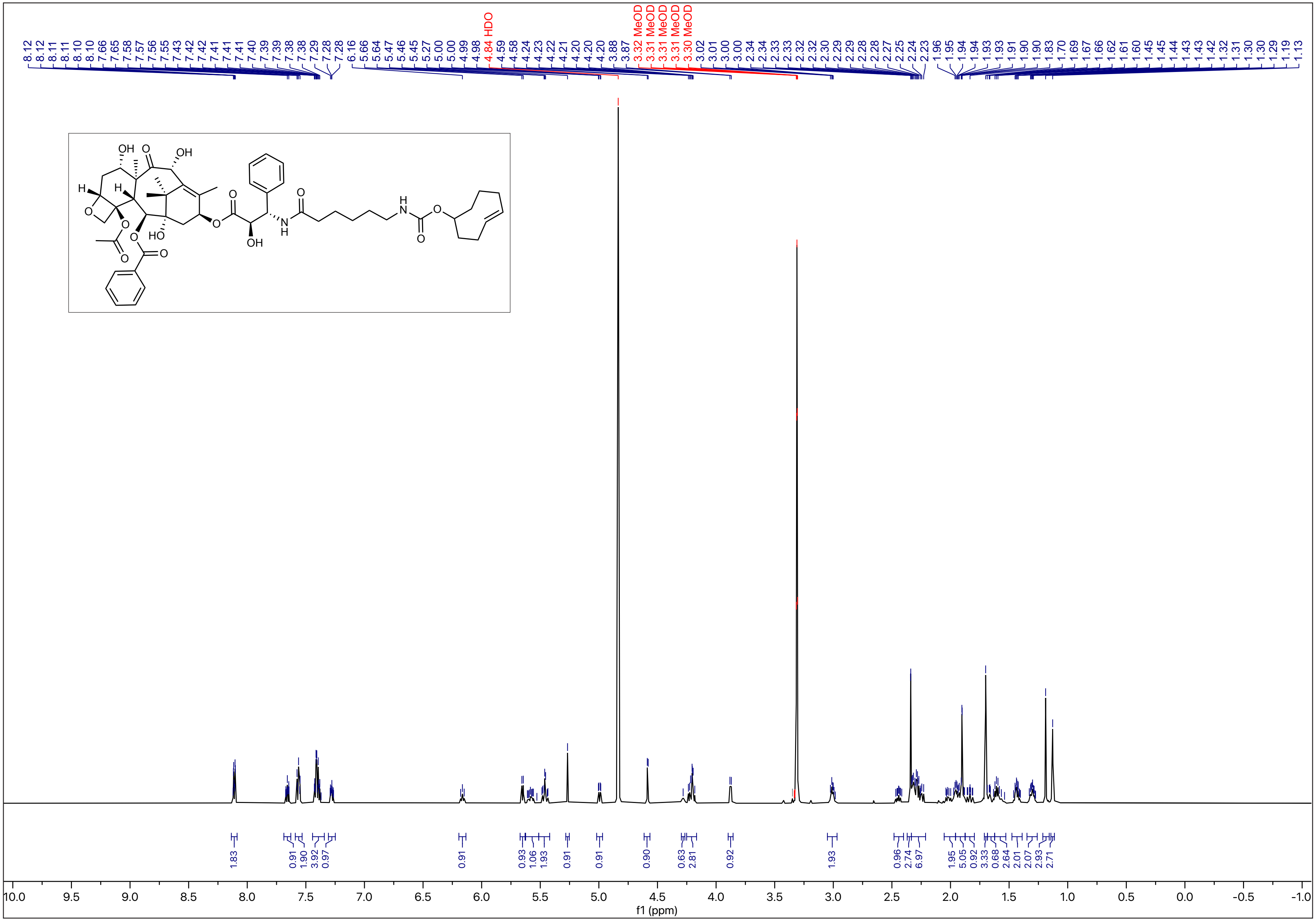

$^{13}$C - NMR (151 MHz, MeOD)

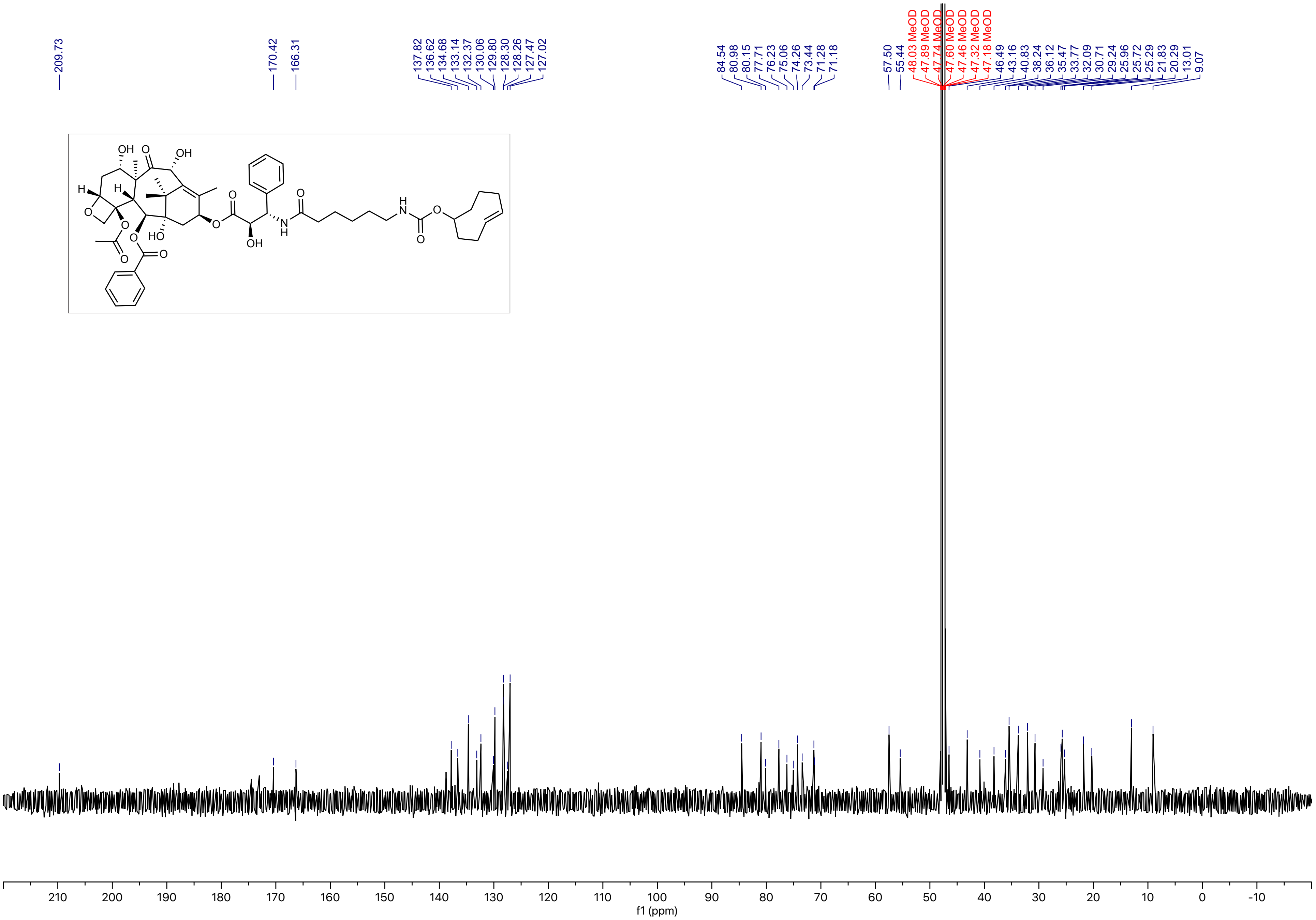

[1]H - NMR (600 MHz, MeOD)

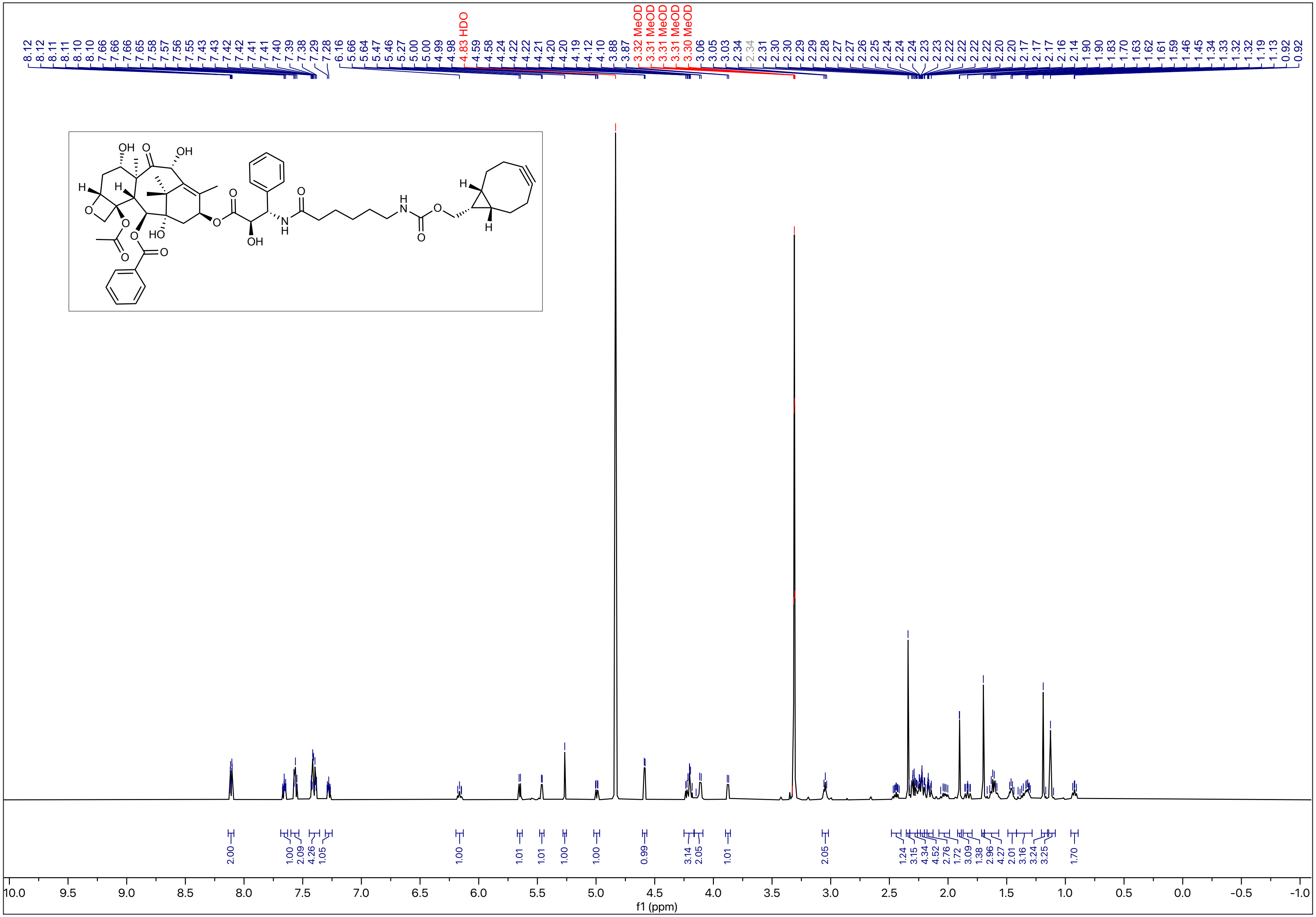

$^{13}C$ - NMR (151 MHz, MeOD)

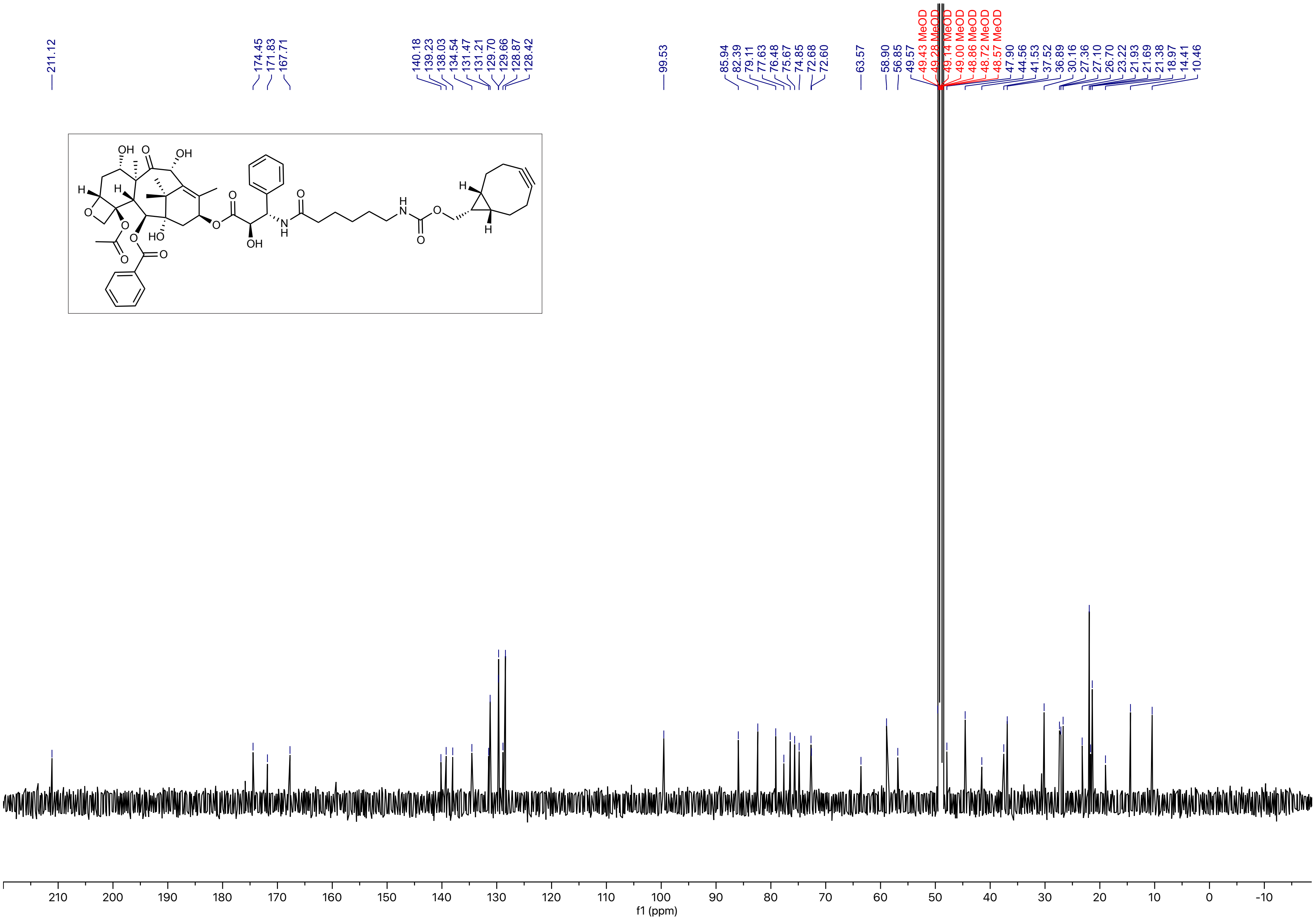

$^{1}H$ - NMR (400 MHz, MeOD)

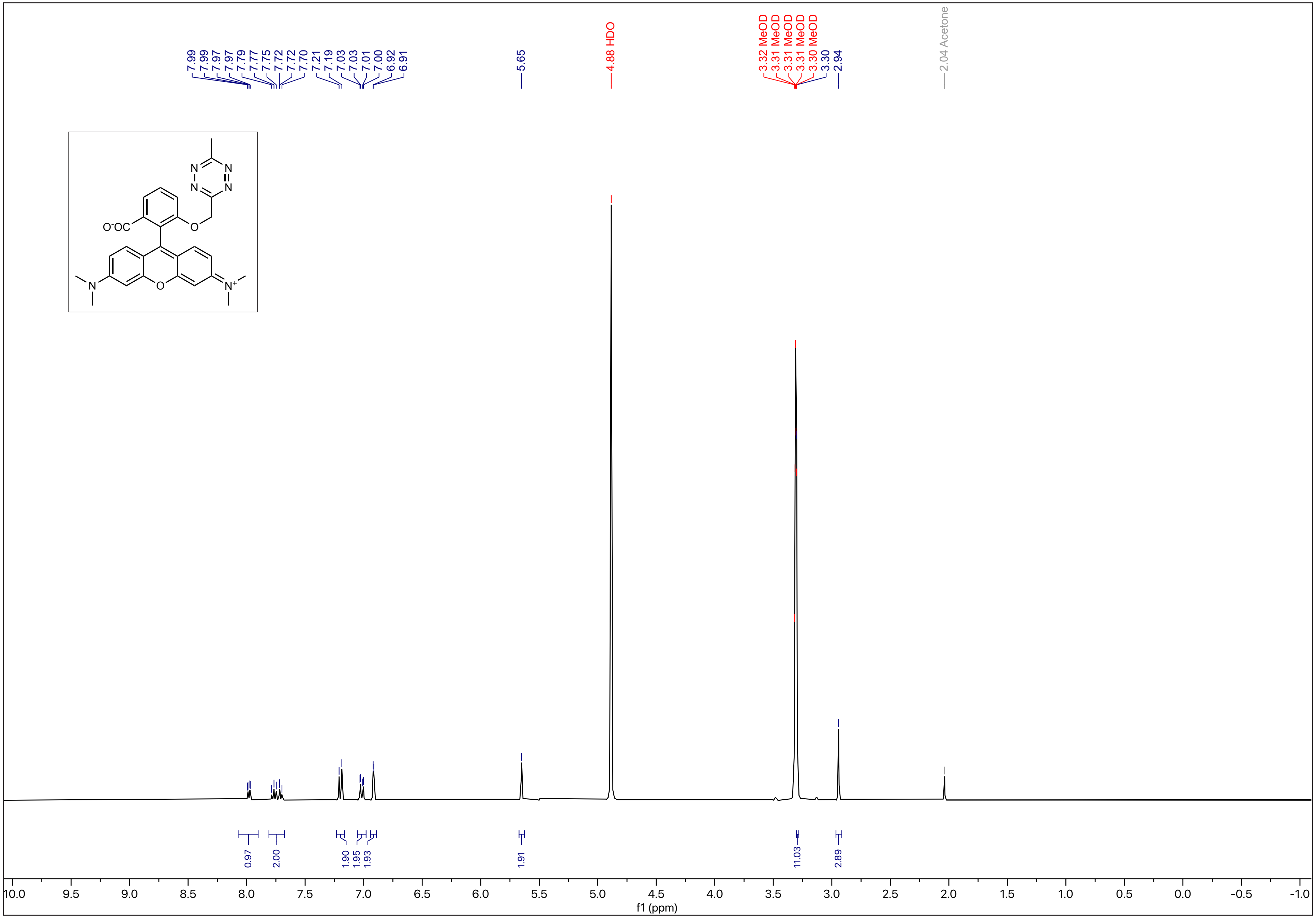
